# Stopover departure decisions and movement patterns of migratory bats during autumn migration

**DOI:** 10.64898/2026.04.29.721682

**Authors:** Sander Lagerveld, Julia Karagicheva, Pepijn de Vries, Eldar Rakhimberdiev, Karina Stienstra, Bart C.A. Noort, Martin J.M. Poot, Thiemo Karwinkel, Georg Rüppel, Vera Brust, Fiona Mathews, Heiko Schmaljohann, Frank van Langevelde

## Abstract

Migrating bats alternate between stopover periods and directed flights. When departing from a stopover site, bats select the night, the specific time within the night, and the flight direction to resume migration. Despite their ecological importance, the factors shaping these stopover departure decisions remain poorly understood. To identify the intrinsic and environmental factors driving departure decisions and movement patterns, we tagged *Nathusius’ pipistrelles*

*Pipistrellus nathusii* at three coastal locations in the Netherlands and tracked 178 individuals during autumn migration, using the MOTUS Wildlife Tracking System. We examined movement patterns and analysed departure probability in relation to a set of individual and environmental covariates in a Bayesian capture-recapture model in state-space formulation. Additionally, we modelled within-night variation in departure timing. Seasonal patterns were strongly influenced by reproductive behaviour, with decreased migration probability during the mating period. Regardless of their seasonal timing, bats departed under moderate tailwinds and dry conditions, optimizing energy efficiency, while avoiding crosswinds and cloud cover, enhancing navigational safety. Most individuals departed shortly after sunset, whereas headwinds delayed nocturnal departure. Movement patterns were diverse, including migration towards lower latitudes, coastal barrier movements, and long-distance roundtrips, suggesting the use of multiple movement strategies. Our study demonstrates that migration patterns in bats emerge from the interaction between intrinsic factors and external conditions, and highlights the importance of both energy efficiency and safety in shaping stopover departure decisions. The presence of multiple movement strategies complicates predictions of spatiotemporal occurrence, emphasising the need to account for behavioural variability in conservation planning, for example in the context of wind energy developments.

## INTRODUCTION

Several bat species undertake long-distance migrations, traveling hundreds to thousands of kilometres between their summer and winter ranges (Fleming and Eby, 2003). Because such distances cannot be completed in a single flight, bats alternate between migratory flights and stopovers to rest and refuel (Jonasson and Guglielmo, 2016; McGuire et al., 2012). Bats use torpor to conserve energy (Geiser, 2004), enabling them to depart on low fuel loads and use short stopovers in comparison to birds (Clerc and McGuire, 2021; McGuire et al., 2012), which often accumulate high fuel loads during extended stopovers (Alerstam and Lindström, 1990; Schmaljohann and Eikenaar, 2017). When embarking on a migratory flight from a stopover site, bats decide on the specific night, specific time within the night, and direction to depart. These decisions affect energy expenditure, time budget and mortality risk during migration (Alerstam and Lindström, 1990; Hedenström, 2009).

The overall spatiotemporal pattern of migration appears to be genetically controlled as bats likely migrate without guidance of related individuals (Baerwald and Barclay, 2016; Van Schaik et al., 2025) and migration occurs within distinct seasonal timeframes (Fleming and Eby, 2003; Lagerveld et al., 2024, 2023; Pettit and O’Keefe, 2017; Rydell et al., 2014).

In addition to their innate migration program, several intrinsic (e.g. sex and age) and external factors (e.g. environmental conditions) influence migratory decisions. Seasonal timing is both sex-dependent (Jonasson and Guglielmo, 2016; Lagerveld et al., 2024; Pētersons, 2004) and age-dependent (Pētersons, 2004; Strelkov, 1969). Environmental conditions such as wind support (Dechmann et al., 2017; Hurme et al., 2025; Lagerveld et al., 2024, 2021) as well as dry weather (Brabant et al., 2021; Lagerveld et al., 2023, 2014) promote departures, whereas strong headwinds, precipitation and overcast conditions induce stopovers (Hüppop and Hill, 2016). Additionally, bat migratory movements coincide with higher temperatures (Brabant et al., 2021; Hurme et al., 2025; Lagerveld et al., 2021; Pettit and O’Keefe, 2017; True et al., 2023), atmospheric pressure changes (Hurme et al., 2025), and may be associated with darker phases of the moon (Cryan and Brown, 2007; Lagerveld et al., 2023).

Despite growing knowledge of bat migration, key aspects of stopover departure decisions remain unresolved. In particular, it is unclear whether cloud cover affects departure decisions. Apart from being associated with precipitation, cloud cover may also hinder navigation by obscuring celestial cues. The role of crosswinds is also poorly known. Moreover, the specific departure time within the night is also not well understood. Bats generally depart shortly after sunset (Bach et al., 2022; Lagerveld et al., 2024; McGuire et al., 2012; True et al., 2023), yet it remains unclear whether environmental conditions influence within-night departure timing.

To fill parts of these knowledge gaps, we investigated autumn stopover departure decisions in the long-distance migratory *Nathusius’ pipistrelle Pipistrellus nathusii*. The species’ nursery roosts are primarily located in northeastern Europe, whereas most hibernation sites are distributed across southern and western Europe, including the United Kingdom (Pētersons, 2004; Russ, 2022; Strelkov, 1969). Most migrants are recorded in coastal regions (Ahlén et al., 2009; Ijäs et al., 2017; Voigt et al., 2023). This coastal concentration likely reflects barrier effects, as *Nathusius’ pipistrelles* generally follow shorelines to avoid large expanses of open water, and often perform back-and-forth movements along the coast while searching for a suitable location to cross over sea (Lagerveld et al., 2024). Autumn migration coincides with the mating season, during which males establish mating roosts along the flyway and remain there for extended periods (Gerell-Lundberg and Gerell, 1994).

We tagged bats at three locations along the Dutch coast during five autumn seasons (2018-2022) and tracked their movements using the Motus wildlife tracking system (Mitchell et al., 2025; Taylor et al., 2017). With the obtained tracking data, we (I) examined movement patterns, (II) analysed night-to-night departures in relation to the seasonal timing, environmental conditions and individual traits, and (III) analysed the within-night departures, in relation to the departure time relative to sunset and environmental conditions.

## MATERIAL & METHODS

### Bat tagging and tracking movements

Bats were captured and tagged over five consecutive autumn seasons (2018–2022), at three coastal sites in the Netherlands: Hoek van Holland (N 51.99, E 4.12), northern North Holland (N 52.84, E 4.71) and the Afsluitdijk (N 53.07, E 5.34). Captured bats were sexed, aged, weighted and measured following (Haarsma, 2008). Individuals assessed to be in poor condition (i.e. visible injuries, lots of parasites, damaged wing membranes) were released immediately.

In total, 566 bats were tagged between 20 August (day 232) and 19 October (day 292), including 369 from bat boxes during the day and 197 at night using Austbat harp traps (Faunatec) and mist nets (Solida Safety line – white). The dataset comprised 103 first-year females, 202 adult females, 91 first-year males and 170 adult males, with similar mean tagging dates across groups (days 255–257).

Tags were attached with skin adhesive (Sauer), ensuring total tag mass remained <5% of the bat’s body mass (Aldridge and Brigham, 1988). Bats were released on-site after the adhesive was set (after about 10 min). We used 0.35 g NTQB2-2 tags (2018–2019) and 0.26 g NTQB2-1 tags (2020–2022; LOTEK Wireless) with burst intervals of 5.9–8.9 s, transmitting uniquely coded signals at 150.1 MHz.

Movements were tracked using the Motus Wildlife Tracking System (Mitchell et al., 2025; Taylor et al., 2017), which gradually increased in the study area from 51 receivers in 2018 to 109 receivers in 2022 (Appendix S1). Detection data and receiver metadata were retrieved from www.motus.org (accessed 19 May 2025). To avoid false positives, detections with less than three consecutive signals were excluded. Additional manual filtering was conducted to exclude detections from receivers affected by high levels of radio noise (e.g. in ports);; or otherwise implausible detections (Appendix S2). All analyses were performed in R version 4.5.1 (R Core Team, 2025)

Receiver locations were used as proxies for the location of the bat at the timestamp when the strongest signal was received, except at the departure location where the timestamp of the last signal received was used, as migratory bats typically perform erratic movements prior to the actual departure (McGuire et al., 2012). We recorded both movements within one night as well as movements over multiple nights. Migratory flights (hereafter *flights*) were defined as a directed flight of 30 km or more. Of the 566 tagged bats, 23 individuals (4%) were not detected, 357 (63%) remained within a 30 km of the tagging site and 186 bats (33%) undertook *flights*. Flight bearings were calculated using the R package sf. (Pebesma, 2018).

As autumn migration is predominantly south westward, *flights* were classified as *regular* when they were oriented in southern or western directions, and as *return* when oriented in northern or eastern directions.

### Environmental data

Weather data were obtained from the Copernicus Climate Data Store (Hersbach et al., 2018) using the R package CopernicusClimate (De Vries, 2025), retrieved 2 July 2025. Data were provided as hourly raster fields (0.25° × 0.25°) and included 2 m air temperature [K], mean sea-level air pressure [hPa], total precipitation [m/h], cloud cover [%], eastward (U) and northward (V) wind speed components [m/s]. Wind data, provided for specific air pressure layers, were converted to altitude layers using the barometric equation (NASA, 1976). We used wind data at 200 m above sea level, as these better represent regional airflow patterns in comparison to near-surface (up to 100-150 m) winds (Garratt, 1992). We calculated the tailwind (wind support in the direction of travel) and crosswind components (wind perpendicular to the direction of travel), using the bearing between the departure and arrival location. When the bat was also detected in-between, the receiver closer to the departure point was chosen to estimate the bearing of departure. Temperature was converted to °C, precipitation to binary values (0 = dry, 1 = precipitation) and atmospheric pressure change was calculated relative to values 24 hours earlier. Lunar phase [degrees] and moon illumination fraction [%] were derived using the R package suncalc (Thieurmel, 2022).

#### (I) Movement patterns

We recorded *flights* of 186 bats, including two *flights* by 76 individuals, three *flights* by 17, four *flights* by seven, five *flights* by three, and six *flights* by a single individual. Flights were recorded for 8% of adult males (13 individuals), 43% of first-year males (39 individuals), 52% of first-year females (54 individuals), and 40% of adult females (80 individuals).

#### (II) Night-to-night departures

Of the 186 tracked bats, 28 initiated their first *flight* on the night of tagging and were excluded as handling may have influenced behaviour and earlier departures could not be ruled out. The remaining 158 bats, including 72 adult females, 49 first-year females, 10 adult males and 27 first-year males, were used to assess minimum stopover duration and the night-to-night departure probability.

Minimum stopover duration at the tagging site was defined as the interval between tagging and the last detection. Individuals tagged during daytime (captured in bat boxes) were assigned as first-time present in the previous night. Differences among the sex and age groups were tested using a Kruskal–Wallis test.

Night-to-night departure probability was analysed in relation to seasonal timing, environmental conditions and individual traits, using a Bayesian capture-recapture state-space model in JAGS (Gimenez et al., 2007; Kéry M. and Schaub M., 2011; Royle, 2008), following (Lagerveld et al., 2026) (*under review*). Migratory state of each individual was represented as a time series. Starting from the tagging date, the migratory state was assigned 1 when an individual has not yet departed from the tagging site and 0 on the night when it was detected at its destination. On nights between the first and last detection when an individual was not observed, the migration state was assigned NA (unknown). For these nights, the latent probability of migration was modelled. For individuals with multiple *flights*, only the first was used, representing departure from the stopover site. The dataset included 146 *flights* (92%) in the *regular* migration direction and 12 (8%) *return* flights.

The night-to-night departure probability was modeled as a likelihood of transition between the states in relation with night in year (mean-centered: linear and quadratic terms), environmental conditions and individual traits. Individual-level random intercepts were included to account for repeated observations. Migration probability was derived as 1−plogis of the linear predictor. Consequently, negative coefficients correspond to increased departure probability, while positive coefficients correspond to decreased departure probability.

Environmental covariates included tailwind and crosswind (both linear and quadratic terms), atmospheric pressure change, temperature, precipitation and cloud cover, averaged between sunset and two hours post-sunset. Temperature was correlated with night in year and was therefore included in the model as residuals from a regression on night in year.

Cloud cover is often associated with precipitation, but may also hinder orientation and navigation by obscuring celestial cues. To separate these overlapping effects, we included a partial interaction term between cloud cover and precipitation. Non-zero precipitation was set as the baseline, enabling the model to estimate the effect of cloudiness under dry conditions. Lunar phase was treated as a cyclic covariate by including both sine and cosine terms.

Individual covariates included sex (male/female), age (adult/first-year), along with their interactions with night in year (linear and quadratic terms). Body mass index (BMI) was calculated as a within-sex-age-group residual of individual body mass at tagging on arm length (Appendix S3)

Aiming to define the most influential factors affecting departure decisions, we started with a model that included all candidate covariates and their interactions. To avoid possible spurious outcomes due to the limited sample size/covariate number ratio, we sequentially removed the covariates with the weakest support, based on the overlap of the 95% posterior credible intervals of parameters with zero and the corresponding evidence ratios. Convergence was confirmed for all parameters 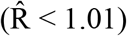 Model simplification continued until the model with the lowest deviance information criterion (DIC) was obtained. The final model was:

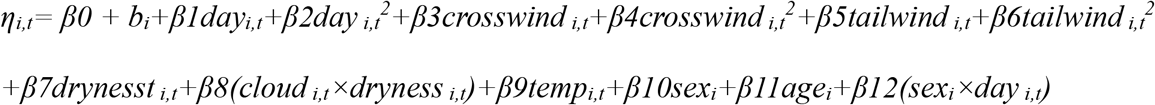

with individual random intercept effects:

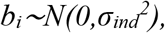

where *p*_*i,t*_*=1−logit*^*−1*^*(η*_*i,t*_) is the probability of migration of individual i on day t.

#### (III) Within-night departures

The same modelling approach was used to analyse the within-night departure probability. For each individual bat with a known departure time (n = 102), we modelled the hourly probability of a transition between states (not yet departed = 1 versus departed = 0) on the night of departure as a function of time since sunset, and weather conditions affecting flight conditions (tailwind, crosswind and precipitation), food availability (temperature) and ambient light levels (cloud cover and moon illumination). The encounter histories started one hour before sunset. Two outliers (departures at 315 and 427 min after sunset) were removed from the analysis. The full model was the model with the lowest DIC:

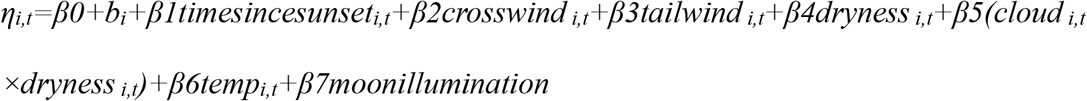

with individual random intercept effects:

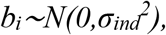

where *p*_*i,t*_*=1−logit*^*−1*^*(η*_*i,t*_) is the probability of migration of individual i on day t.

## RESULTS

### (I) Movement patterns

Of the 186 tracked bats, 169 individuals (91%) initiated migration in the regular autumn direction (between south and west-northwest), while 17 individuals (9%) performed return *flights* (north to northeast). Among those initially migrating in the regular direction, 30 individuals (18%) later performed one or more return *flights*. Conversely, five individuals (29%) that initially made return *flights* subsequently migrated in the regular direction. Overall, 139 bats (75%) exclusively followed the regular migration direction, 12 bats (6%) performed only return *flights* and 35 bats (19%) exhibited back-and-forth *flights*. Three individuals were detected in the United Kingdom, while the remaining 183 were recorded solely on the European mainland (Figure 1 and Appendix S4).

**Figure 1.**
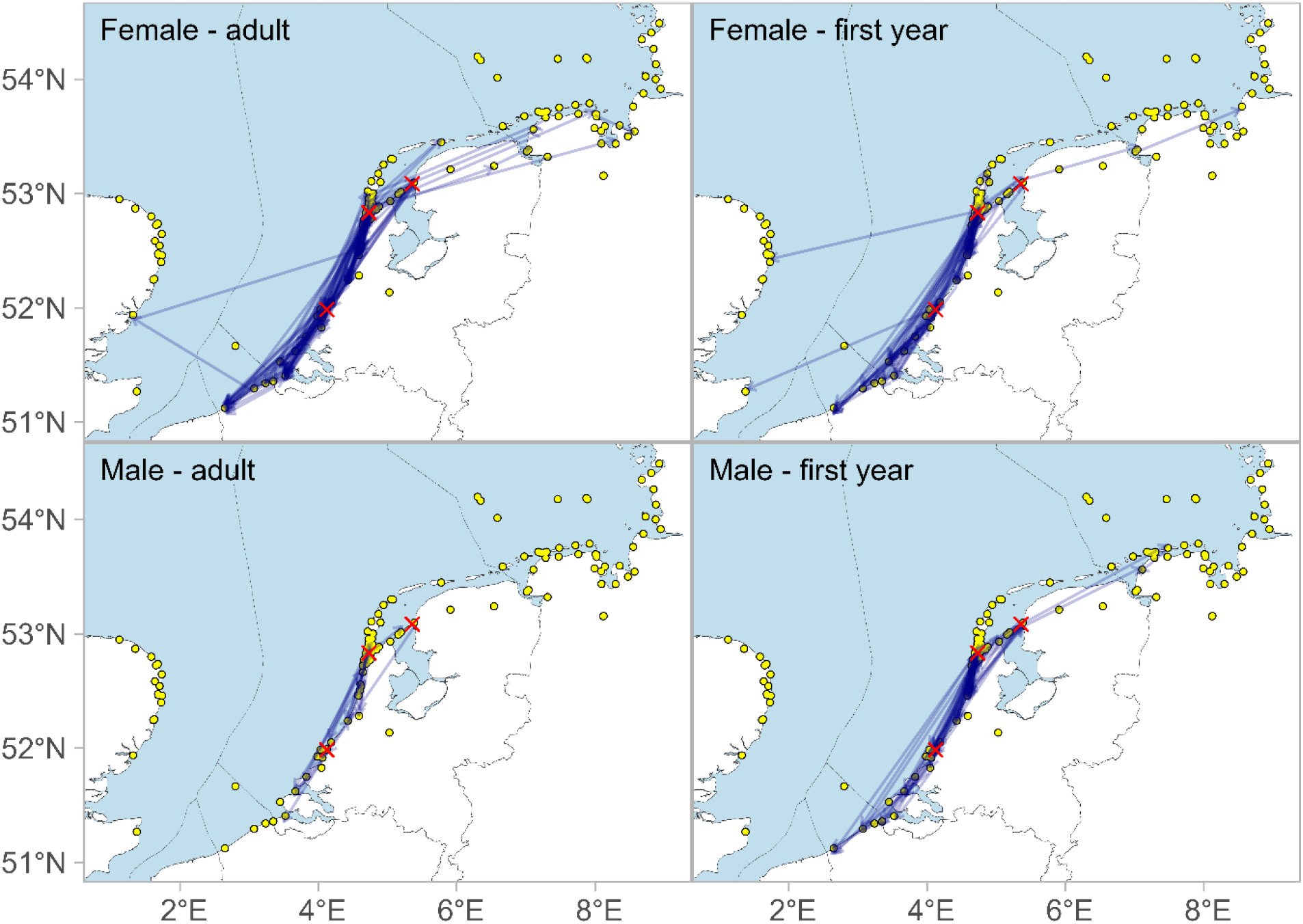
Movements >30 km from the tagging sites (n = 186). Receivers are indicated as yellow dots, tagging locations as red crosses. The flights are represented as straight lines between start and end point, not necessarily reflecting the actual flight paths. See Supplementary file 4 for the movements of each individual.

Among bats exhibiting back-and-forth movements, 26% covered distances exceeding 300 km (Figure 2), including one adult female that travelled at least 715 km within three weeks. This individual travelled from the Afsluitdijk (N53.10 E5.48) to southern Belgium (N51.29 E3.07), subsequently crossed the North Sea to the east coast of the United Kingdom (N51.94 E1.32), and subsequently returned to the tagging site (Figure 1 and Appendix S4: Deployment 42408). We found no evidence that the occurrence of back-and-forth *flights* differed among the sex and age groups (Fisher’s exact test, p = 0.19).

**Figure 2.**
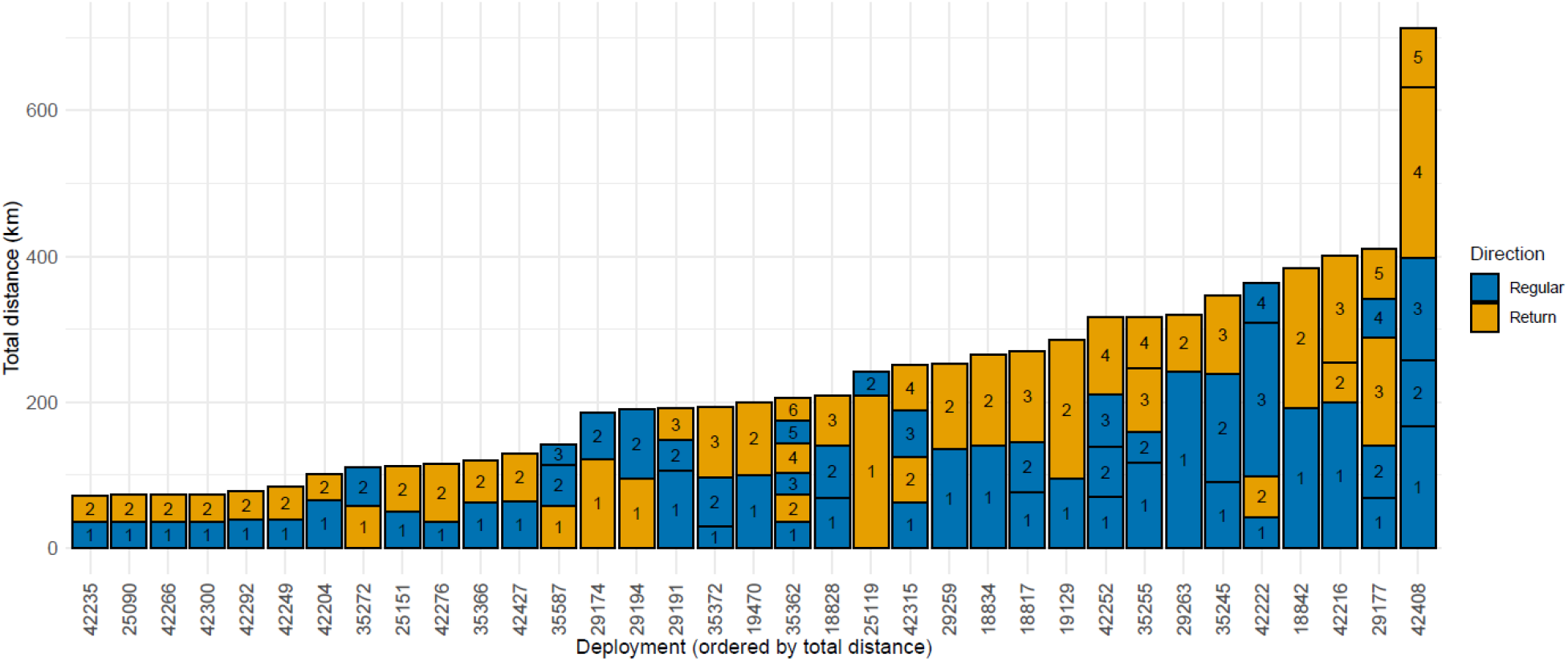
Minimum distance back-and-forth flights by flight number and direction

### (II) Night-to-night departures

The overall median minimum stopover duration was two days. A considerable proportion of bats (46%) stayed only one day, while maximum observed duration was 58 days. Stopover duration did not differ significantly among sex and age groups (Kruskal–Wallis: χ^2^(3) = 5.71, p = 0.127). In general, minimum stopover duration generally increased as the season progressed (Figure 3).

**Figure 3.**
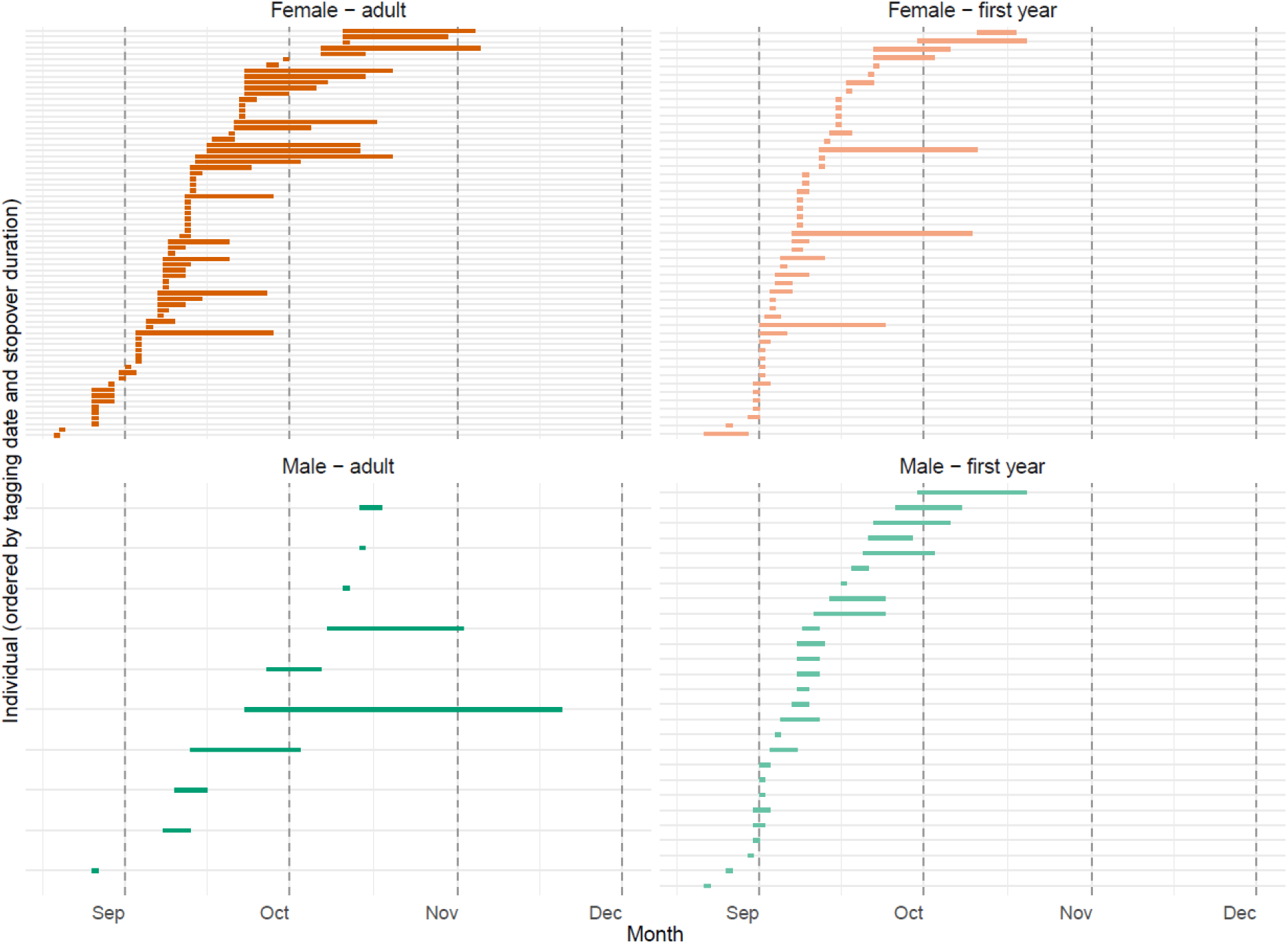
Minimum stopover duration for each individual per sex and age group, ordered by tagging date and stopover duration.

The model showed strong support for a non-linear seasonal pattern with higher departure probability in the beginning and at the end of the migration period (*β2* = −0.002, 95% CI: −0.003 to −0.001, f =1.00). Males and adults tended to have lower departure probabilities than females and first-year individuals, although these effects were weakly supported (β10 = 0.747, 95% CI: −0.007 to 1.615, f =0.97; β11 = 0.538, CI: −0.134 to 1.274, f =0.94. However, later in the season males had a significantly lower probability of migration (β12 = 0.049, CI: 0.006 to 0.096, f =0.99, Figure 4).

**Figure 4.**
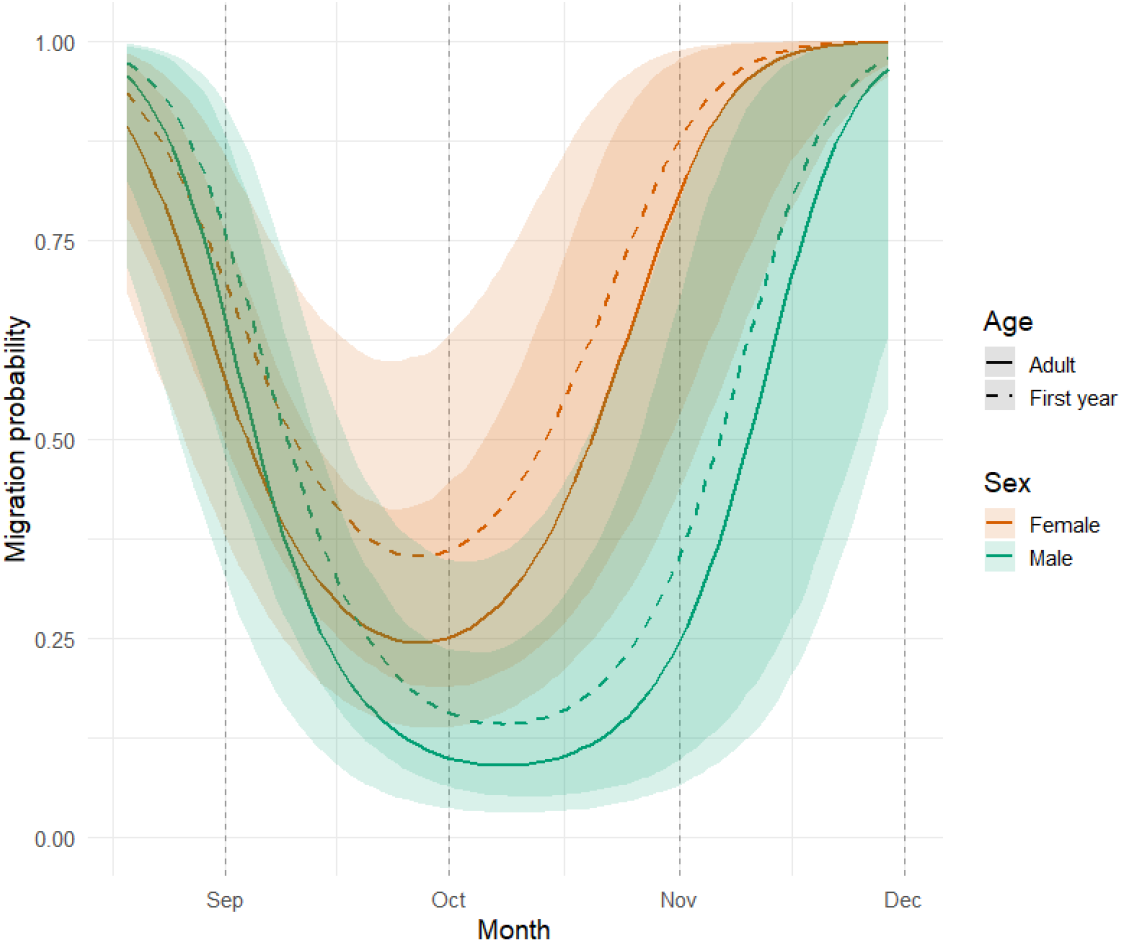
Departure probability throughout the season for each sex and age group

Wind conditions had pronounced non-linear effects on departure decisions (β6 = 0.009, CI: 0.003 to 0.016, f =1.00; β4 = 0.026, CI: 0.017 to 0.036, f =1.00), indicating the highest departure probability during moderate tailwinds and low (near-zero) crosswinds (Figure 5). The maximum migration probability was predicted under tailwinds of 9.9 m/s and crosswinds of 0.9 m/s from the left, which corresponds to eastern winds for most flights (92%) in the dataset.

**Figure 5.**
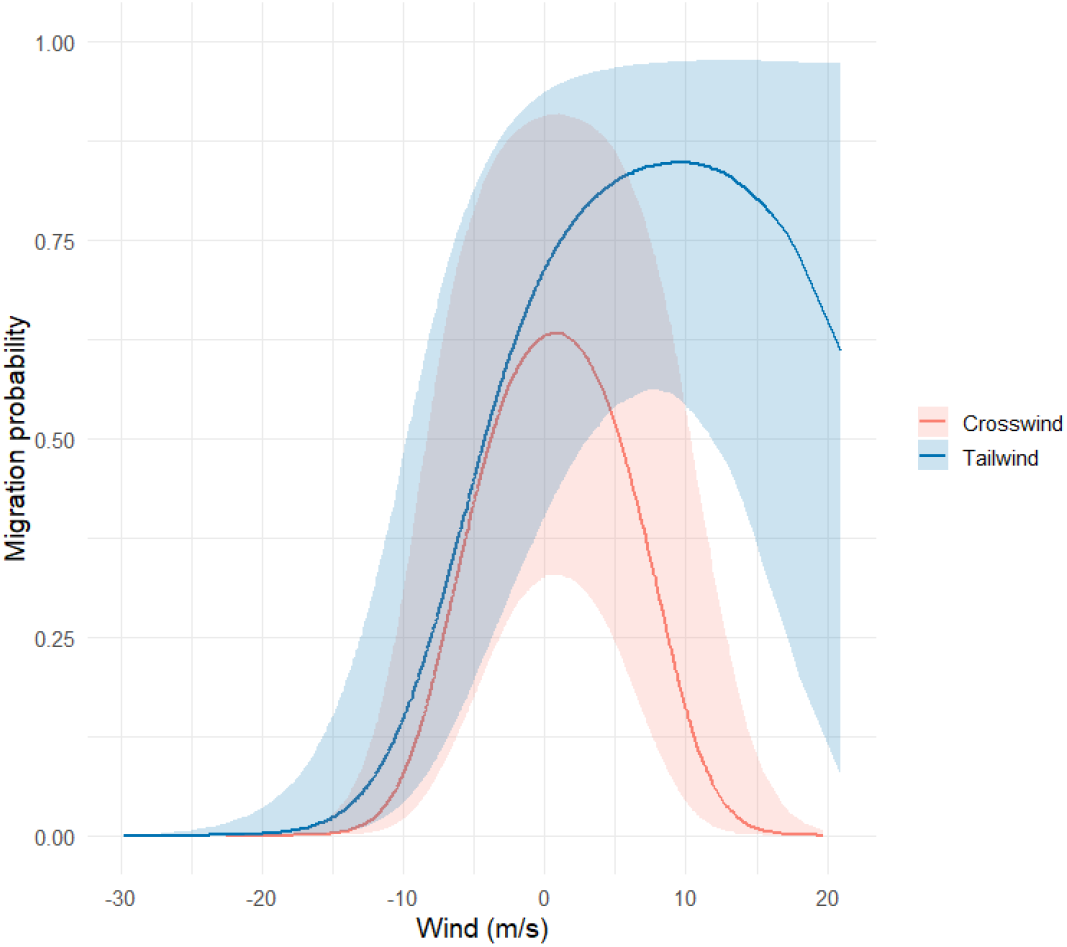
Departure probability depending on tailwind and crosswind

Precipitation strongly reduced departure probability (β7 = −1.151, CI: − −1.951 to −0.344, f =1.00). The positive cloud cover × precipitation interaction (β8 = 1.262, CI: 0.071 to 2.498, f =0.98), showed a statistically significant negative effect of cloud cover under dry conditions on departure probability. Temperature showed a weak, but not significant, effect (β9 = −0.119, CI: −0.300 to 0.056, f =0.91). Substantial individual-level variation was observed (σ_ind = 1.096, CrI: 0.538 to 1.727, f =1.00) (Appendix S5).

### (III) Within-night departures

Most bats (88%) departed within two hours after sunset and 9% between 2-3 hours after sunset. The remaining three individuals departed 182, 315 and 427 min after sunset (Figure 6).

**Figure 6.**
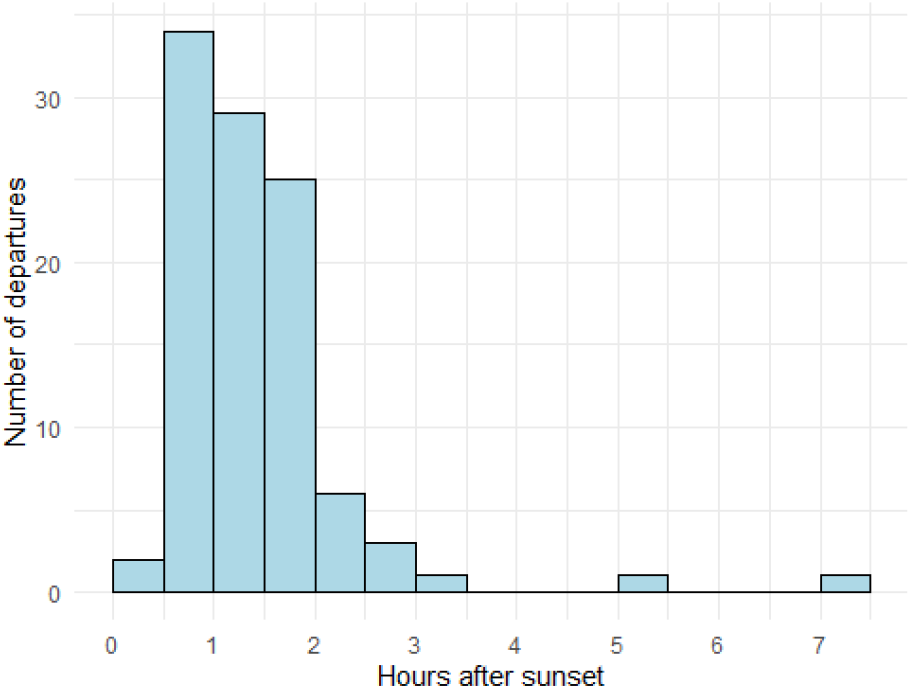
departure time after sunset

The model showed a strong effect of time since sunset on departure probability (β = −24.08, 95% CI: −37.65 to −12.23, f = 1.00), indicating that departures were concentrated early in the night. Tailwind had a clear positive effect (β = −0.80, 95% CI: −1.70 to −0.12, f = 0.99), meaning that increased headwinds delayed departures (Figure 7). Crosswind showed a weaker and uncertain negative effect (β = 0.49, 95% CI: −0.23 to 1.38, f = 0.91), suggesting a tendency to delay nocturnal departures. Other environmental covariates were not supported with wide confidence intervals overlapping with zero. The individual variation was very large (σ_ind = 13.94 (CrI: 6.77–22.61)), suggesting substantial individual heterogeneity not explained by covariates in the model (Appendix S5).

**Figure 7.**
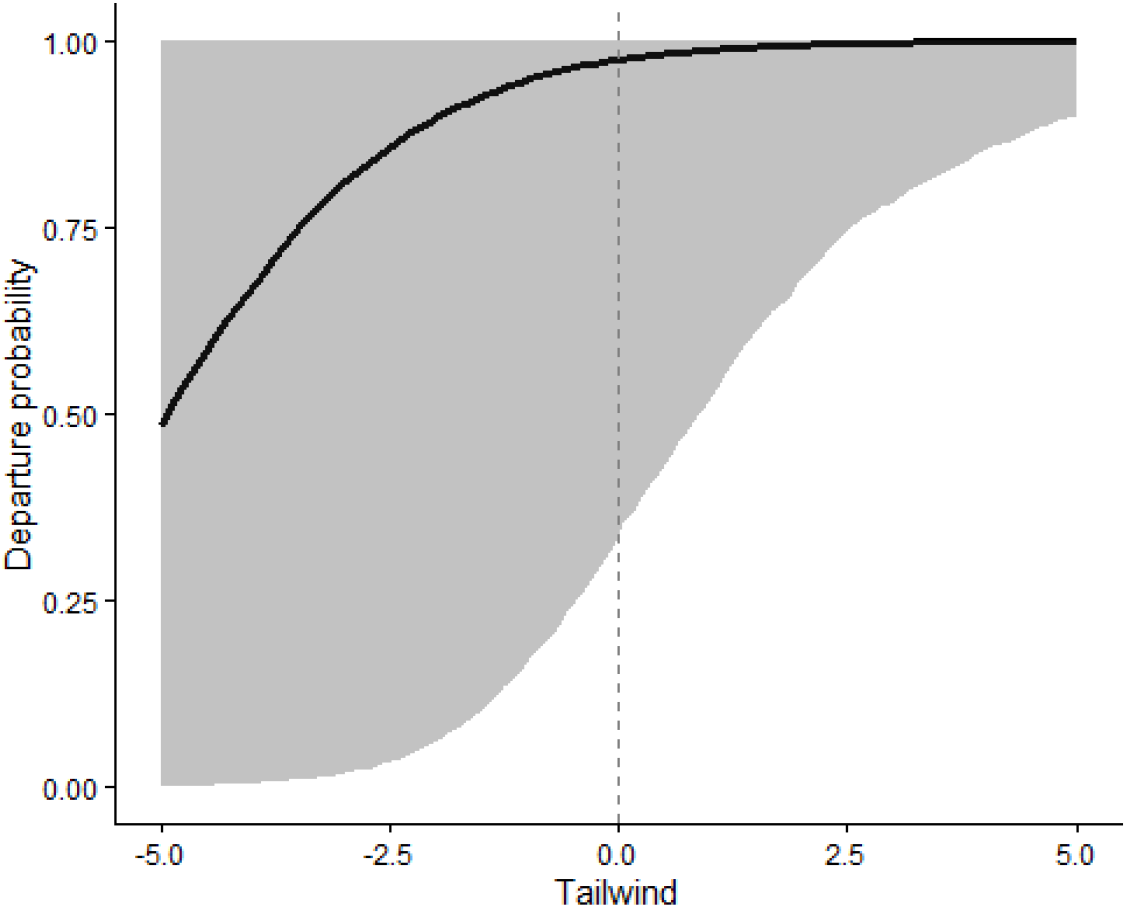
migration probability depending on tailwind. Note that negative tailwind refers to headwind.

## DISCUSSION

We tracked Nathusius’ pipistrelles during autumn migration to examine movement patterns and stopover departure decisions at both night-to-night and within-night scales. Movement patterns suggest multiple overlapping strategies, including migration towards hibernation sites, barrier-type movements, and mating-related movements. Seasonal migration probability peaked in late August, declined during the mating period, and increased again from mid-October onwards, indicating that mating activities likely constrain migration during early autumn. Later in the season, females departed earlier than males and first-year bats of both sexes tended to depart earlier than adults. Regardless of timing, bats departed under moderate tailwinds and dry conditions to optimize energy expenditure, while crosswinds and cloud cover were avoided to enhance safety. Bats typically departed soon after sunset, while headwinds delayed nocturnal departure, further highlighting energy-efficiency, even at fine temporal scales.

### (I) Movement patterns

Most bats (75%) migrated in the expected southwest direction, whereas 19% showed back-and-forth movements along the coast, and 6% moved in the opposite direction, possibly reflecting return movements where the southwest-bound movement was not recorded. These back-and-forth flights likely reflect barrier movements to find a suitable location to cross over sea (cf Lagerveld et al., 2024). Also in other regions, movements of migratory bats along large expenses of open water have been observed, opposite or perpendicular to the general migration direction (McGuire et al., 2012; Voigt et al., 2023). The observed back-and-forth movements suggest that at least part of the population may attempt to cross the North Sea towards the United Kingdom. The small number of individuals detected there is likely underestimated due to limited receiver coverage and rapid inland dispersal.

We also recorded long-distance roundtrips, including one individual that crossed the North Sea and returned three weeks later. This indicates that some movements are not strictly migratory, but may also be driven by other factors such as mating. Together, these findings suggest that autumn movement patterns reflect migration, barrier and mating-related movements.

### (II) Night-to-night departures

Departure decisions were shaped by both intrinsic and environmental factors, with mating behaviour likely structuring seasonal patterns. Adult males were largely sedentary (8% departed), consistent with their role at mating sites, whereas females and first-year bats showed substantially higher departure rates, indicating more transient individuals in these groups. Across all groups, departure probability peaked in late August, declined during September–mid-October, and increased thereafter (Figure 4). This decline coincides with the mating season in western Europe (cf Lina and Reinhold, 1997), during which males establish mating roosts where they remain extended periods (Gerell-Lundberg and Gerell, 1994) Females visit these roosts for one to several days. Among them, first-year individuals who attain sexual maturity at an age of about three months (Gerell-Lundberg and Gerell, 1994).

First-year males, which reach sexual maturity after about one year (Gerell-Lundberg and Gerell, 1994), also join these temporary harems (Van Schaik et al., 2025). The overlap between mating season and reduced migration probability strongly suggest that mating activities constrain migratory movements from early September to mid-October.

In the breeding areas, adult females depart first, subsequently the first-year bats and last the adult males (Strelkov, 1969). At the end of the migration period first-year bats of both sexes tended to depart earlier than adults, with females leaving earlier than males. This pattern suggests that the adult females migrate slower than first-year females, possibly because adult females invest more time in mating activities, resulting in a convergence in departure timing later in the season in the western parts of their range. First-year males migrate more slowly than first-year females and exhibit departure timing similar to that of adult males, which depart last, as has also been observed in other regions (cf Pētersons, 2004; Strelkov, 1969).

Departure probability was not related to body mass index, further supporting the idea that departures do not depend on high fuel loads (cf Clerc and McGuire, 2021; McGuire et al., 2012). Weather conditions strongly influenced departures. Bats preferentially departed with moderate tailwinds, confirming the importance of wind support for migratory bats cf Dechmann et al., 2017; Hurme et al., 2025; Lagerveld et al., 2026, 2024, 2021). Numerous studies also identify wind support as a major determinant of migratory departures in songbirds (Brust et al., 2019; Packmor et al., 2020; Rüppel et al., 2023). When songbirds encounter crosswinds during flight over land they often drift sideways, but compensate for drift near the coastline (Horton et al., 2016). Migrating songbirds over sea avoid crosswinds (Packmor et al., 2020). Bats departures peaked in low crosswinds (0.9 m/s) from the left side, which for most of the flights in the dataset (91%) was from the east. This suggests that bats generally avoid crosswinds and actively compensate for drift to minimise the risk of displacement over open water.

Precipitation reduced departure probability, likely due to increased energetic costs (cf Voigt et al., 2011) and reduced foraging and orientation efficiency (Geipel et al., 2019). Similarly, cloud cover decreased migration probability, suggesting that bats prefer clear skies to facilitate orientation. Some species of bats are known to use the Earth’s magnetic field for orientation, which is calibrated by the solar azimuth at sunset (Holland et al., 2010; Schneider et al., 2023).

Migratory bat activity is often associated with higher temperatures (Brabant et al., 2021; Hurme et al., 2025; Lagerveld et al., 2021; Pettit and O’Keefe, 2017; True et al., 2023), pointing to a relationship with insect availability, which may further be modulated by the lunar phase (cf Lagerveld et al., 2023). In our study we found a positive, but not significant, effect of increased temperatures on seasonal departures, while lunar phase had no detectable effect.

Increasing atmospheric pressure generally indicates stable weather and favourable flight conditions over the next few days. Several bird migration studies have observed higher departure probabilities when atmospheric pressure increases (Cooper et al., 2023b; Rüppel et al., 2023; Woodworth et al., 2015). Our results did not show an effect of changes in atmospheric pressure on the departure probability. This may indicate that bats rely on current local weather conditions, rather than taking anticipated future weather into account.

Overall, our findings indicate that the seasonal migration patterns are strongly modulated by reproductive behaviour, while their night-to-night departure decision is based on current atmospheric conditions: tailwinds and dry weather to optimize energy use, clear skies and minimal crosswinds to enhance safety, and possibly higher temperatures to increase foraging opportunities.

### (III) Within-night departure timing

Most bats (88%) departed within two hours after sunset, consistent with previous studies (cf. Bach et al., 2022; Lagerveld et al., 2024; McGuire et al., 2012; True et al., 2023). This narrow departure window mirrors patterns observed in migratory songbirds (Cooper et al., 2023a; Klinner et al., 2025; Müller et al., 2016) and suggests shared orientation mechanisms, such as compass calibration at sunset (Cochran et al., 2004; Schneider et al., 2023).

Early departures potentially increase the duration of the migratory flight and the distance covered (Müller et al., 2016). Songbirds embarking on long-distance migratory flight depart generally within two hours after sunset, while individuals performing non-migratory movements away from the stopover site depart later and less synchronously over the course of the night (Cooper et al., 2023a). Bat departure timing showed little variation, with headwinds being the only factor delaying departure. This further highlights the importance of energy efficiency, even at fine temporal scales, and suggests that within-night departure timing is highly constrained.

## CONCLUSION

Our study demonstrates that migration patterns in bats emerge from the interaction between intrinsic factors and external conditions. Seasonal patterns in autumn are strongly shaped by reproductive behaviour, whereas night-to-night departure decisions are primarily driven by immediate atmospheric conditions that optimise energy efficiency and navigational safety. The diversity of observed movements, including migration towards lower latitudes, coastal barrier movements and long-distance roundtrips, indicates the use of multiple movement strategies, complicating predictions of their spatiotemporal occurrence. Understanding this behavioural variability will be essential for developing effective conservation strategies, particularly in regions facing increasing anthropogenic pressures such as wind energy developments.

## Data availability

The data and code supporting this study will be made available upon publication of the manuscript.

## Contributions

Study design: Sander Lagerveld, Fieldwork: Karine Stienstra, Bart C.A. Noort and Sander Lagerveld. Data analysis: Julia Karagicheva, Sander Lagerveld, Pepijn de Vries, Eldar Rakhimberdiev. Original draft: Sander Lagerveld and Julia Karagicheva. Editing and review: all authors. Funding: Sander Lagerveld, Martin J.M. Poot, Frank van Langevelde.

## Ethics

Fieldwork was conducted under Nature Legislation Permit No. 2018-057682 (Wageningen Marine Research) and in accordance with Animal Welfare Protocols AVD248002016459 / VZZ-18-005 (2018–2020) and AVD24800202114476 / VZZ-2021-001 (2021–2023; Dutch Mammal Society).

## Acknowledgements

We thank the more than 150 individuals who contributed to this study. In particular, we are grateful to Jan Boshamer, Daan Dekeukeleire, Anne-Jifke Haarsma, René Janssen, and all volunteers for their efforts during tagging activities. Maurice la Haye and Nanneke van der Wal provided the animal welfare protocols. Installation and maintenance of the Motus receiver network were carried out primarily by Martijn Keur and Cor Sonneveld (Netherlands), René Janssen (Belgium), Ewan Parsons (UK), and Heinz-Hinrich Blikslager, Thomas Klinner, Thomas Mertens, Mario de Neidels, and Florian Packmor (Germany). We thank the Motus community and Birds Canada for coordination and support, and honour the memory of Ommo Hüppop for his invaluable contribution to the Motus network. We are also grateful to Martin Baptist, Meik Verdonk and Henri Zomer for their constructive comments, which greatly improved the manuscript.

