## Appendix S1 for "Stopover departure decisions and movement patterns of migratory bats during autumn migration"

### Appendix S1: southern North Sea Motus receiver network

The Motus Wildlife Tracking System (www.motus.org) is an international collaborative research network that employs automated radio telemetry. It is specifically designed to track the movements of smaller animals, such as bats and birds, during a prolonged period of time without the need to recapture them. Motus is therefore widely used for migration studies. The system operates through a network of stationary automated receiving stations and lightweight coded VHF radio tags, used to identify and track individual animals. In Europe 150.1 MHz is used as common frequency, while 166 MHz is used in Northern & Southern America. Motus is a program of Birds Canada in partnership with collaborating researchers and organizations. Figures S1-1 – S1-5 show the development of the receiver network in the southern North Sea area from 2018 - 2022. Generally receivers are located along coastlines and are virtually absent from inland locations. The highest receiver coverage is in northwestern Germany and along the Westcoast of the Netherlands.

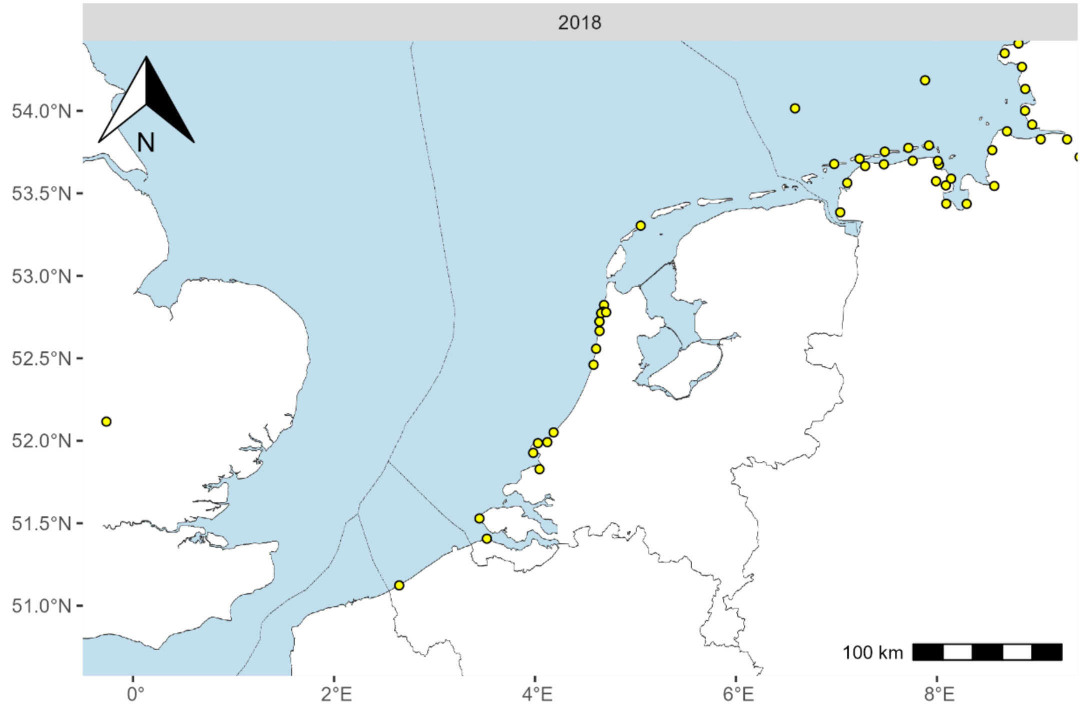


*Figure S1-1: Operational receivers (n=51) in December 2018*

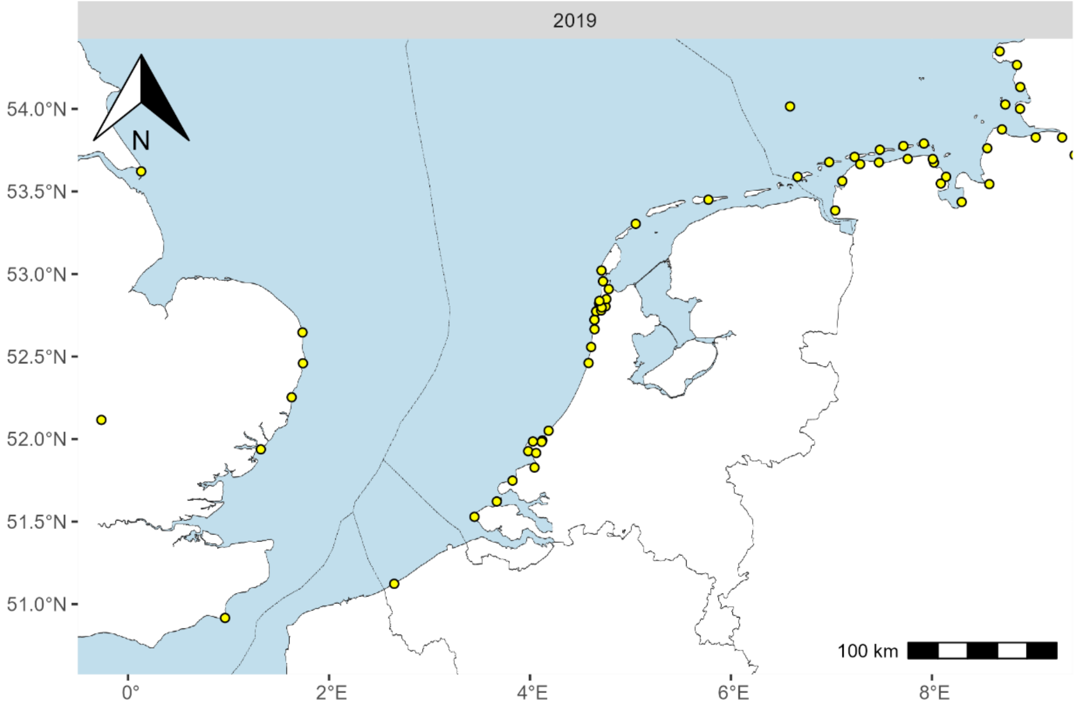


*Figure S1-2: Operational receivers (n=61) in December 2019*

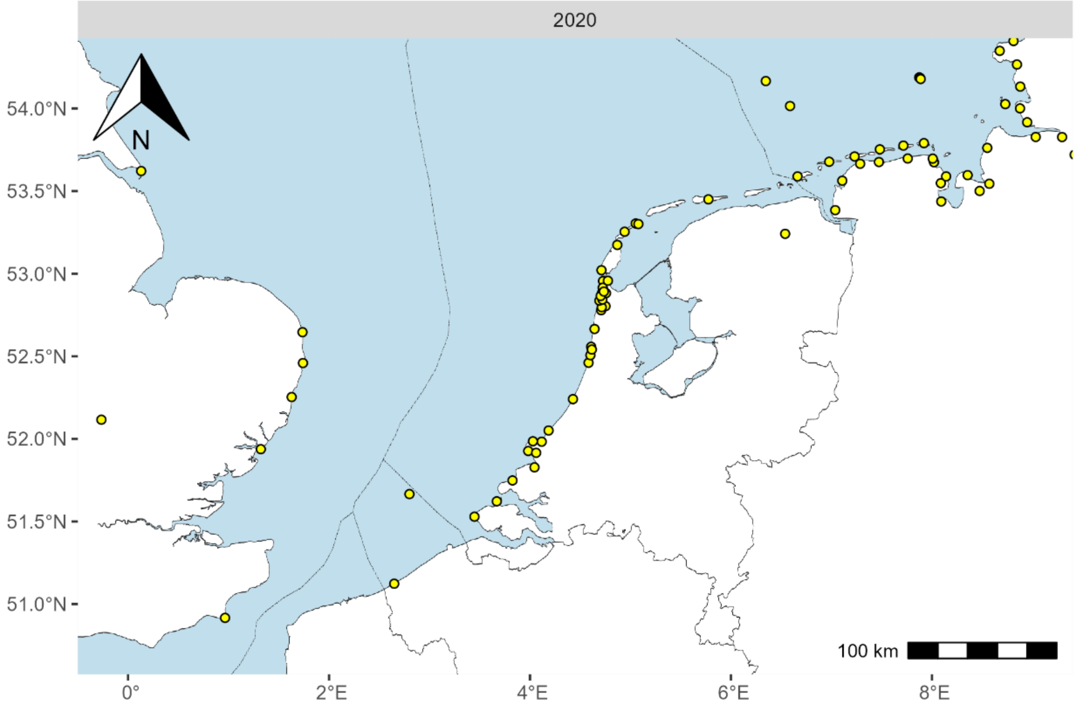


*Figure S1-3: Operational receivers (n=79) in December 2020*

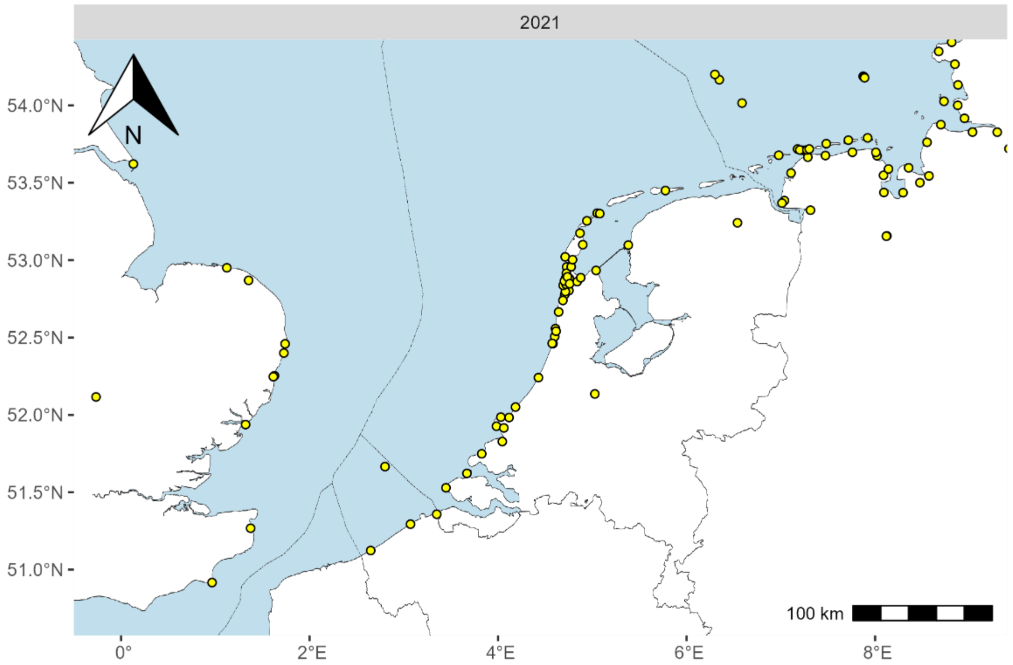


*Figure S1-4: Operational receivers (n=103) in December 2021*

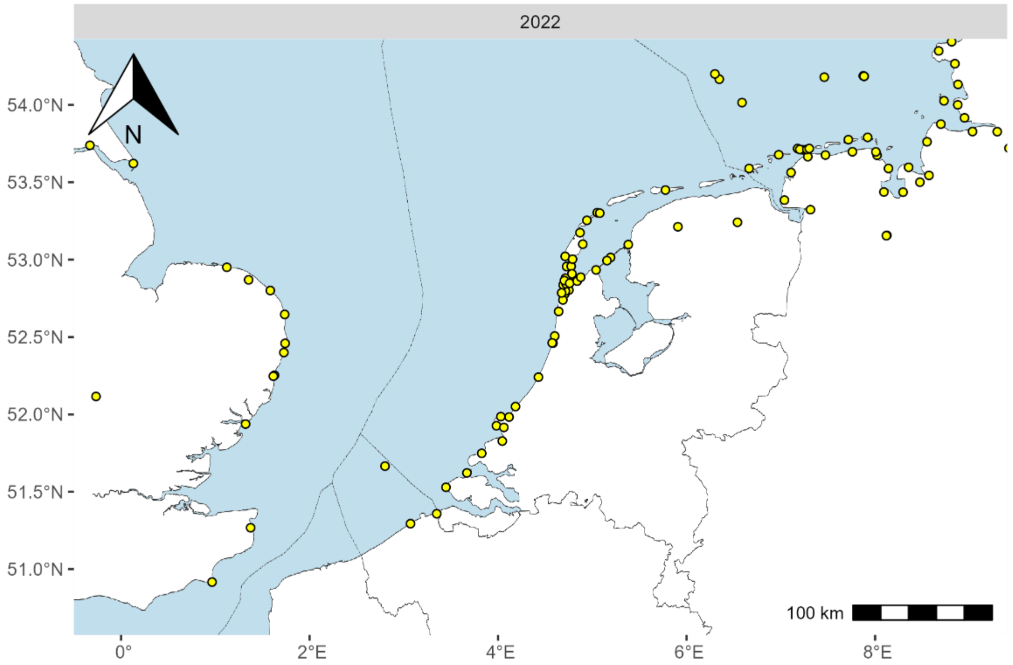


*Figure S1-5: Operational receivers (n=109) in December 2022*
