## Appendix S2 for "Stopover departure decisions and movement patterns of migratory bats during autumn migration"

### Appendix S2: removal of false positives

Detections with run lengths < 3 were filtered out, in accordance with the MOTUS R Book (Birds Canada, 2022). In addition, unlikely detections were removed from specific receivers at locations with high levels of radio noise (table S2-1).

*Tabe S2-1: false positives*

| Year | Tag Deployment | Receiver Deployment | Receiver | Date | Comments |
|---|---|---|---|---|---|
| 2018 | 18817 | 4130 | Bremerhaven | all | lots of false positives on this receiver in 2018 |
| 2018 | 18817 | 4315 | Castricum aan zee | all | lots of false positives on this receiver in 2018 (likely caused by cell phone mast) |
| 2018 | 18818 | 5023 | 29_Sillenstede | all | unlikely (one detection during daylight hours) |
| 2018 | 18840 | 4315 | Castricum aan zee | all | lots of false positives on this receiver in 2018 (likely caused by cell phone mast) |
| 2018 | 18840 | 4638 | Den Helder Vuurtoren | all | during daylights hours |
| 2018 | 18840 | 4454 | Ijmuiden vuurtoren | all | lots of false positives on this receiver from 31/10 onwards (also tags from abroad) |
| 2018 | 18840 | 4469 | 13_Ruthenstrom | all | during daylights hours |
| 2018 | 18840 | 4130 | Bremerhaven | all | lots of false positives on this receiver in 2018 |
| 2018 | 18844 | 4315 | Castricum aan zee | all | lots of false positives on this receiver in 2018 (likely caused by cell phone mast) |
| 2018 | 18852 | 4367 | 26_Utlandshörn | after 30-9-2018 00:00 | detections of a dropped off tag after 2018-09-29 |
| 2018 | 18859 | 4454 | Ijmuiden vuurtoren | all | 1 unlikely, 1 during daylight hours |
| 2018 | 18859 | 4315 | Castricum aan zee | all | lots of false positives on this receiver in 2018 (likely caused by cell phone mast) |
| 2018 | 18861 | 4892 | Huiberts | after 5-9-2018 00:00 | detections of a dropped off tag after 2018-09-04 |
| 2018 | 19022 | 4711 | Noorderhaven Julianadorp | after 7-10-2018 00:00 | detections of a dropped off tag after 2018-10-06 |
| 2018 | 19023 | 3702 | 23_Carolinensiel | all | unlikely |
| 2018 | 19023 | 4892 | Huiberts | after 13-9-2018 00:00 | detections of a dropped off tag after 2018-09-12 |
| 2018 | 19092 | 4532 | HelgolandFG1 | all | 1 unlikely (15/10, together with deployment 19119) |
| 2018 | 19093 | 4817 | Jomfruland Bird Observatory | all | late season unlikely detections (also during the day) in Norway |
| 2018 | 19093 | 4817 | Jomfruland Bird Observatory | all | unlikely (in Norway), also during daylight hours |
| 2018 | 19095 | 3943 | 08_Büsum | all | false positives of 5 tags on the same day (9/10/2018) |
| 2018 | 19099 | 3943 | 08_Büsum | all | false positives of 5 tags on the same day (9/10/2018) |
| 2018 | 19099 | 3943 | 08_Büsum | all | false positives of 5 tags on the same day (9/10/2018) |
| 2018 | 19100 | 4556 | 116_Borkum 1 | all | unlikely |
| 2018 | 19100 | 4556 | 116_Borkum 1 | all | unlikely |
| 2018 | 19101 | 3943 | 08_Büsum | all | false positives of 5 tags on the same day (9/10/2018) |
| 2018 | 19107 | 3943 | 08_Büsum | all | false positives of 5 tags on the same day (9/10/2018) |
| 2018 | 19119 | 4532 | HelgolandFG1 | all | 1 unlikely, (15/10, together with deployment 19092) |
| 2018 | 19120 | 4310 | ECN | after 15-10-2018 00:00 | detections of a dropped off tag after 2018-10-14 |
| 2019 | 25079 | 5624 | Single RX receiver_2 (Lund) | all | unlikely (in Sweden), also during daylight hours |
| 2019 | 25079 | 5376 | Ekologihuset | all | unlikely (in Sweden), also during daylight hours, and during the time frame it was detected in the Netherlands |
| 2019 | 25099 | 5624 | Single RX receiver_2 | all | unlikely (one detection during daylight hours) |
| 2019 | 25136 | 4532 | HelgolandFG1 | all | 1 unlikely (10/9 during daylight hours) |
| 2019 | 25169 | | all receivers | all | detections of a dropped off tag after 2019-09-04 |
| 2019 | 25177 | 5503 | Callantsoog Inland | after 27-8-2019 00:00 | detections of a dropped off tag after 2019-08-26 |
| 2019 | 25179 | 5463 | t Zand | after 18-9-2019 00:00 | detections of a dropped off tag after 2019-09-17 |
| 2019 | 25179 | 5621 | Eendekooi 't Zand | after 18-9-2019 00:00 | detections of a dropped off tag after 2019-09-17 |
| 2019 | 25181 | | all receivers | all | detections of a dropped off tag after 2019-09-04 |
| 2019 | 25220 | 5621 | Eendekooi 't Zand | after 25-9-2019 00:00 | detections of a dropped off tag after 2019-09-24 |
| 2019 | 26144 | 5621 | Eendekooi 't Zand | after 1-10-2019 06:00 | detections of a dropped off tag after 2019-10-01 06:00 |
| 2019 | 26146 | 5621 | Eendekooi 't Zand | after 29-9-2019 01:30 | detections of a dropped off tag after 2019-09-29 01:30 |
| 2019 | 26155 | 5376 | Ekologihuset | all | unlikely (in Sweden), during daylight hours |
| 2019 | 29197 | 4892 | Huiberts | after 16-9-2020 05:00 | detections of a dropped off tag after 2019-09-16 05:00 |

*Tabe S2-1: false positives (continued)*

| Year | Tag Deployment | Receiver Deployment | Receiver | Date | Comments |
|---|---|---|---|---|---|
| 2020 | 29174 | 6185 | 120_HelgolandFG1 | all | numerous false positives of various tags from several countries from 24/10/2020 onwards |
| 2020 | 29206 | 5463 | t Zand | after 17-9-2020 00:00 | detections of a dropped off tag after 2020-09-16 05:00 |
| 2020 | 29206 | 5621 | Eendekooi 't Zand | after 17-9-2020 00:00 | detections of a dropped off tag after 2020-09-16 05:00 |
| 2020 | 29213 | 5376 | Ekologihuset | all | unlikely (one detection in southern Sweden) |
| 2020 | 29214 | 7341 | Castricum Ringbaan | all | from 7/11 onwards, numerous false positives of various tags from several countries, caused by the presense of several tags at the ringing site |
| 2020 | 29215 | 7341 | Castricum Ringbaan | all | numerous false positives of various tags from several countries from 6/11/2020 onwards |
| 2020 | 29217 | 4711 | Noorderhaven Julianadorp | after 4-10-2020 00:00 | detections of a dropped off tag after 2020-10-04 |
| 2020 | 29220 | 7315 | Vlieland Kazerne | all | Likely caused by tagged songbirds present al this location (08/11/2020) |
| 2020 | 29224 | 7341 | Castricum Ringbaan | all | numerous false positives of various tags from several countries from 6/11/2020 onwards |
| 2020 | 29229 | 7341 | Castricum Ringbaan | all | numerous false positives of various tags from several countries from 6/11/2020 onwards |
| 2020 | 29230 | 7341 | Castricum Ringbaan | all | numerous false positives of various tags from several countries from 6/11/2020 onwards |
| 2020 | 29231 | 7341 | Castricum Ringbaan | all | from 7/11 onwards, numerous false positives of various tags from several countries, caused by the presense of several tags at the ringing site |
| 2020 | 29232 | | all receivers | all | detections of a dropped off tag after 2020-10-22 |
| 2020 | 29233 | 6185 | 120_HelgolandFG1 | all | numerous false positives of various tags from several countries from 24/10/2020 onwards |
| 2020 | 29244 | 6133 | Callantsoog Inland | after 22-9-2020 00:00 | detections of a dropped off tag after 2020-09-21 |
| 2020 | 29275 | 7341 | Castricum Ringbaan | all | from 7/11 onwards, numerous false positives of various tags from several countries, caused by the presense of several tags at the ringing site |
| 2020 | 29280 | 7341 | Castricum Ringbaan | all | from 7/11 onwards, numerous false positives of various tags from several countries, caused by the presense of several tags at the ringing site |
| 2021 | 35236 | 7351 | 17_Fedderwardersiel | all | numerous false positives of various tags from several countries |
| 2021 | 35249 | 4892 | Huiberts | after 14-9-2021 00:00 | detections of a dropped off tag after 2021-09-13 |
| 2021 | 35252 | 4711 | Noorderhaven Julianadorp | after 16-9-2021 00:00 | detections of a dropped off tag after 2021-09-15 |
| 2021 | 35259 | 7351 | 17_Fedderwardersiel | all | numerous false positives of various tags from several countries |
| 2021 | 35269 | 4711 | Noorderhaven Julianadorp | after 12-9-2021 00:00 | detections of a dropped off tag after 2021-09-11 |
| 2021 | 35567 | 7709 | Safari | all | unlikely, detections in the same period in the tagging area |
| 2021 | 35568 | 7668 | Kennemerwind | after 25-9-2021 00:00 | detections of a dropped off tag after 2021-09-24 |
| 2021 | 35572 | 7692 | Norderney 3 | all | unlikely, probably caused by the presense of various tagged northern wheatears |
| 2021 | 36044 | 4892 | Huiberts | after 28-9-2021 00:00 | detections of a dropped off tag after 2021-09-27 |
| 2021 | 36052 | 7351 | 17_Fedderwardersiel | all | numerous false positives of various tags from several countries |
| 2021 | 36052 | 7351 | 17_Fedderwardersiel | all | numerous false positives of various tags from several countries |
| 2022 | 36331 | 8886 | Borkum 1 | all | numerous false positives of various tags from several countries from 1/10/2022 onwards |
| 2022 | 42200 | 8886 | Borkum 1 | all | numerous false positives of various tags from several countries from 1/10/2022 onwards |
| 2022 | 42201 | 7230 | Vogelzand | after 2-9-2022 05:00 | detections of a dropped off tag after 2022-9-2 05:00 |
| 2022 | 42205 | 8886 | Borkum 1 | all | numerous false positives of various tags from several countries from 1/10/2022 onwards |
| 2022 | 42212 | 4711 | Noorderhaven Julianadorp | after 15-9-2021 00:00 | detections of a dropped off tag after 2022-09-14 |
| 2022 | 42214 | 7351 | 17_Fedderwardersiel | all | numerous false positives of various tags from several countries |
| 2022 | 42214 | 7315 | Vlieland Kazerne | all | Likely caused by tagged songbirds present al this location (21/10/2022) |
| 2022 | 42216 | 7315 | Vlieland Kazerne | all | unlikely (4/11), caused by tagged songbirds present al this location |
| 2022 | 42217 | 7351 | 17_Fedderwardersiel | all | numerous false positives of various tags from several countries |
| 2022 | 42222 | 7709 | Safari | all | same day detections in Noord Holland |
| 2022 | 42222 | 7709 | Safari | all | same day detections in Noord Holland |
| 2022 | 42224 | 8886 | Borkum 1 | all | numerous false positives of various tags from several countries from 1/10/2022 onwards |
| 2022 | 42234 | 8886 | Borkum 1 | all | numerous false positives of various tags from several countries from 1/10/2022 onwards |

*Tabe S2-1: false positives (continued)*

| Year | Tag Deployment | Receiver Deployment | Receiver | Date | Comments |
|---|---|---|---|---|---|
| 2022 | 42252 | 7315 | Vlieland Kazerne | all | during daylight hours (19/10), caused by tagged songbirds present al this location |
| 2022 | 42256 | 8886 | Borkum 1 | all | numerous false positives of various tags from several countries from 1/10/2022 onwards |
| 2022 | 42256 | 9047 | Caister-on-Sea Lifeboat | all | numerous false positives of various tags from several countries from 30/11/2022 onwards |
| 2022 | 42258 | 8886 | Borkum 1 | all | numerous false positives of various tags from several countries from 1/10/2022 onwards |
| 2022 | 42259 | 8886 | Borkum 1 | all | numerous false positives of various tags from several countries from 1/10/2022 onwards |
| 2022 | 42261 | 8886 | Borkum 1 | all | numerous false positives of various tags from several countries from 1/10/2022 onwards |
| 2022 | 42261 | 7351 | 17_Fedderwardersiel | all | numerous false positives of various tags from several countries |
| 2022 | 42263 | 8886 | Borkum 1 | all | numerous false positives of various tags from several countries from 1/10/2022 onwards |
| 2022 | 42264 | 8886 | Borkum 1 | all | numerous false positives of various tags from several countries from 1/10/2022 onwards |
| 2022 | 42295 | 4892 | Huiberts | after 11-11-2022 00:00 | detections of a dropped off tag after 2022-11-10 |
| 2022 | 42303 | 8886 | Borkum 1 | all | numerous false positives of various tags from several countries from 1/10/2022 onwards |
| 2022 | 42308 | 4892 | Huiberts | after 26-9-2022 05:00 | detections of a dropped off tag after 2022-09-26 05:00 |
| 2022 | 42311 | 7351 | 17_Fedderwardersiel | all | numerous false positives of various tags from several countries |
| 2022 | 42312 | 4892 | Huiberts | after 14-9-2022 06:00 | detections of a dropped off tag after 2022-09-14 06:00 |
| 2022 | 42315 | 8886 | Borkum 1 | all | numerous false positives of various tags from several countries from 1/10/2022 onwards |
| 2022 | 42315 | 4892 | Huiberts | after 24-9-2022 00:00 | detections of a dropped off tag after 2022-09-24 00:00 |
| 2022 | 42316 | 8886 | Borkum 1 | all | numerous false positives of various tags from several countries from 1/10/2022 onwards |
| 2022 | 42317 | 8886 | Borkum 1 | all | numerous false positives of various tags from several countries from 1/10/2022 onwards |
| 2022 | 42317 | 7351 | 17_Fedderwardersiel | all | numerous false positives of various tags from several countries |
| 2022 | 42320 | 8886 | Borkum 1 | all | numerous false positives of various tags from several countries from 1/10/2022 onwards |
| 2022 | 42320 | 7351 | 17_Fedderwardersiel | all | numerous false positives of various tags from several countries |
| 2022 | 42323 | 8886 | Borkum 1 | all | numerous false positives of various tags from several countries from 1/10/2022 onwards |
| 2022 | 42326 | 8886 | Borkum 1 | all | numerous false positives of various tags from several countries from 1/10/2022 onwards |
| 2022 | 42326 | 7351 | 17_Fedderwardersiel | all | numerous false positives of various tags from several countries |
| 2022 | 42326 | 8886 | Borkum 1 | all | numerous false positives of various tags from several countries from 1/10/2022 onwards |
| 2022 | 42326 | 7351 | 17_Fedderwardersiel | all | numerous false positives of various tags from several countries |
| 2022 | 42407 | 7351 | 17_Fedderwardersiel | all | numerous false positives of various tags from several countries |
| 2022 | 42410 | 7351 | 17_Fedderwardersiel | all | numerous false positives of various tags from several countries |
| 2022 | 42426 | 8886 | Borkum 1 | all | numerous false positives of various tags from several countries from 1/10/2022 onwards |
| 2022 | 42432 | 8886 | Borkum 1 | all | numerous false positives of various tags from several countries from 1/10/2022 onwards |
| 2022 | 42434 | 8886 | Borkum 1 | all | numerous false positives of various tags from several countries from 1/10/2022 onwards |

### References


Birds Canada, 2022. Fetch and use data from the Motus Wildlife Tracking System.
