## Appendix S3 for "Stopover departure decisions and movement patterns of migratory bats during autumn migration"

### Appendix S3: Arm length, Weight and Body Mass Index (BMI)

Females were slightly larger than males, as indicated by arm length, but no differences were observed among age classes within each sex (Table S3-1 and Figure S3-1). Adult bats were heavier than first-year bats within each sex. Among all groups, adult females were the heaviest, while first-year females and adult males had similar body weights (Table S3-2 and Figure S3-2). Calculating the residuals of the regression of Weight on Arm length per SexAge category (Figure S3-3) resulted in the BMI, which did not differ among the SexAge categories, though the variation was a bit higher in adult females (Table S3-3 and Figure S3-4).

*Table S3-1 : Armlength per SexAge category*

| Category | Number of individuals | Arm length [mm] | | | | |
|---|---|---|---|---|---|---|
| | | median | mean | sd | min | max |
| F_adult | 202 | 34.6 | 34.7 | 0.9 | 32.2 | 37.0 |
| F_first | 102 | 34.5 | 34.4 | 0.8 | 32.2 | 36.7 |
| M_adult | 167 | 33.6 | 33.6 | 0.8 | 31.7 | 35.1 |
| M_first | 90 | 33.6 | 33.7 | 0.9 | 31.8 | 36.8 |
| All bats | 561 | 34.2 | 34.1 | 1.0 | 31.7 | 37.0 |

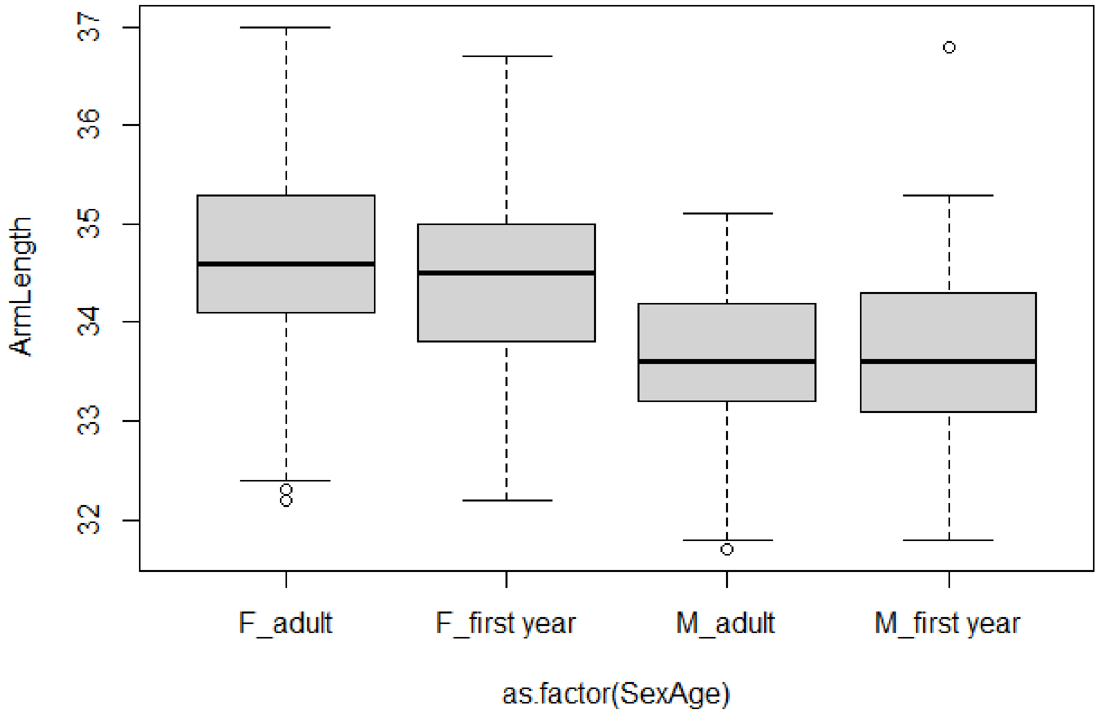


*Figure S3-1: Boxplot of armlength per SexAge category*

*Table S3-2 : Weight per SexAge category*

| Category | Number of individuals | Weight [g] | | | | |
|---|---|---|---|---|---|---|
| | | median | mean | sd | min | max |
| F_adult | 202 | 8.8 | 8.9 | 1.1 | 6.2 | 12.2 |
| F_first year | 102 | 8.1 | 8.2 | 0.8 | 6.2 | 10.1 |
| M_adult | 167 | 8.1 | 8.1 | 0.9 | 5.2 | 10.8 |
| M_first year | 90 | 7.8 | 7.9 | 0.8 | 6.0 | 10.0 |
| All bats | 561 | 8.2 | 8.4 | 1.0 | 5.2 | 12.2 |

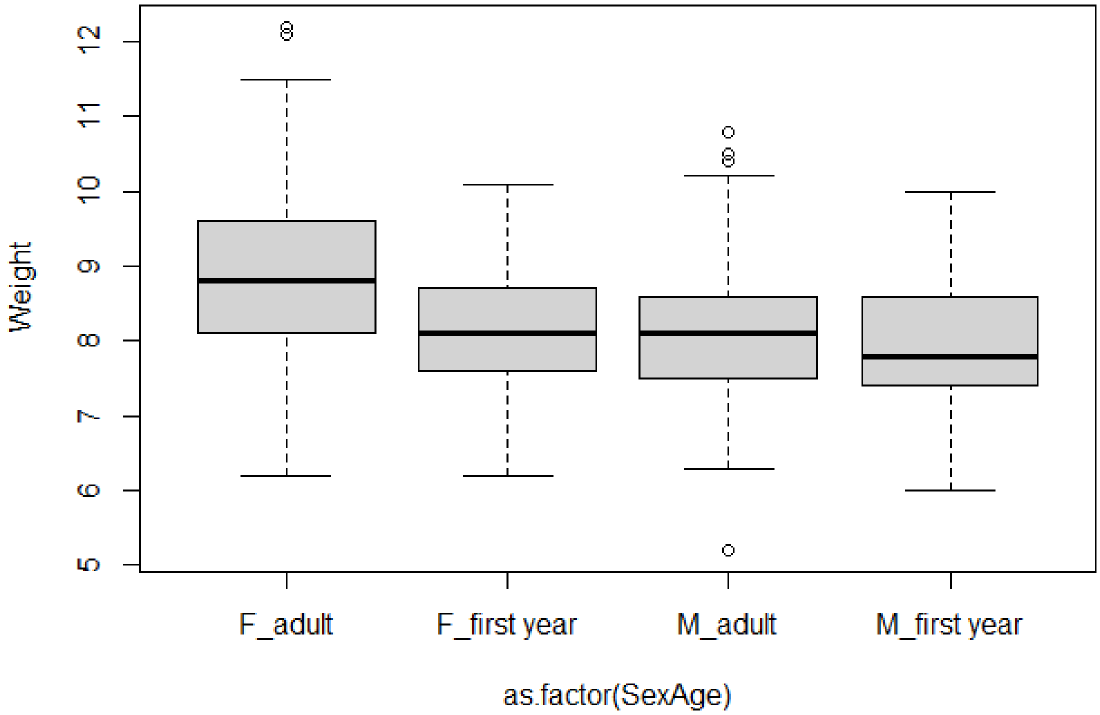


*Figure S3-2 Boxplot of weights per SexAge category*

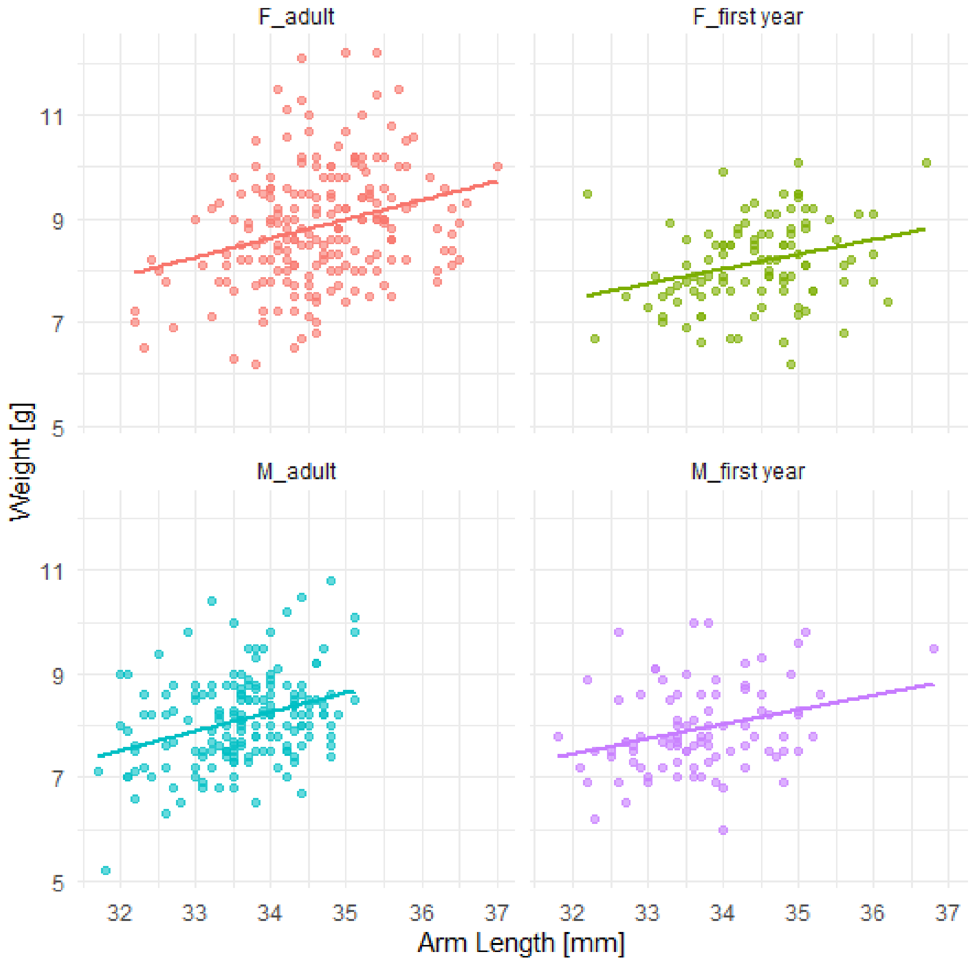


*Figure S3-3: Regression of Weight on Arm length per SexAge category*

*Table S3-3 : BMI per SexAge category*

| Category | Number of individuals | Body Mass Index | | | | |
|---|---|---|---|---|---|---|
| | | median | mean | sd | min | max |
| F_adult | 202 | -0.12 | -3.54E-18 | 1.09 | -2.35 | 3.33 |
| F_first year | 102 | -0.04 | -2.44E-17 | 0.80 | -2.08 | 1.99 |
| M_adult | 167 | -0.06 | 1.78E-17 | 0.82 | -2.24 | 2.44 |
| M_first year | 90 | -0.10 | -1.17E-17 | 0.81 | -2.02 | 2.18 |
| All bats | 561 | -0.09 | -2.43E-18 | 0.92 | -2.35 | 3.33 |

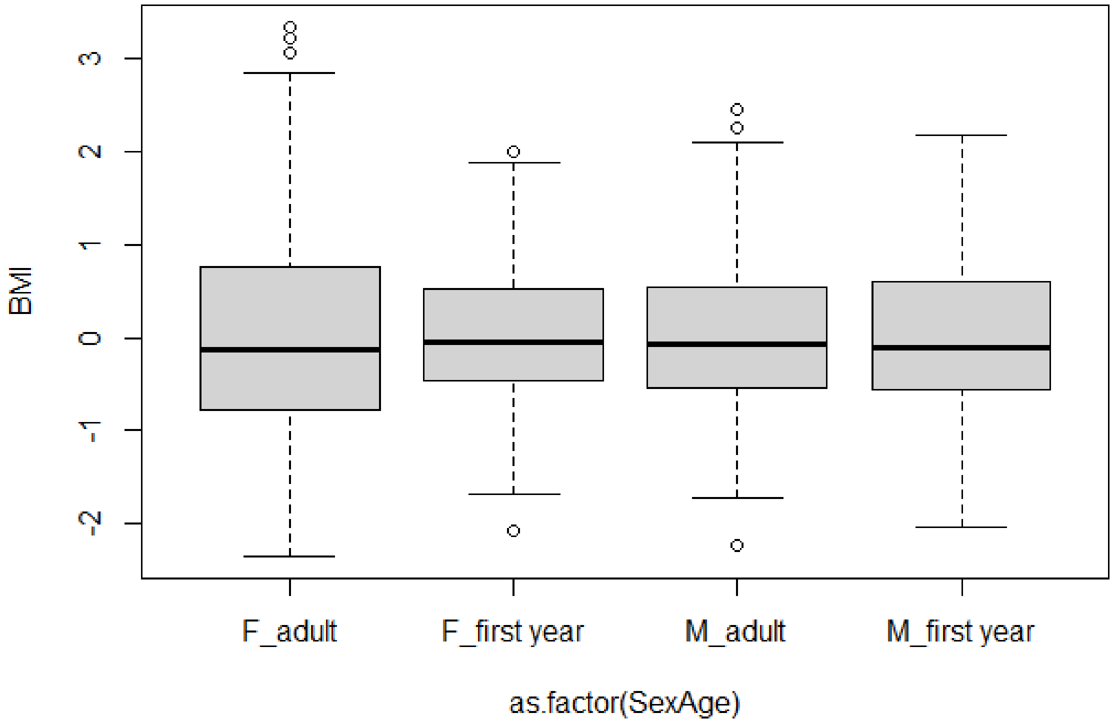


*Figure SF3-4 : Body Mass Index per SexAge category*
