## Appendix S4 for "Stopover departure decisions and movement patterns of migratory bats during autumn migration"

**Appendix S4: flights of all individuals**

These figures shows the observed flights of individual bats over 30 km, with arrows indicating flight direction. Flights occurring within a single night are shown in blue, while movements spanning multiple nights are shown in orange. Each flight is annotated with start and end timestamps, along with the timestamps of tagging and last detection. MOTUS receivers are indicated by yellow dots.

### Deployment ID 18816 (adult F)

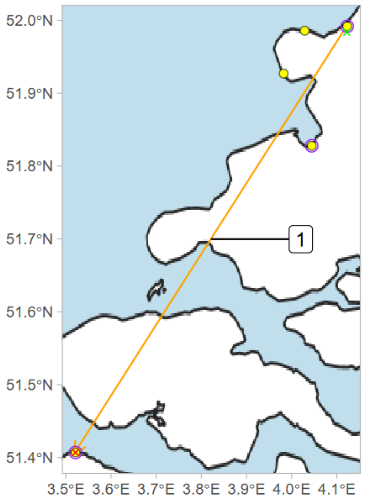


release: 2018-09-04 15:00
1) 2018-09-04 19:05 - 2018-09-05 21:11
final detection: 2018-09-05 21:12

### Deployment ID 18817 (adult F)

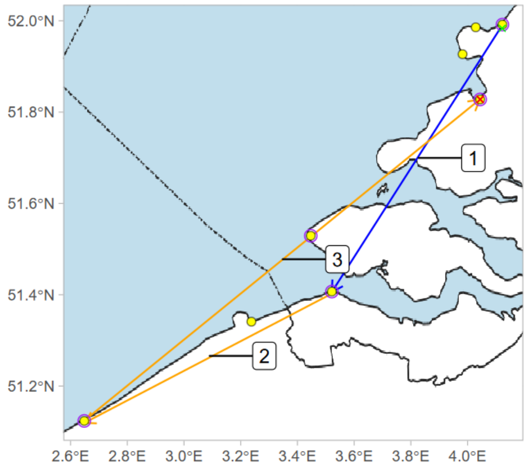


release: 2018-09-04 15:00
1) 2018-09-04 19:01 - 2018-09-04 22:25
2) 2018-09-04 22:25 - 2018-09-08 22:14
3) 2018-09-08 22:14 - 2018-11-18 00:54
final detection: 2018-11-18 00:54

### Deployment ID 18822 (first year F)

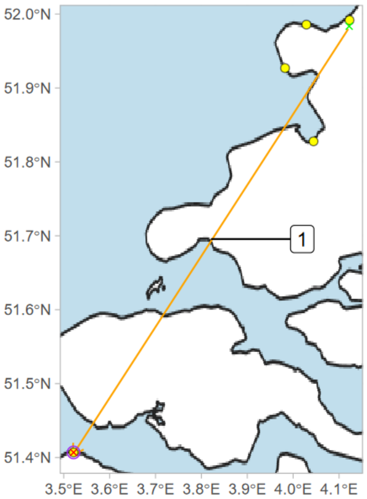


release: 2018-09-13 10:00
1) 2018-09-13 18:03 - 2018-09-16 22:35
final detection: 2018-09-16 22:36

### Deployment ID 18824 (adult F)

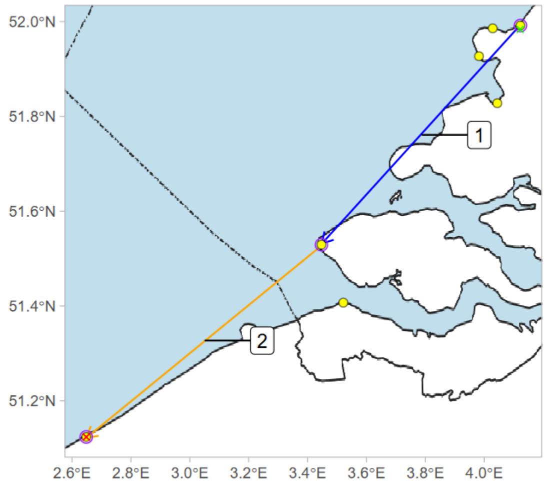


release: 2018-09-13 10:00
1) 2018-09-13 18:43 - 2018-09-13 22:16
2) 2018-09-13 22:16 - 2018-09-14 20:49
final detection: 2018-09-14 20:51

### Deployment ID 18825 (adult F)

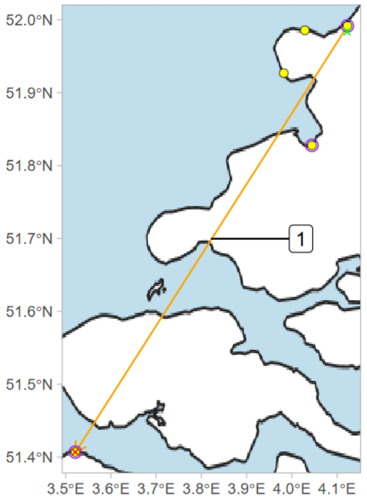


release: 2018-09-08 10:00
1) 2018-09-09 18:52 - 2018-09-17 23:23
final detection: 2018-09-17 23:25

### Deployment ID 18828 (adult F)

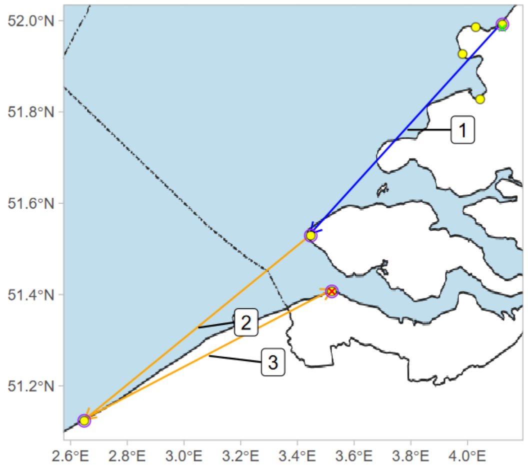


release: 2018-09-13 10:00
1) 2018-09-13 18:38 - 2018-09-13 21:59
2) 2018-09-13 21:59 - 2018-09-14 20:13
3) 2018-09-14 20:13 - 2018-09-19 00:28
final detection: 2018-09-19 18:32

Deployment ID 18829 (adult F)

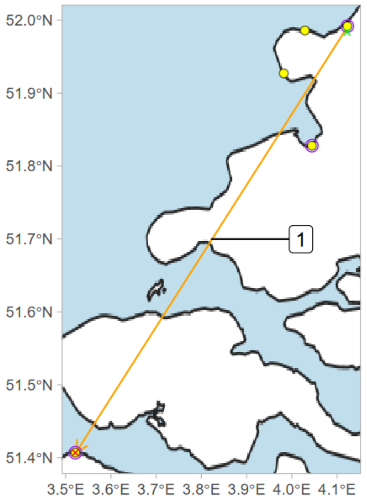


release: 2018-09-04 15:00
1) 2018-09-04 19:51 - 2018-09-05 21:33
final detection: 2018-09-05 21:38

Deployment ID 18831 (first year F)

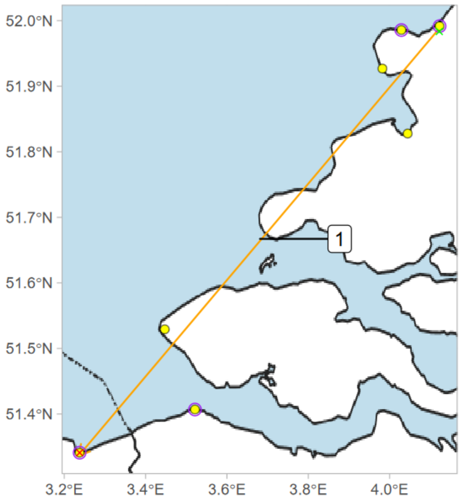


release: 2018-09-04 15:00
1) 2018-09-04 19:06 - 2018-09-05 20:34
final detection: 2018-09-05 20:35

Deployment ID 18832 (first year F)

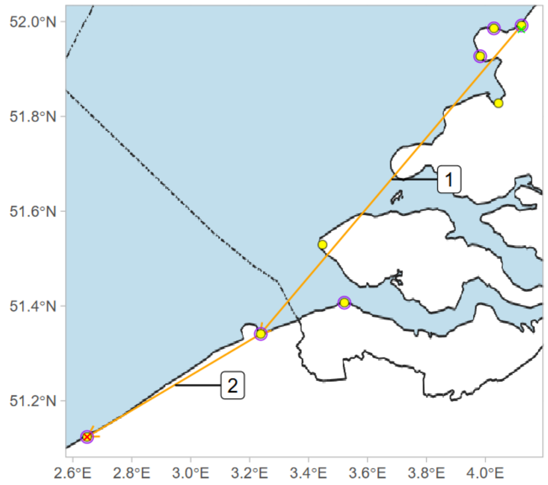


release: 2018-09-04 15:00
1) 2018-09-04 19:12 - 2018-09-05 20:04
2) 2018-09-05 20:04 - 2018-09-06 20:34
final detection: 2018-09-06 20:37

Deployment ID 18834 (first year F)

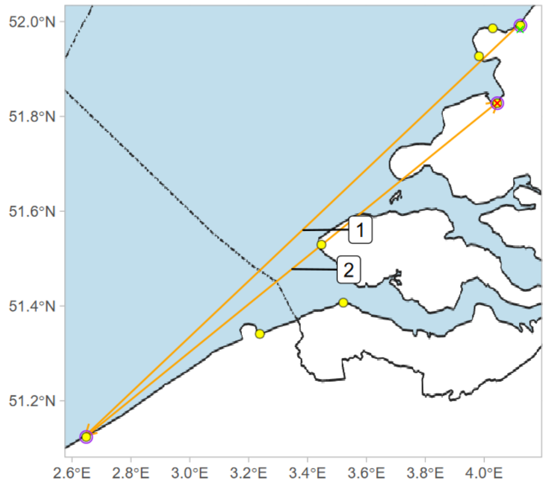


release: 2018-09-13 10:00
1) 2018-09-13 18:55 - 2018-09-14 22:05
2) 2018-09-14 22:05 - 2018-09-19 22:56
final detection: 2018-09-19 22:59

Deployment ID 18842 (adult F)

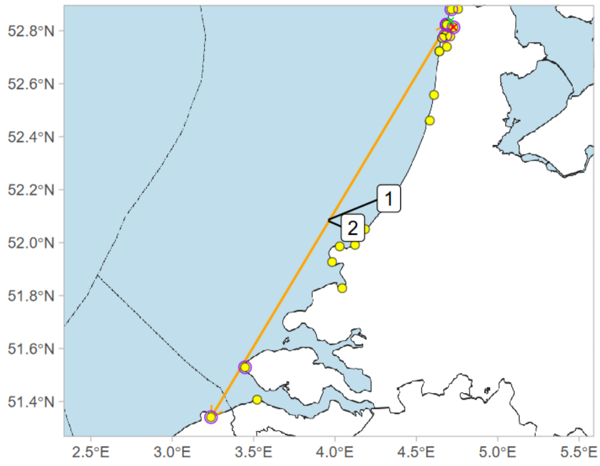


release: 2018-08-30 16:00
1) 2018-08-30 19:22 - 2018-09-01 00:59
2) 2018-09-01 00:59 - 2018-09-22 22:29
final detection: 2018-09-24 20:32

Deployment ID 18848 (first year F)

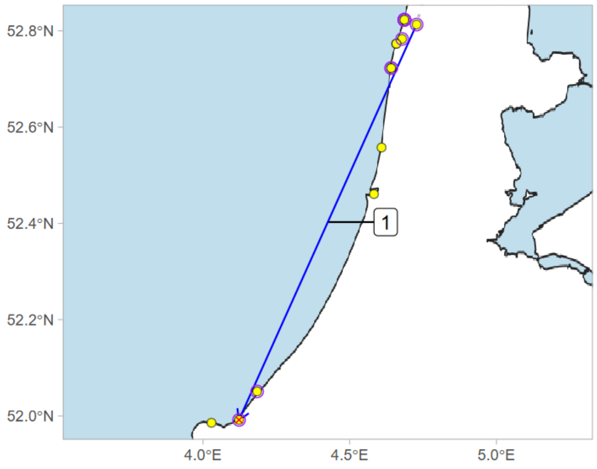


release: 2018-08-23 12:00
1) 2018-08-30 20:35 - 2018-08-30 23:47
final detection: 2018-08-31 00:10

Deployment ID 18852 (adult F)

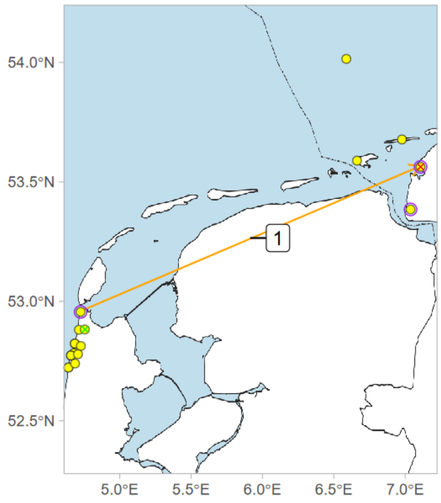


release: 2018-09-28 09:00
1) 2018-09-28 19:32 - 2018-09-29 23:19
final detection: 2018-09-29 23:59

Deployment ID 18859 (first year M)

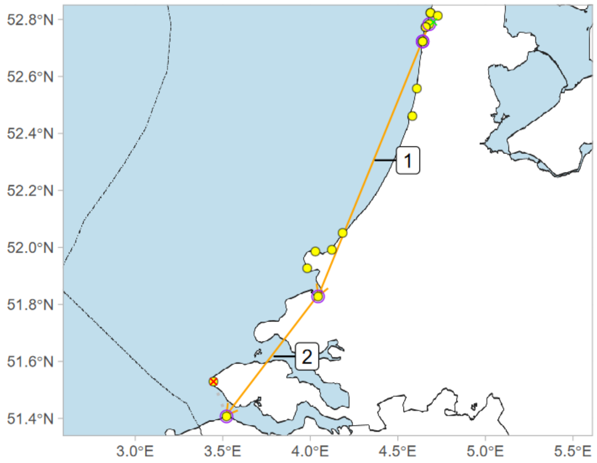


release: 2018-08-23 06:00
1) 2018-08-23 19:42 - 2018-08-28 03:51
2) 2018-08-28 03:51 - 2018-08-28 21:45
final detection: 2018-08-31 00:01

Deployment ID 19016 (adult F)

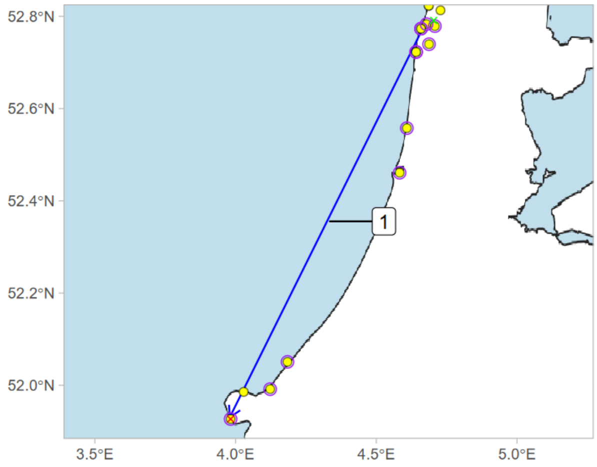


release: 2018-09-04 10:00
1) 2018-09-28 18:12 - 2018-09-28 20:15
final detection: 2018-09-28 20:18

Deployment ID 19021 (adult F)

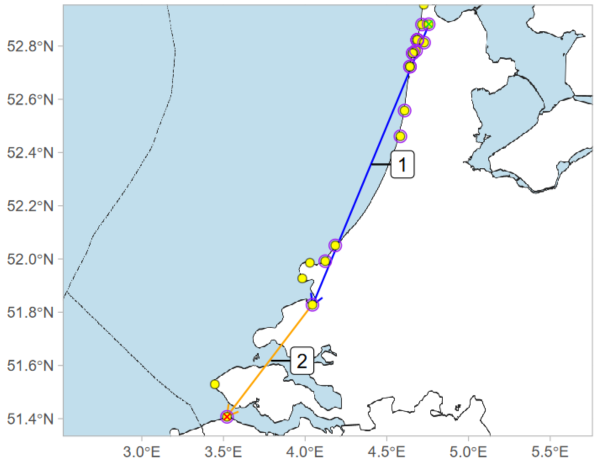


release: 2018-09-04 08:00
1) 2018-09-04 20:00 - 2018-09-04 23:36
2) 2018-09-04 23:36 - 2018-09-05 21:36
final detection: 2018-09-05 21:57

Deployment ID 19028 (adult F)

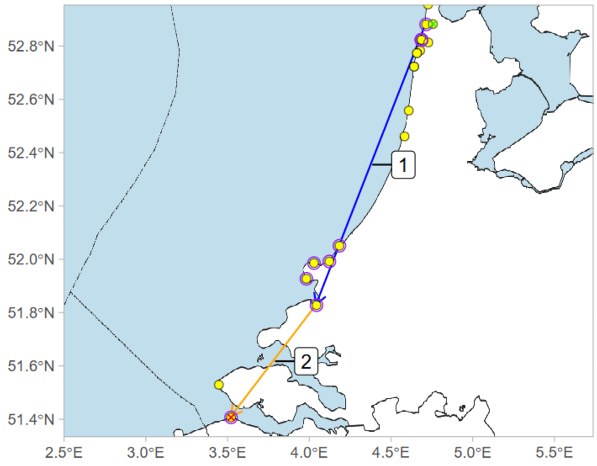


release: 2018-09-04 08:00
1) 2018-09-04 19:15 - 2018-09-04 23:19
2) 2018-09-04 23:19 - 2018-09-05 20:10
final detection: 2018-09-05 20:19

Deployment ID 19090 (adult F)

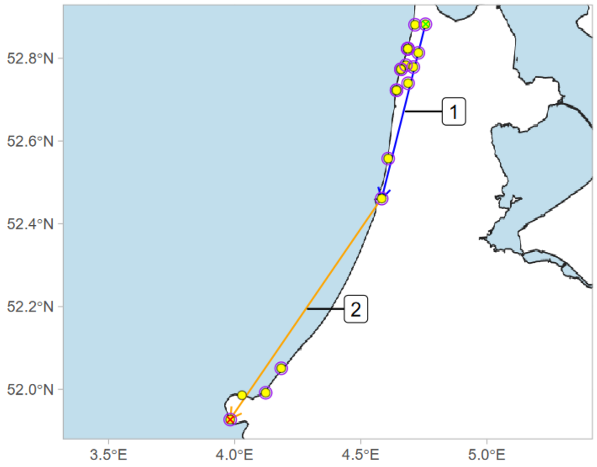


release: 2018-09-08 10:00
1) 2018-09-28 00:35 - 2018-09-28 01:45
2) 2018-09-28 01:45 - 2018-09-28 18:50
final detection: 2018-09-28 18:55

Deployment ID 19092 (adult F)

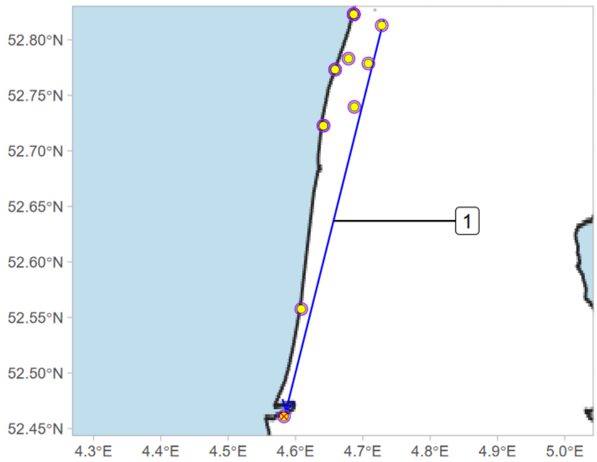


release: 2018-09-13 10:00
1) 2018-09-28 18:47 - 2018-09-28 19:40
final detection: 2018-09-28 19:43

Deployment ID 19094 (adult F)

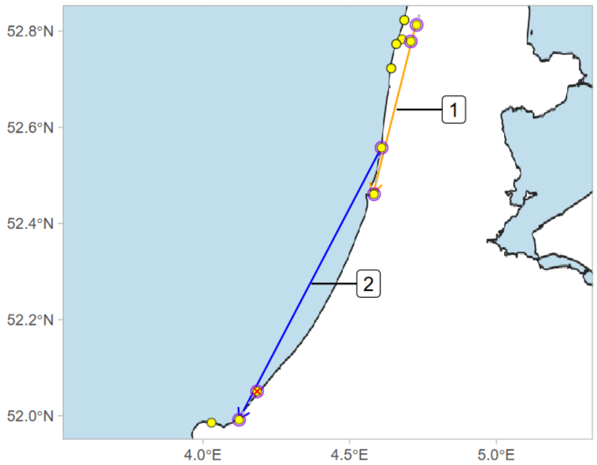


release: 2018-09-08 10:00
1) 2018-09-15 23:51 - 2018-09-16 23:08
2) 2018-09-17 22:03 - 2018-09-18 01:43
final detection: 2018-09-18 01:49

Deployment ID 19100 (adult F)

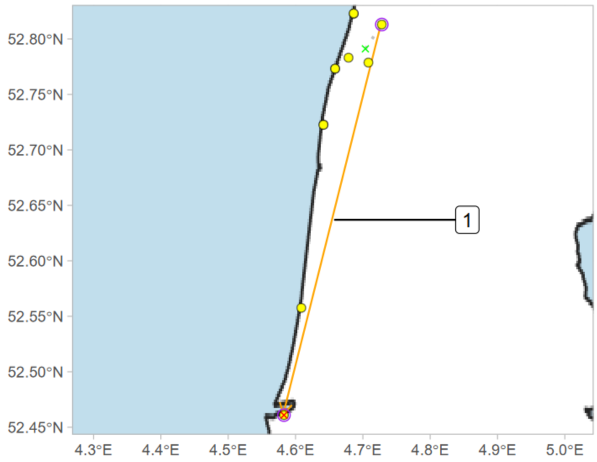


release: 2018-09-08 10:00
1) 2018-09-08 19:03 - 2018-09-19 00:43
final detection: 2018-09-19 00:44

Deployment ID 19110 (first year F)

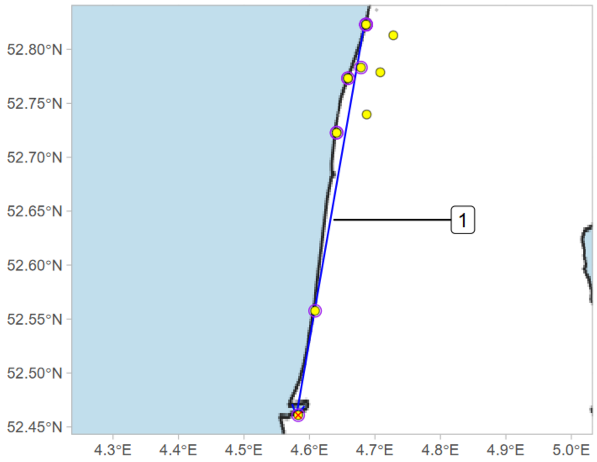


release: 2018-09-08 10:00
1) 2018-10-10 18:14 - 2018-10-10 20:15
final detection: 2018-10-10 20:19

Deployment ID 19119 (adult M)

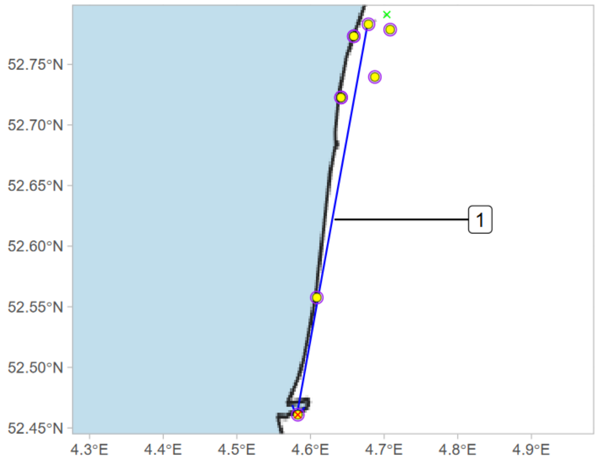


release: 2018-09-28 09:00
1) 2018-10-07 18:16 - 2018-10-07 20:05
final detection: 2018-10-07 20:10

Deployment ID 19125 (adult F)

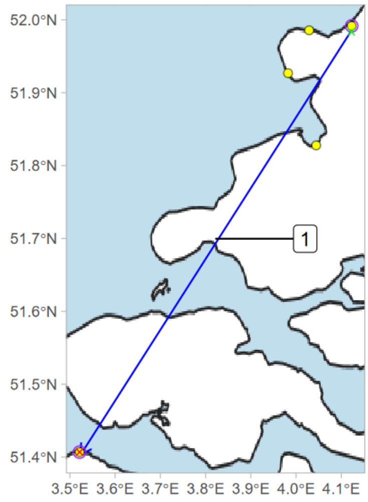


release: 2018-09-13 10:00
1) 2018-09-13 18:37 - 2018-09-13 22:50
final detection: 2018-09-13 22:51

Deployment ID 19126 (first year F)

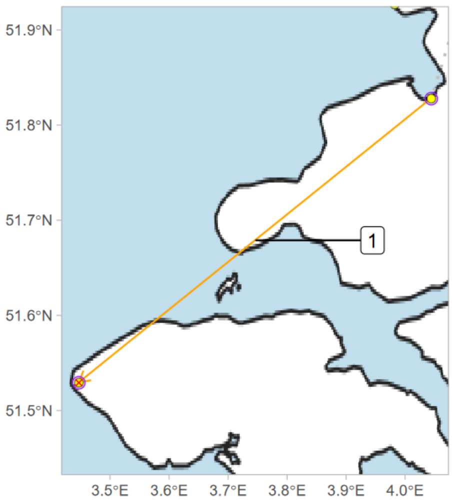


release: 2018-09-08 10:00
1) 2018-09-09 20:04 - 2018-09-12 20:39
final detection: 2018-09-12 20:45

Deployment ID 19129 (adult F)

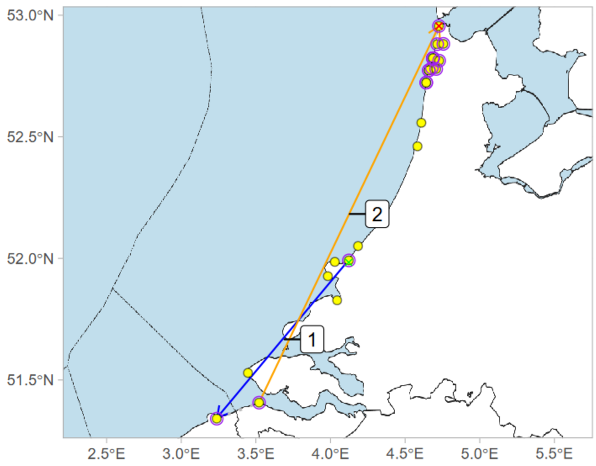


release: 2018-09-13 10:00
1) 2018-09-13 18:29 - 2018-09-13 23:19
2) 2018-09-14 22:21 - 2018-09-17 21:24
final detection: 2018-09-17 21:43

Deployment ID 19132 (adult F)

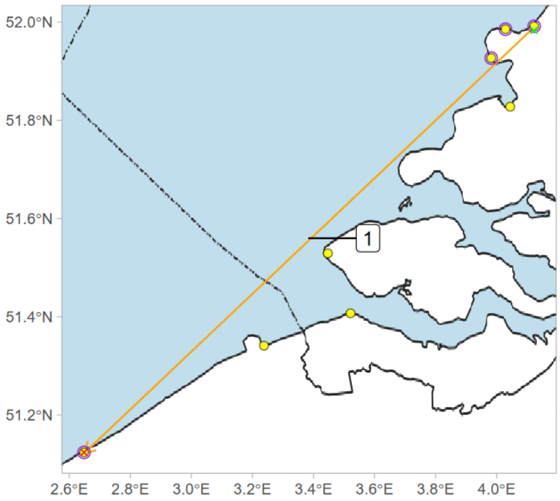


release: 2018-09-13 10:00
1) 2018-09-13 18:46 - 2018-09-14 20:26
final detection: 2018-09-14 20:28

Deployment ID 19133 (adult F)

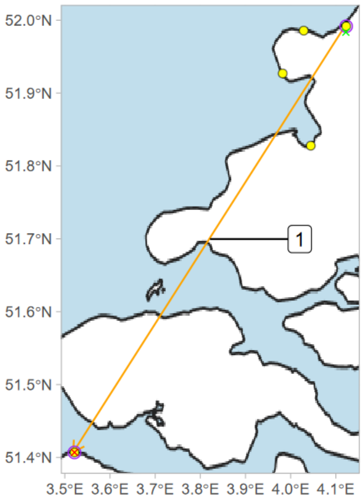


release: 2018-09-13 10:00
1) 2018-09-13 18:56 - 2018-09-14 22:23
final detection: 2018-09-14 22:25

Deployment ID 19138 (adult F)

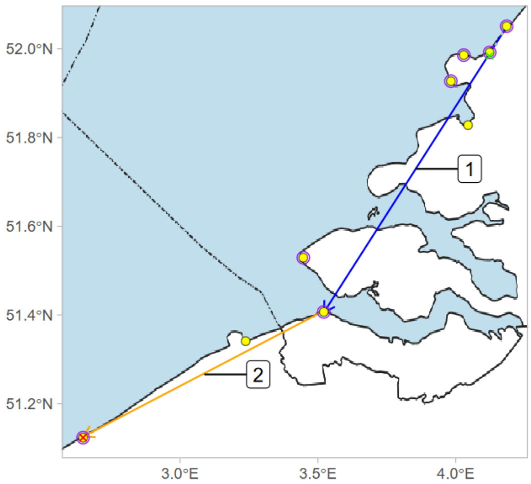


release: 2018-09-08 10:00
1) 2018-09-12 20:18 - 2018-09-12 22:39
2) 2018-09-12 22:39 - 2018-09-13 20:19
final detection: 2018-09-13 20:23

Deployment ID 19468 (adult F)

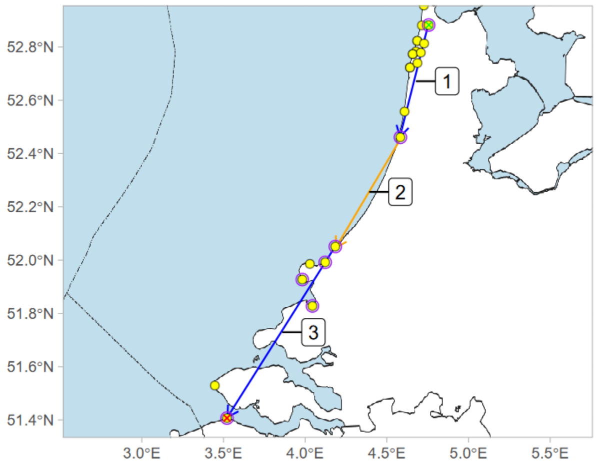


release: 2018-09-22 09:00
1) 2018-10-05 18:16 - 2018-10-05 21:06
2) 2018-10-05 22:27 - 2018-10-07 17:47
3) 2018-10-07 18:01 - 2018-10-07 21:20
final detection: 2018-10-07 21:37

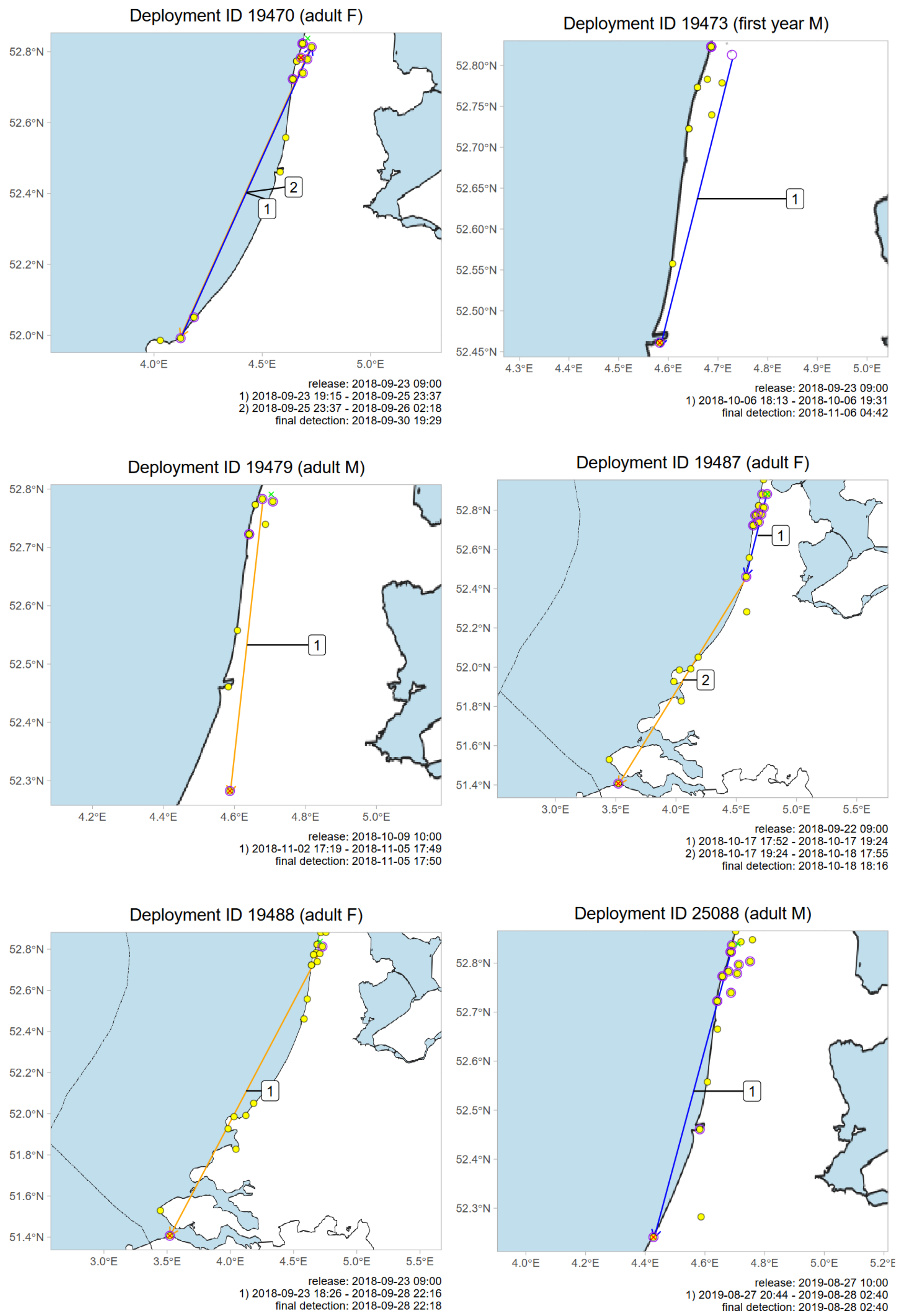
Deployment ID 19470 (adult F)
52.8°N
52.6°N
52.4°N
52.2°N
52.0°N
4.0°E
4.5°E
5.0°E
1
2
release: 2018-09-23 09:00
1) 2018-09-23 19:15 - 2018-09-25 23:37
2) 2018-09-25 23:37 - 2018-09-26 02:18
final detection: 2018-09-30 19:29
Deployment ID 19473 (first year M)
52.80°N
52.75°N
52.70°N
52.65°N
52.60°N
52.55°N
52.50°N
52.45°N
4.3°E
4.4°E
4.5°E
4.6°E
4.7°E
4.8°E
4.9°E
5.0°E
1
release: 2018-09-23 09:00
1) 2018-10-06 18:13 - 2018-10-06 19:31
final detection: 2018-11-06 04:42
Deployment ID 19479 (adult M)
52.8°N
52.7°N
52.6°N
52.5°N
52.4°N
52.3°N
4.2°E
4.4°E
4.6°E
4.8°E
5.0°E
1
release: 2018-10-09 10:00
1) 2018-11-02 17:19 - 2018-11-05 17:49
final detection: 2018-11-05 17:50
Deployment ID 19487 (adult F)
52.8°N
52.6°N
52.4°N
52.2°N
52.0°N
51.8°N
51.6°N
51.4°N
3.0°E
3.5°E
4.0°E
4.5°E
5.0°E
5.5°E
1
2
release: 2018-09-22 09:00
1) 2018-10-17 17:52 - 2018-10-17 19:24
2) 2018-10-17 19:24 - 2018-10-18 17:55
final detection: 2018-10-18 18:16
Deployment ID 19488 (adult F)
52.8°N
52.6°N
52.4°N
52.2°N
52.0°N
51.8°N
51.6°N
51.4°N
3.0°E
3.5°E
4.0°E
4.5°E
5.0°E
5.5°E
1
release: 2018-09-23 09:00
1) 2018-09-23 18:26 - 2018-09-28 22:16
final detection: 2018-09-28 22:18
Deployment ID 25088 (adult M)
52.8°N
52.7°N
52.6°N
52.5°N
52.4°N
52.3°N
4.0°E
4.2°E
4.4°E
4.6°E
4.8°E
5.0°E
5.2°E
1
release: 2019-08-27 10:00
1) 2019-08-27 20:44 - 2019-08-28 02:40
final detection: 2019-08-28 02:40

Deployment ID 25090 (adult F)

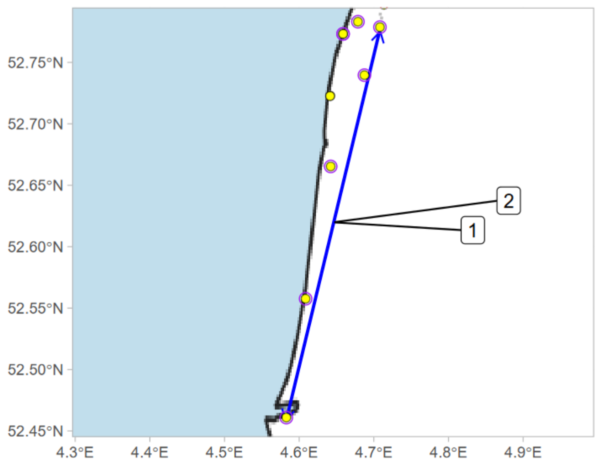


release: 2019-08-27 10:00
1) 2019-08-30 19:18 - 2019-08-30 21:54
2) 2019-08-30 23:34 - 2019-08-31 00:26
final detection: 2019-08-31 03:49

Deployment ID 25096 (first year F)

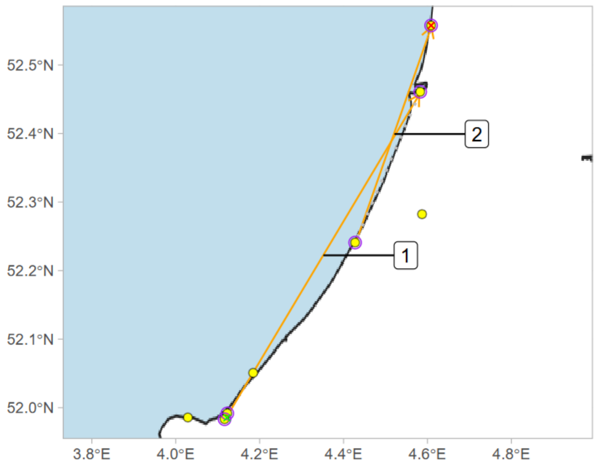


release: 2019-09-10 10:00
1) 2019-09-10 18:45 - 2019-09-13 20:33
2) 2019-09-13 21:22 - 2019-09-18 21:12
final detection: 2019-09-18 21:13

Deployment ID 25106 (adult F)

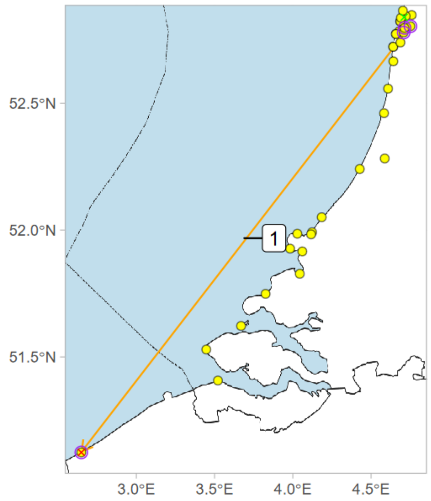


release: 2019-08-27 10:00
1) 2019-08-29 20:17 - 2019-09-09 23:36
final detection: 2019-09-09 23:36

Deployment ID 25115 (first year F)

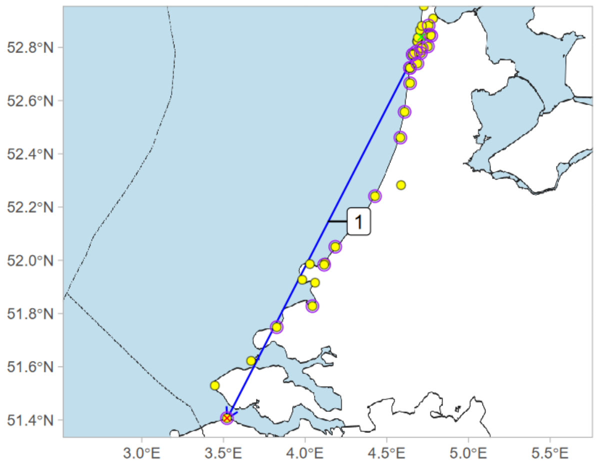


release: 2019-10-01 10:00
1) 2019-10-20 17:07 - 2019-10-20 21:19
final detection: 2019-10-20 21:32

Deployment ID 25118 (adult F)

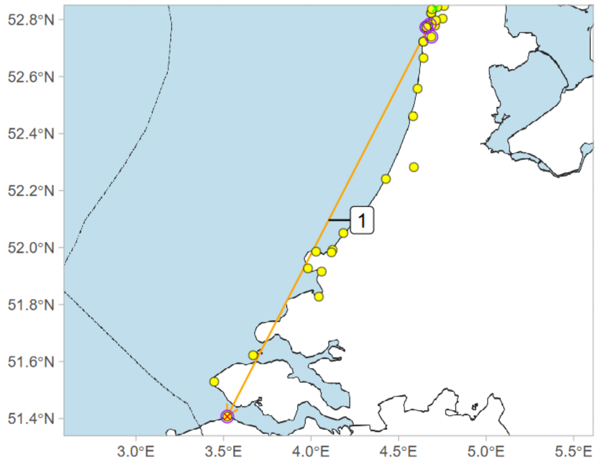


release: 2019-08-27 10:00
1) 2019-08-27 21:32 - 2019-09-04 21:29
final detection: 2019-09-04 21:31

Deployment ID 25119 (first year M)

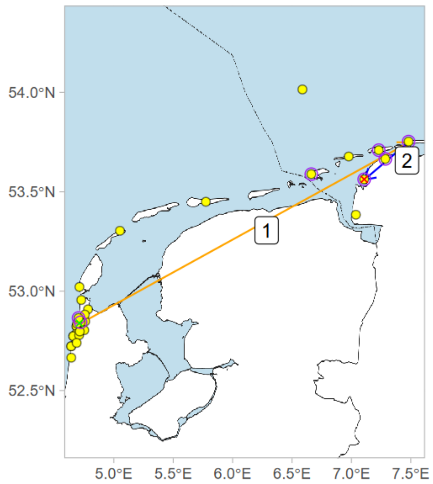


release: 2019-08-27 10:00
1) 2019-08-27 19:31 - 2019-08-30 01:22
2) 2019-08-30 19:13 - 2019-08-30 23:52
final detection: 2019-08-30 23:58

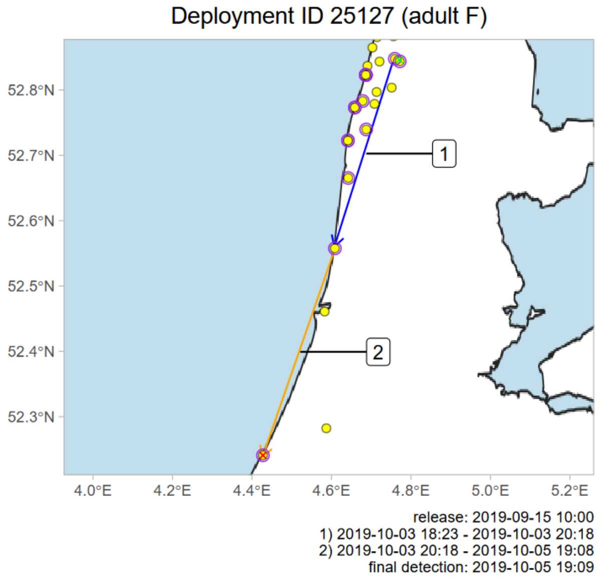
Deployment ID 25127 (adult F)
1
2
release: 2019-09-15 10:00
1) 2019-10-03 18:23 - 2019-10-03 20:18
2) 2019-10-03 20:18 - 2019-10-05 19:08
final detection: 2019-10-05 19:09

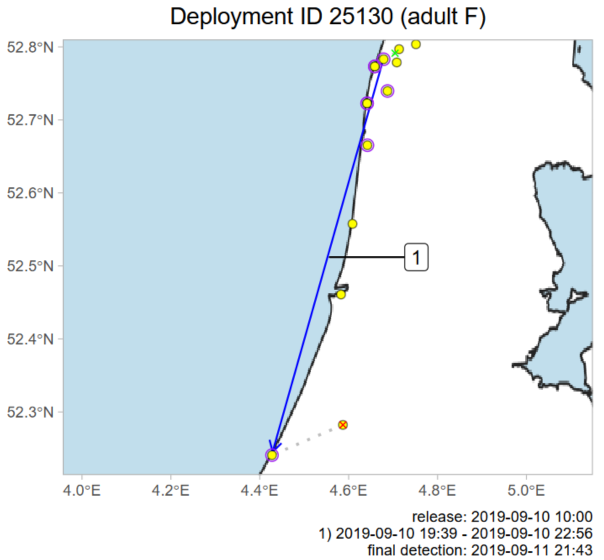
Deployment ID 25130 (adult F)
1
release: 2019-09-10 10:00
1) 2019-09-10 19:39 - 2019-09-10 22:56
final detection: 2019-09-11 21:43

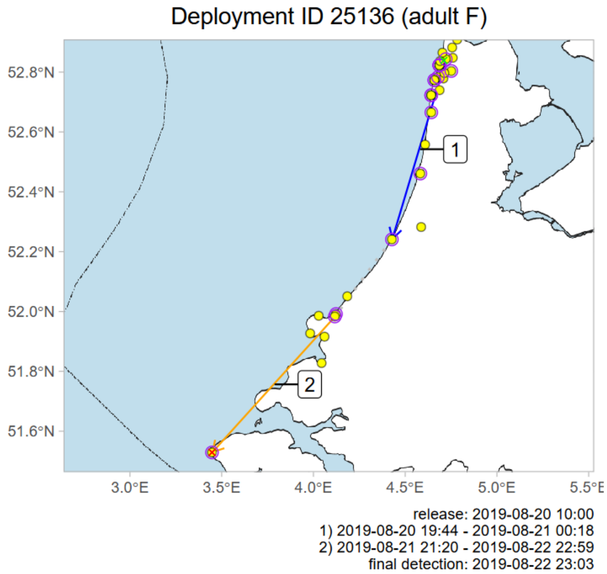
Deployment ID 25136 (adult F)
1
2
release: 2019-08-20 10:00
1) 2019-08-20 19:44 - 2019-08-21 00:18
2) 2019-08-21 21:20 - 2019-08-22 22:59
final detection: 2019-08-22 23:03

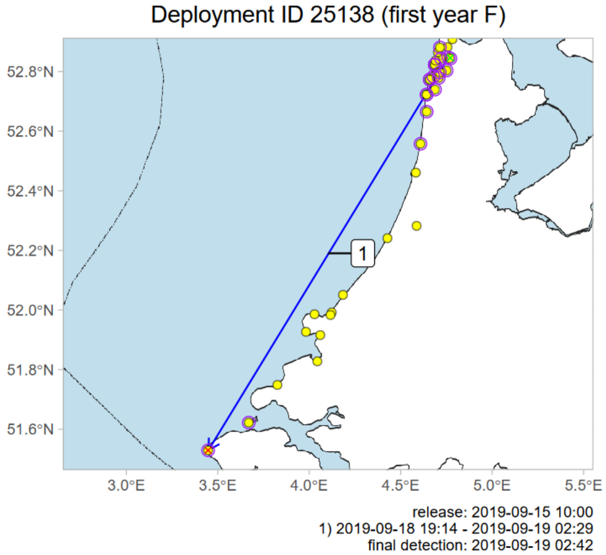
Deployment ID 25138 (first year F)
1
release: 2019-09-15 10:00
1) 2019-09-18 19:14 - 2019-09-19 02:29
final detection: 2019-09-19 02:42

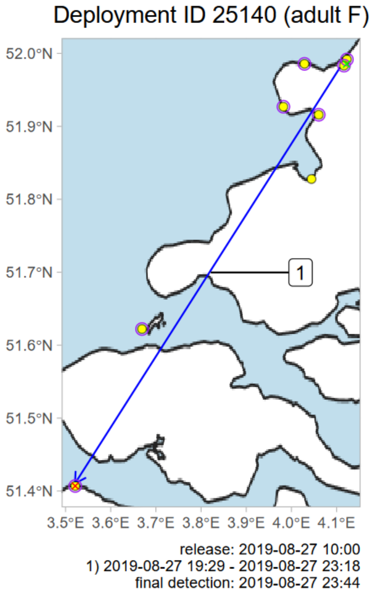
Deployment ID 25140 (adult F)
1
release: 2019-08-27 10:00
1) 2019-08-27 19:29 - 2019-08-27 23:18
final detection: 2019-08-27 23:44

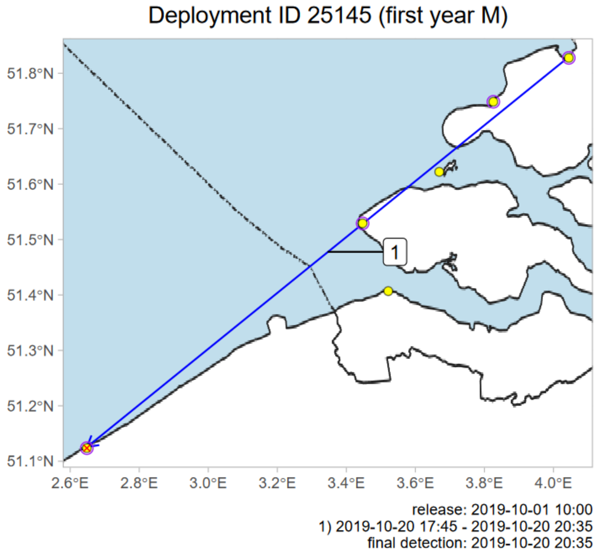
Deployment ID 25145 (first year M)
1
release: 2019-10-01 10:00
1) 2019-10-20 17:45 - 2019-10-20 20:35
final detection: 2019-10-20 20:35

Deployment ID 25146 (first year F)

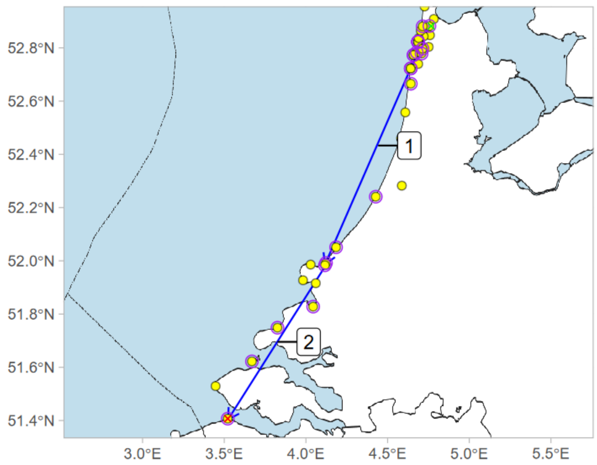


release: 2019-09-10 10:00
1) 2019-09-10 18:47 - 2019-09-11 01:04
2) 2019-09-13 18:59 - 2019-09-13 20:59
final detection: 2019-09-13 21:00

Deployment ID 25151 (adult F)

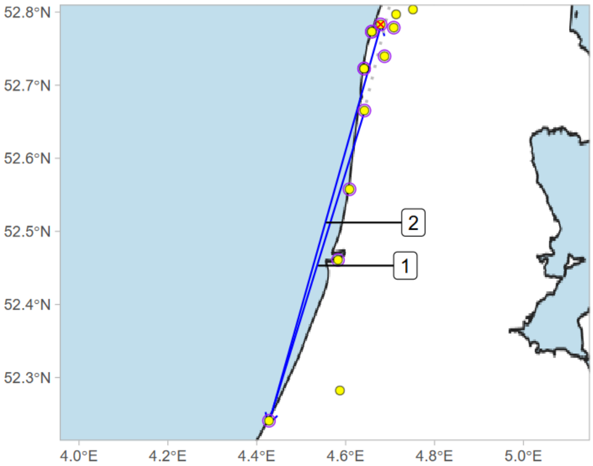


release: 2019-08-27 10:00
1) 2019-08-30 19:26 - 2019-08-30 23:33
2) 2019-08-30 23:43 - 2019-08-31 01:20
final detection: 2019-09-02 11:41

Deployment ID 25155 (adult F)

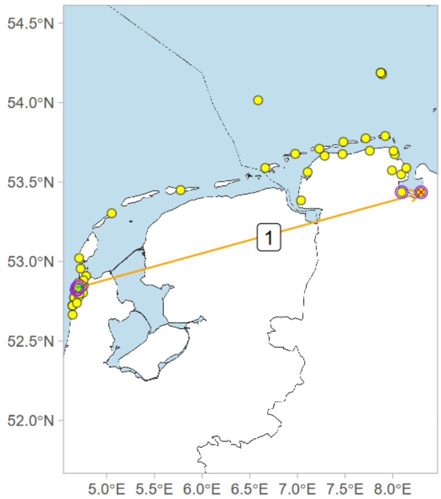


release: 2019-08-27 10:00
1) 2019-08-27 19:28 - 2019-08-30 01:19
final detection: 2019-08-30 01:19

Deployment ID 25170 (first year F)

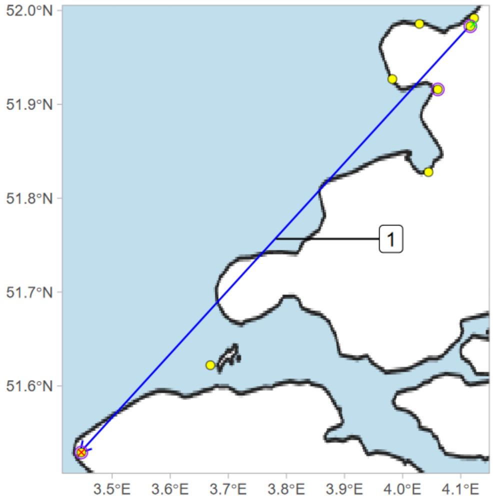


release: 2019-08-27 10:00
1) 2019-08-27 20:23 - 2019-08-28 00:41
final detection: 2019-08-28 01:02

Deployment ID 25180 (first year F)

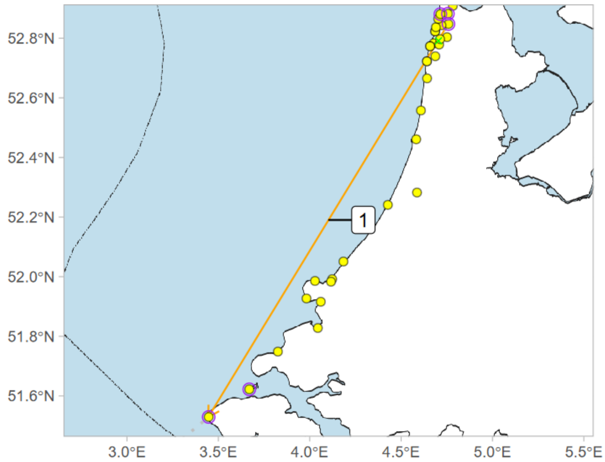


release: 2019-09-03 10:00
1) 2019-09-05 19:54 - 2019-09-09 21:57
final detection: 2019-09-10 21:45

Deployment ID 25221 (adult F)

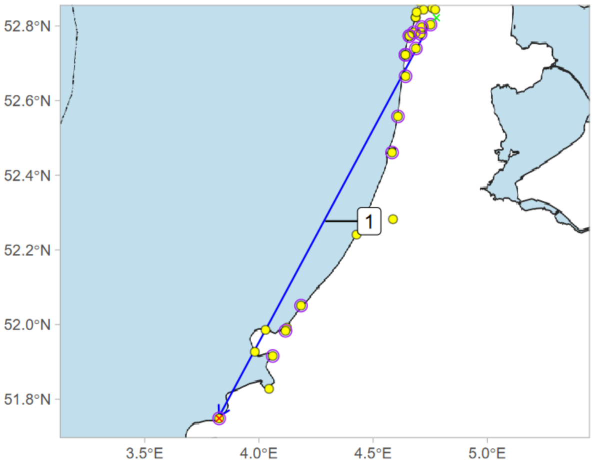


release: 2019-09-24 10:00
1) 2019-10-20 17:16 - 2019-10-20 20:48
final detection: 2019-10-20 20:58

Deployment ID 25226 (adult F)

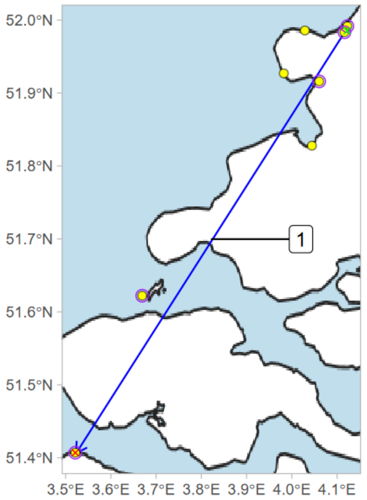


release: 2019-08-27 10:00
1) 2019-08-27 19:09 - 2019-08-27 23:47
final detection: 2019-08-27 23:53

Deployment ID 25958 (adult F)

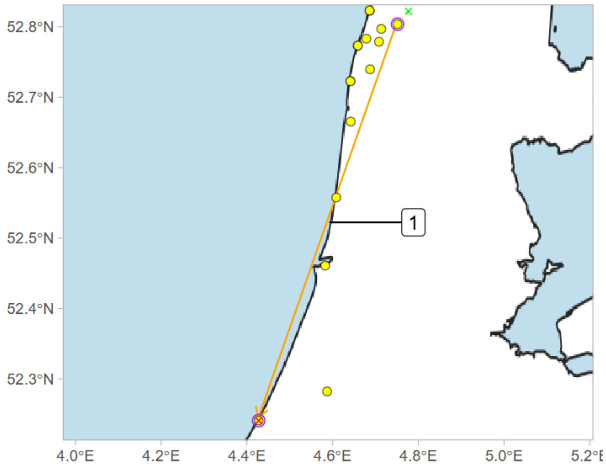


release: 2019-09-17 12:00
1) 2019-10-14 22:47 - 2019-10-24 01:26
final detection: 2019-10-24 01:29

Deployment ID 25968 (adult F)

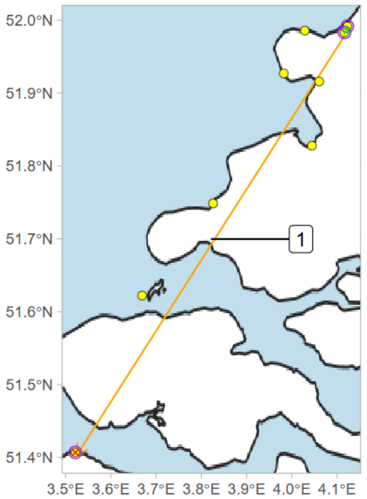


release: 2019-10-01 10:00
1) 2019-10-01 17:57 - 2019-10-05 20:44
final detection: 2019-10-05 20:44

Deployment ID 25970 (adult F)

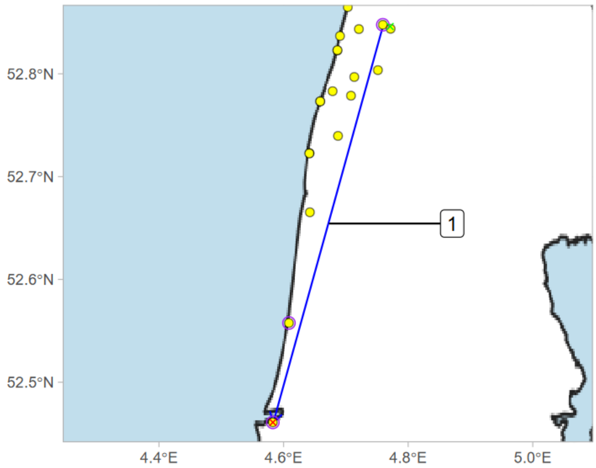


release: 2019-09-15 10:00
1) 2019-10-20 18:31 - 2019-10-20 20:57
final detection: 2019-10-20 20:58

Deployment ID 26140 (adult F)

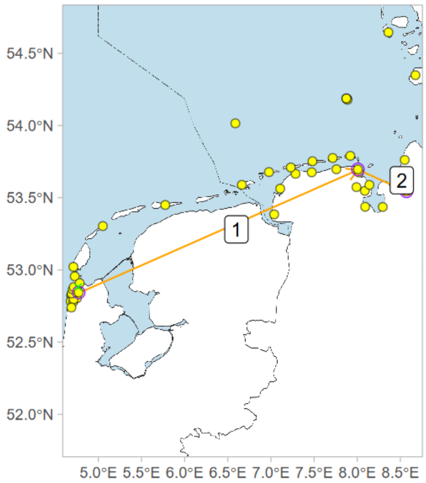


release: 2019-09-24 10:00
1) 2019-10-08 17:38 - 2019-10-10 19:42
2) 2019-10-10 19:42 - 2019-10-13 22:59
final detection: 2019-10-13 22:59

Deployment ID 26143 (adult F)

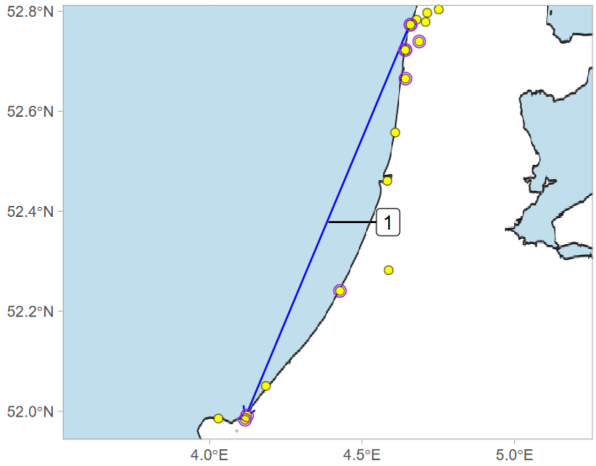


release: 2019-09-17 12:00
1) 2019-10-14 17:44 - 2019-10-14 22:41
final detection: 2019-10-17 19:45

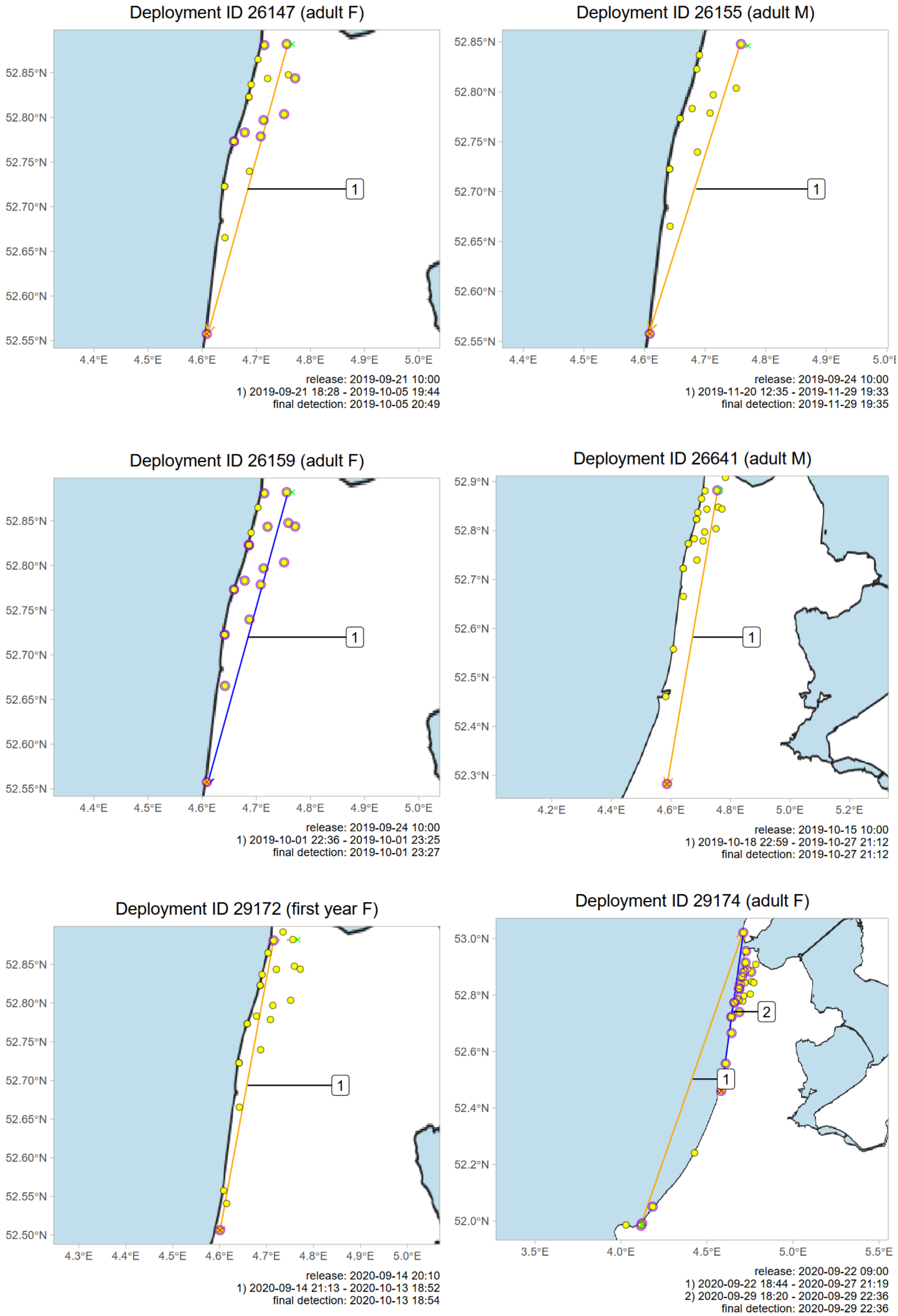

Deployment ID 26147 (adult F)
52.85°N
52.80°N
52.75°N
52.70°N
52.65°N
52.60°N
52.55°N
4.4°E
4.5°E
4.6°E
4.7°E
4.8°E
4.9°E
5.0°E
1
release: 2019-09-21 10:00
1) 2019-09-21 18:28 - 2019-10-05 19:44
final detection: 2019-10-05 20:49
Deployment ID 26155 (adult M)
52.85°N
52.80°N
52.75°N
52.70°N
52.65°N
52.60°N
52.55°N
4.4°E
4.5°E
4.6°E
4.7°E
4.8°E
4.9°E
1
release: 2019-09-24 10:00
1) 2019-11-20 12:35 - 2019-11-29 19:33
final detection: 2019-11-29 19:35
Deployment ID 26159 (adult F)
52.85°N
52.80°N
52.75°N
52.70°N
52.65°N
52.60°N
52.55°N
4.4°E
4.5°E
4.6°E
4.7°E
4.8°E
4.9°E
5.0°E
1
release: 2019-09-24 10:00
1) 2019-10-01 22:36 - 2019-10-01 23:25
final detection: 2019-10-01 23:27
Deployment ID 26641 (adult M)
52.9°N
52.8°N
52.7°N
52.6°N
52.5°N
52.4°N
52.3°N
4.2°E
4.4°E
4.6°E
4.8°E
5.0°E
5.2°E
1
release: 2019-10-15 10:00
1) 2019-10-18 22:59 - 2019-10-27 21:12
final detection: 2019-10-27 21:12
Deployment ID 29172 (first year F)
52.85°N
52.80°N
52.75°N
52.70°N
52.65°N
52.60°N
52.55°N
52.50°N
4.3°E
4.4°E
4.5°E
4.6°E
4.7°E
4.8°E
4.9°E
5.0°E
1
release: 2020-09-14 20:10
1) 2020-09-14 21:13 - 2020-10-13 18:52
final detection: 2020-10-13 18:54
Deployment ID 29174 (adult F)
53.0°N
52.8°N
52.6°N
52.4°N
52.2°N
52.0°N
3.5°E
4.0°E
4.5°E
5.0°E
5.5°E
2
1
release: 2020-09-22 09:00
1) 2020-09-22 18:44 - 2020-09-27 21:19
2) 2020-09-29 18:20 - 2020-09-29 22:36
final detection: 2020-09-29 22:36

Deployment ID 29175 (first year F)

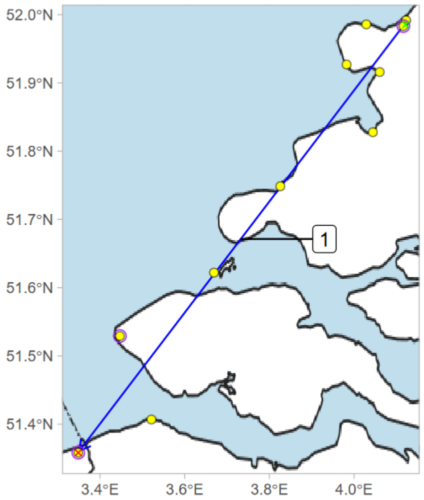


release: 2020-09-13 11:30
1) 2020-09-13 18:55 - 2020-09-14 01:39
final detection: 2020-09-14 01:40

Deployment ID 29176 (first year F)

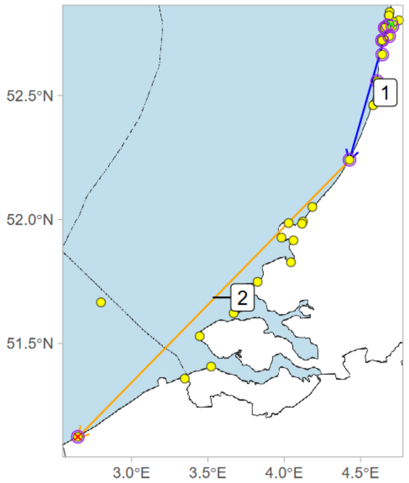


release: 2020-09-02 22:00
1) 2020-09-06 21:06 - 2020-09-07 00:01
2) 2020-09-08 22:04 - 2020-09-10 22:45
final detection: 2020-09-10 22:46

Deployment ID 29177 (adult F)

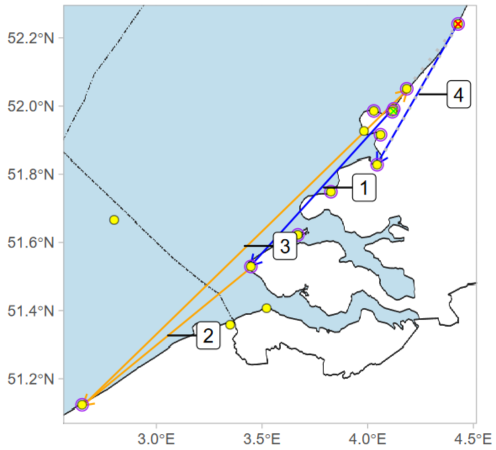


release: 2020-09-01 10:00
1) 2020-09-01 19:15 - 2020-09-01 22:10
2) 2020-09-01 22:10 - 2020-09-02 21:35
3) 2020-09-02 21:35 - 2020-09-07 22:16
4) 2020-09-09 21:12 - 2020-09-10 00:27
final detection: 2020-09-14 20:23

Deployment ID 29179 (adult F)

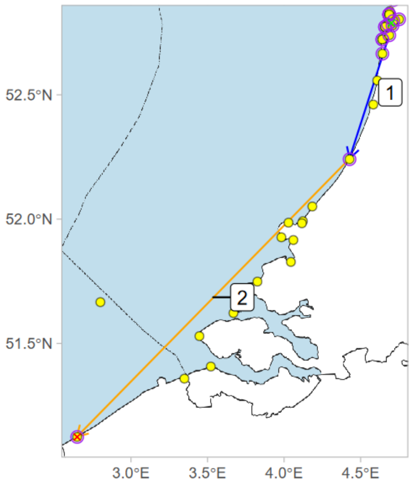


release: 2020-09-13 07:00
1) 2020-09-13 18:58 - 2020-09-13 23:20
2) 2020-09-13 23:20 - 2020-09-14 22:26
final detection: 2020-09-14 22:30

Deployment ID 29181 (first year F)

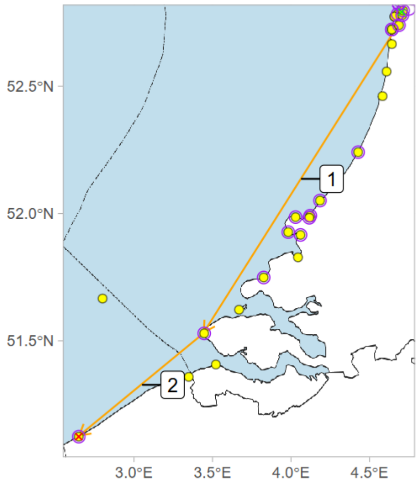


release: 2020-09-04 10:00
1) 2020-09-09 00:40 - 2020-09-10 01:57
2) 2020-09-10 01:57 - 2020-09-10 21:56
final detection: 2020-09-10 21:57

Deployment ID 29188 (first year F)

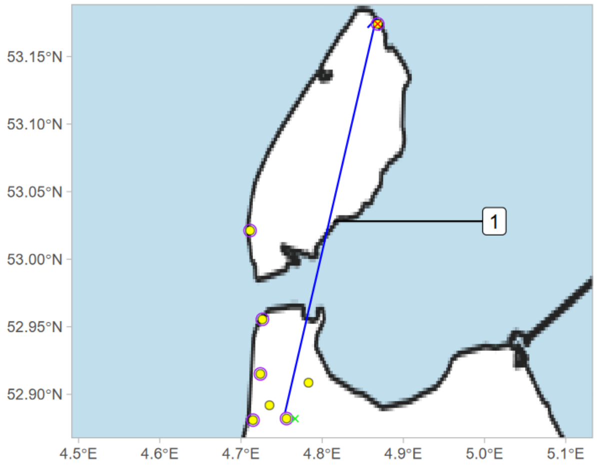


release: 2020-09-22 20:00
1) 2020-09-22 22:43 - 2020-09-22 23:39
final detection: 2020-09-22 23:40

Deployment ID 29191 (first year M)

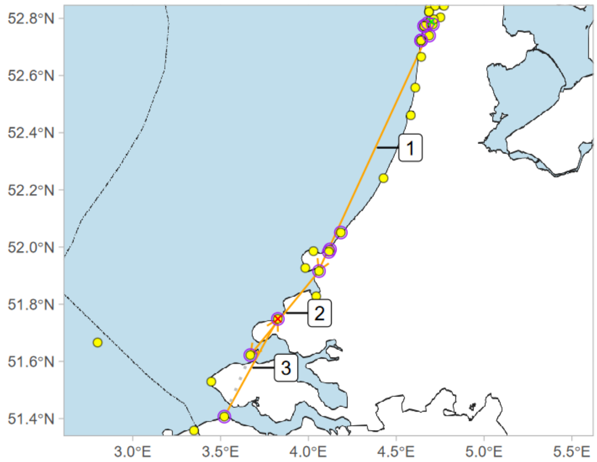


release: 2020-09-02 23:00
1) 2020-09-07 23:22 - 2020-09-10 02:10
2) 2020-09-10 02:10 - 2020-09-18 00:55
3) 2020-09-21 02:41 - 2020-09-21 20:38
final detection: 2020-09-21 20:40

Deployment ID 29194 (adult F)

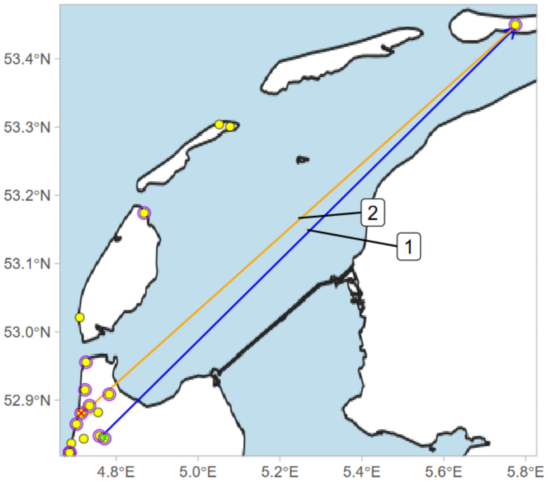


release: 2020-09-13 12:00
1) 2020-09-13 18:41 - 2020-09-13 23:07
2) 2020-09-13 23:07 - 2020-09-14 20:02
final detection: 2020-09-14 20:16

Deployment ID 29195 (first year F)

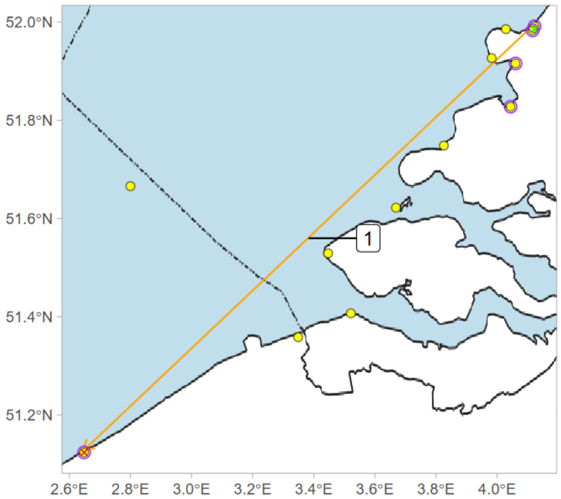


release: 2020-09-01 10:00
1) 2020-09-01 19:23 - 2020-09-02 21:57
final detection: 2020-09-02 21:58

Deployment ID 29200 (first year F)

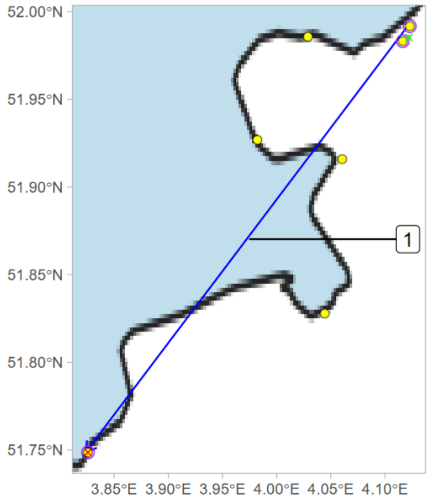


release: 2020-09-15 17:30
1) 2020-09-15 18:56 - 2020-09-15 21:23
final detection: 2020-09-15 21:24

Deployment ID 29201 (adult F)

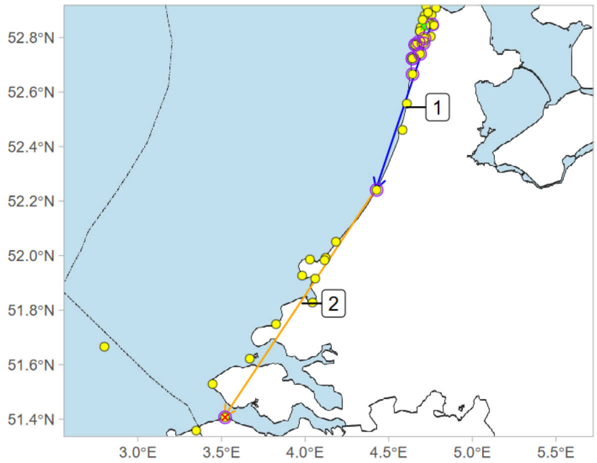


release: 2020-09-13 10:00
1) 2020-09-14 20:10 - 2020-09-14 23:36
2) 2020-09-14 23:36 - 2020-09-16 00:35
final detection: 2020-09-16 01:01

Deployment ID 29207 (adult M)

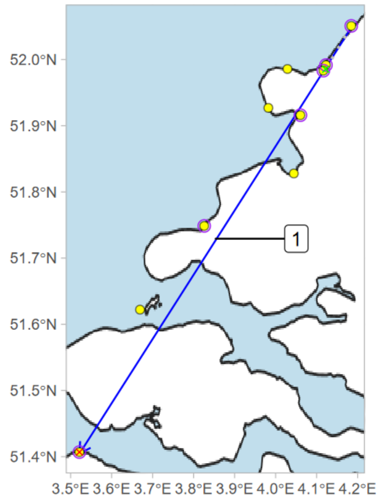


release: 2020-10-13 20:45
1) 2020-10-14 17:31 - 2020-10-14 19:15
final detection: 2020-10-14 19:17

Deployment ID 29210 (adult F)

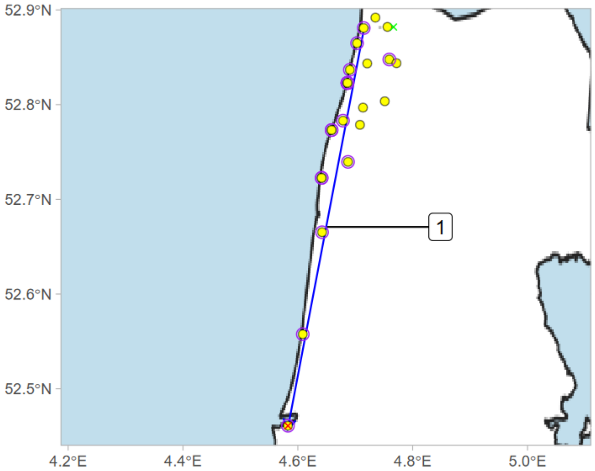


release: 2020-09-14 22:15
1) 2020-09-14 23:23 - 2020-09-15 01:36
final detection: 2020-09-15 01:37

Deployment ID 29212 (first year F)

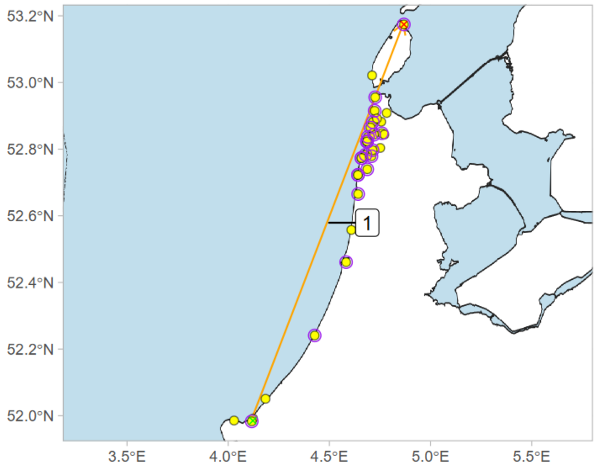


release: 2020-09-01 10:00
1) 2020-09-01 19:37 - 2020-09-14 00:21
final detection: 2020-09-14 00:30

Deployment ID 29213 (first year F)

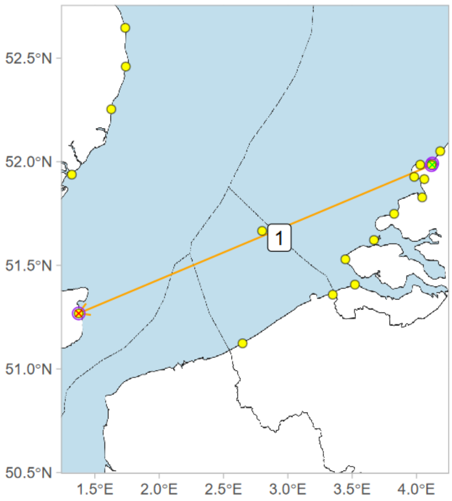


release: 2020-09-15 17:00
1) 2020-09-15 19:39 - 2020-09-22 04:45
final detection: 2020-09-22 04:45

Deployment ID 29219 (first year F)

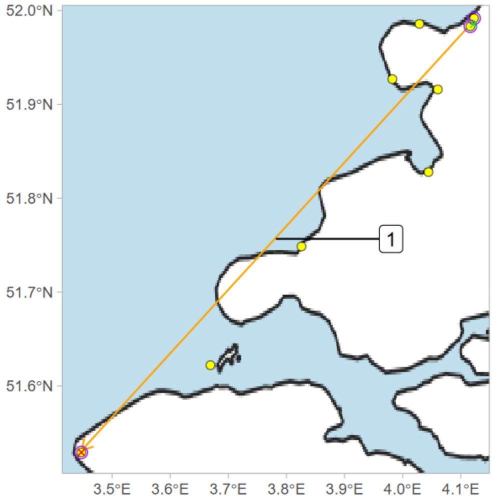


release: 2020-09-22 05:40
1) 2020-09-22 18:25 - 2020-09-29 23:55
final detection: 2020-09-29 23:56

Deployment ID 29233 (first year F)

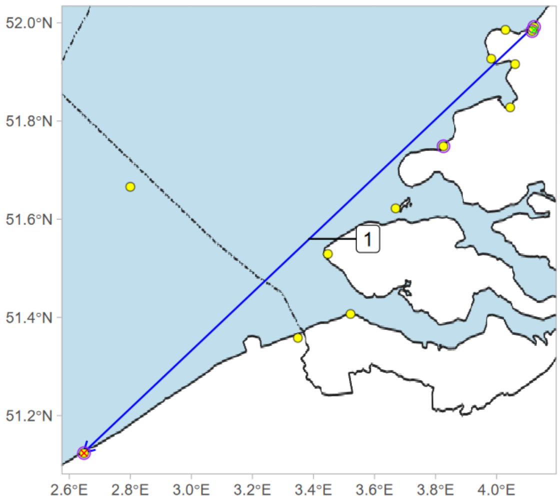


release: 2020-09-01 10:00
1) 2020-09-01 19:02 - 2020-09-02 02:16
final detection: 2020-09-02 21:50

Deployment ID 29234 (first year F)

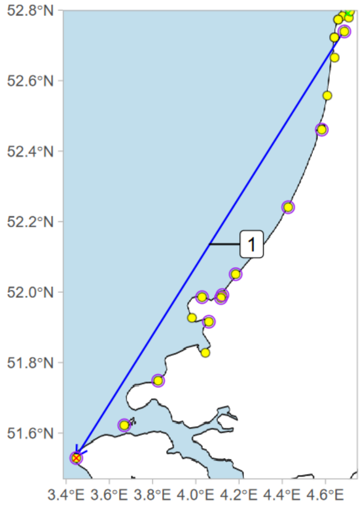


release: 2020-09-08 11:30
1) 2020-09-09 20:05 - 2020-09-10 01:31
final detection: 2020-09-10 01:36

Deployment ID 29259 (adult M)

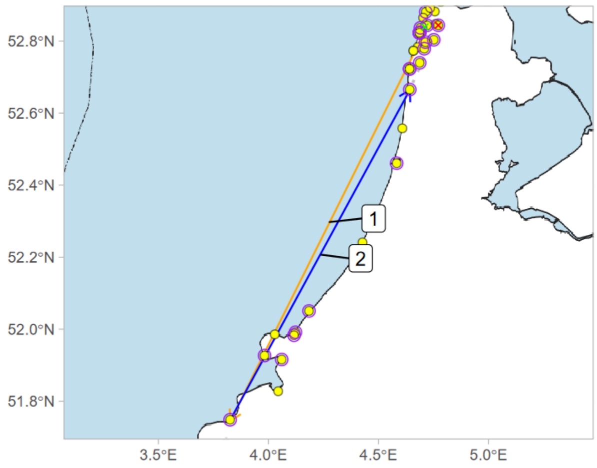


release: 2020-09-13 10:00
1) 2020-10-02 19:19 - 2020-10-03 21:03
2) 2020-10-05 17:43 - 2020-10-05 19:48
final detection: 2020-10-07 17:47

Deployment ID 29262 (first year F)

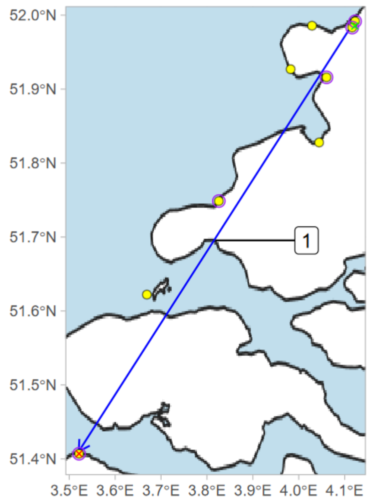


release: 2020-09-15 17:30
1) 2020-09-15 19:53 - 2020-09-16 02:14
final detection: 2020-09-16 02:18

Deployment ID 29263 (first year M)

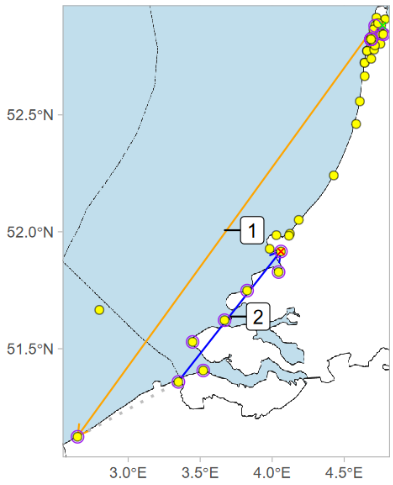


release: 2020-09-08 07:00
1) 2020-09-09 00:17 - 2020-09-11 00:25
2) 2020-09-23 23:00 - 2020-09-24 00:10
final detection: 2020-09-24 00:11

Deployment ID 29277 (first year F)

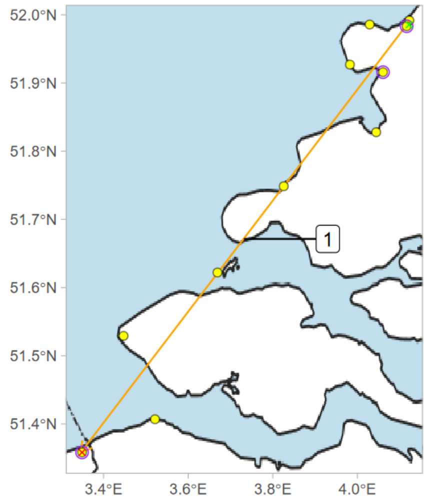


release: 2020-09-04 05:30
1) 2020-09-05 22:12 - 2020-09-09 23:20
final detection: 2020-09-09 23:23

Deployment ID 29280 (adult F)

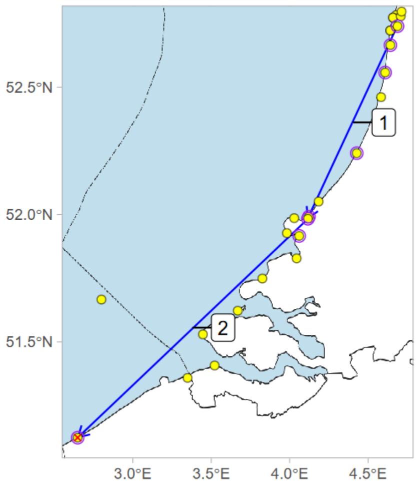


release: 2020-09-13 10:00
1) 2020-09-13 20:44 - 2020-09-14 02:13
2) 2020-09-14 19:14 - 2020-09-14 23:27
final detection: 2020-09-14 23:28

Deployment ID 29294 (first year F)

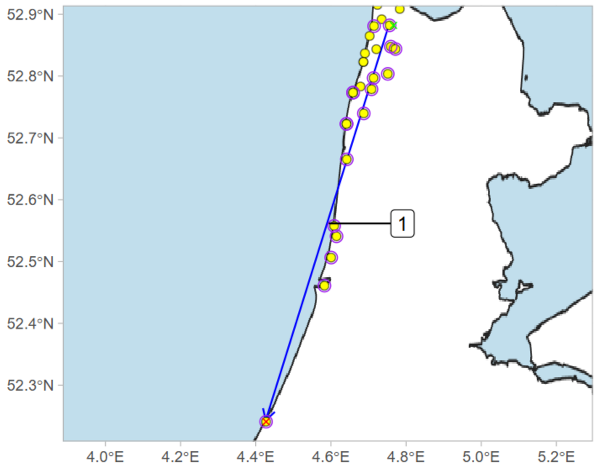


release: 2020-09-11 20:55
1) 2020-10-10 17:44 - 2020-10-10 19:58
final detection: 2020-10-10 20:01

Deployment ID 29874 (adult F)

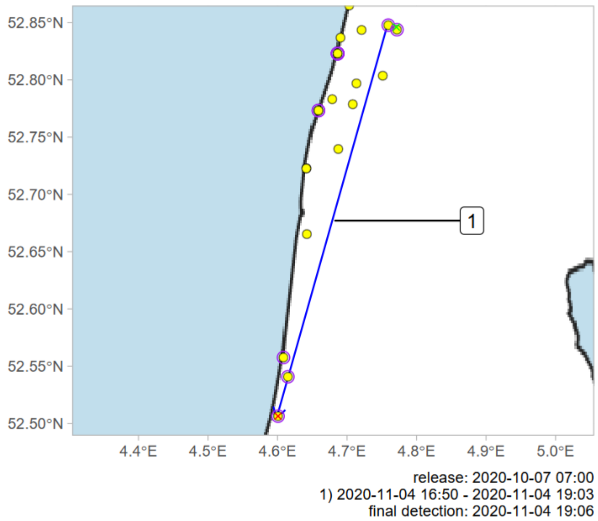


Deployment ID 30088 (adult F)

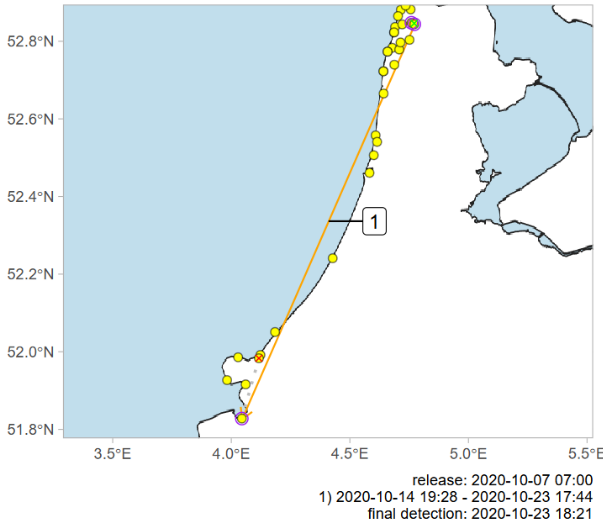


Deployment ID 35227 (adult F)

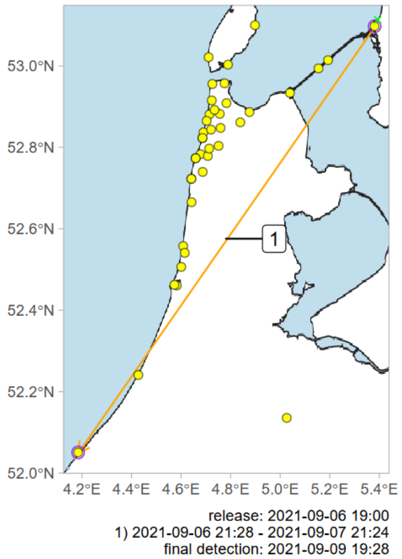


Deployment ID 35230 (adult F)

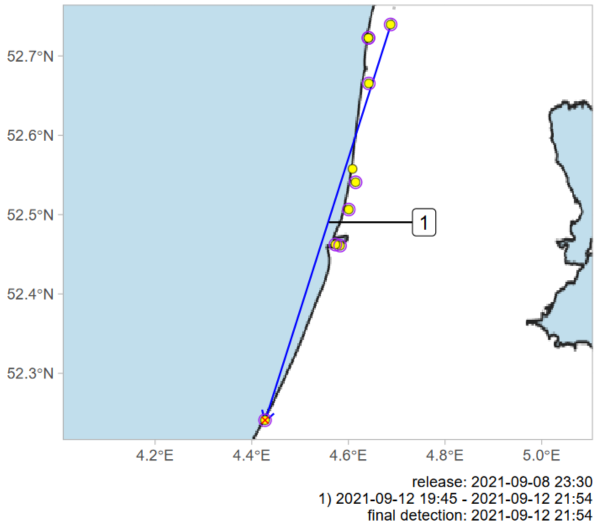


Deployment ID 35235 (adult F)

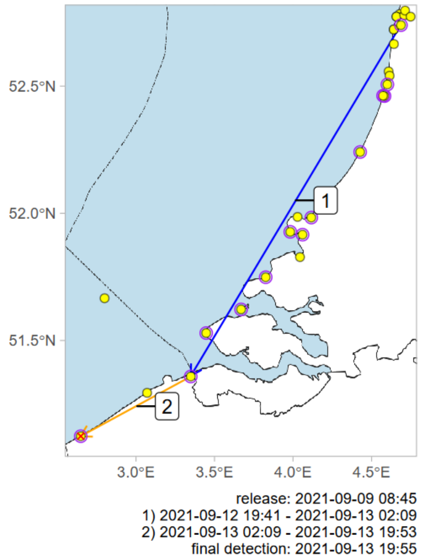


Deployment ID 35237 (adult F)

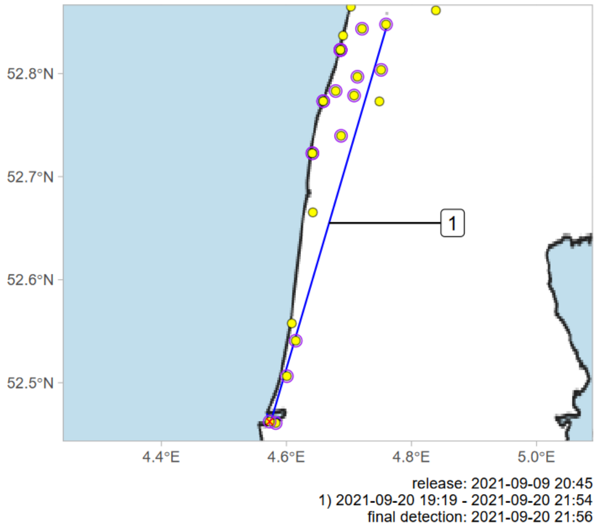

Deployment ID 35239 (adult F)

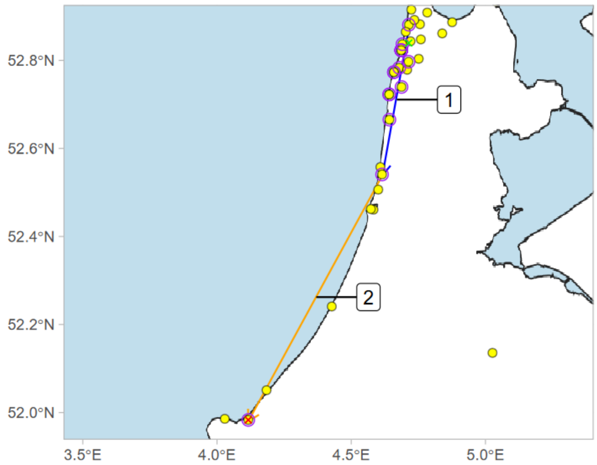


release: 2021-08-21 12:00
1) 2021-08-21 19:38 - 2021-08-21 21:13
2) 2021-08-21 21:13 - 2021-08-23 19:57
final detection: 2021-08-23 19:57

Deployment ID 35241 (adult F)

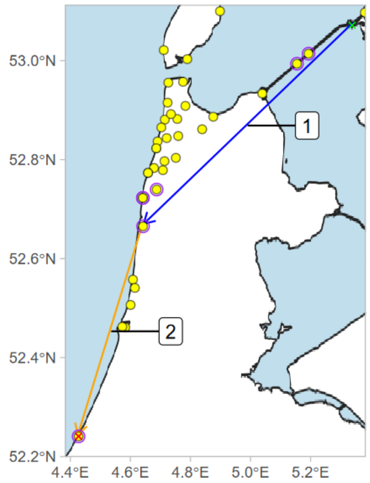


release: 2021-09-07 20:10
1) 2021-09-07 20:10 - 2021-09-08 01:00
2) 2021-09-08 01:00 - 2021-09-08 21:00
final detection: 2021-09-08 21:02

Deployment ID 35245 (adult F)

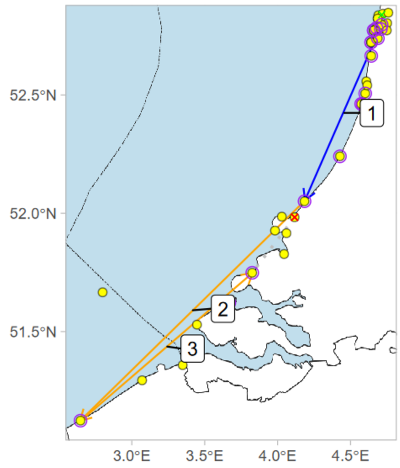


release: 2021-09-09 12:00
1) 2021-09-09 19:45 - 2021-09-10 00:36
2) 2021-09-10 00:36 - 2021-09-12 23:31
3) 2021-09-12 23:31 - 2021-09-18 21:57
final detection: 2021-09-21 19:44

Deployment ID 35247 (first year F)

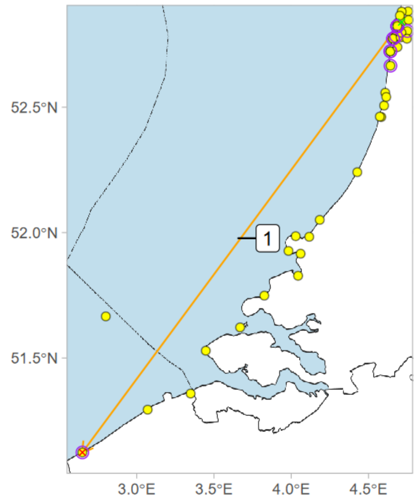


release: 2021-09-09 12:00
1) 2021-09-09 19:04 - 2021-09-14 00:11
final detection: 2021-09-14 00:12

Deployment ID 35251 (first year F)

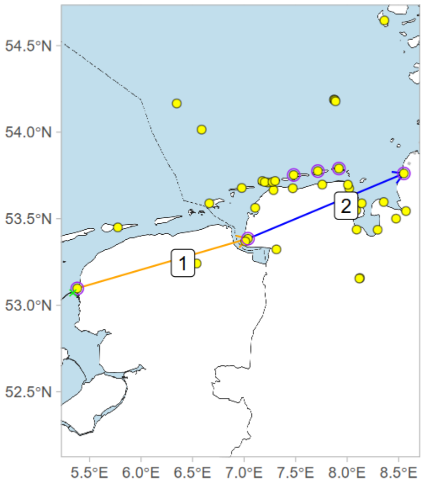


release: 2021-09-21 23:30
1) 2021-09-21 23:56 - 2021-09-22 19:19
2) 2021-09-22 19:19 - 2021-09-22 23:39
final detection: 2021-09-27 23:42

Deployment ID 35253 (adult F)

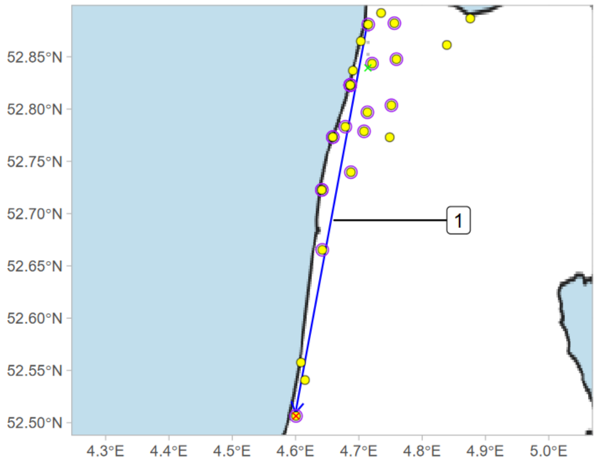


release: 2021-09-24 09:00
1) 2021-10-06 19:59 - 2021-10-06 20:55
final detection: 2021-10-06 21:02

Deployment ID 35254 (adult F)

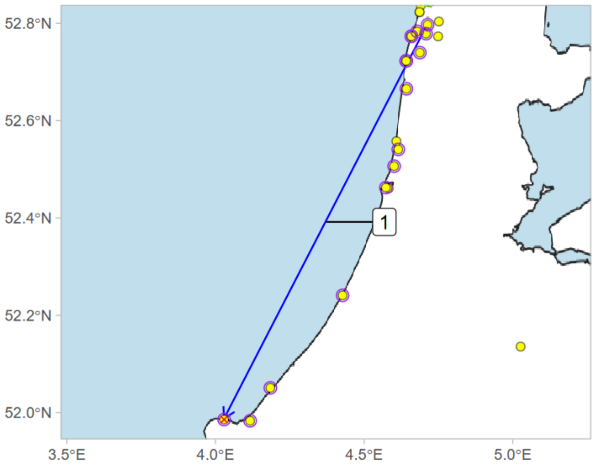


release: 2021-09-09 22:35
1) 2021-09-12 19:47 - 2021-09-13 00:18
final detection: 2021-09-13 00:18

Deployment ID 35255 (adult M)

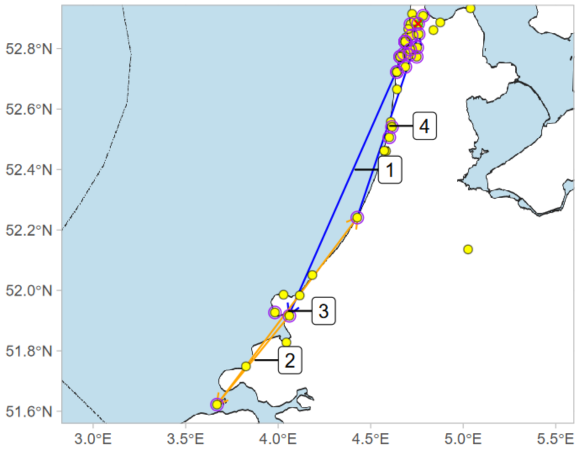


release: 2021-09-08 22:00
1) 2021-09-13 18:50 - 2021-09-14 02:41
2) 2021-09-14 18:42 - 2021-09-17 00:14
3) 2021-09-17 00:14 - 2021-09-24 02:22
4) 2021-09-24 02:22 - 2021-09-24 04:59
final detection: 2021-09-26 03:18

Deployment ID 35257 (adult M)

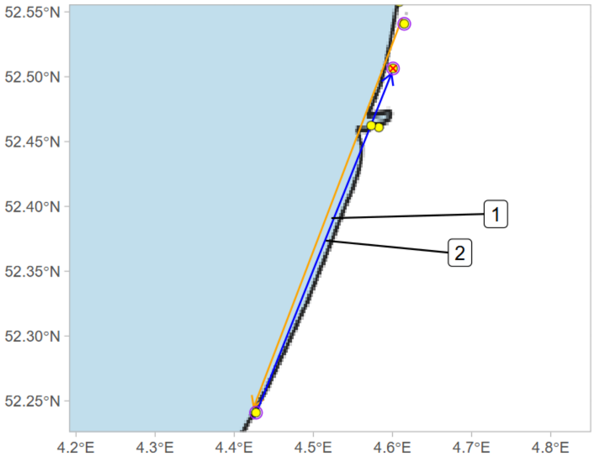


release: 2021-09-10 01:10
1) 2021-09-16 21:09 - 2021-09-17 21:24
2) 2021-09-17 21:24 - 2021-09-18 00:22
final detection: 2021-09-18 19:39

Deployment ID 35267 (adult M)

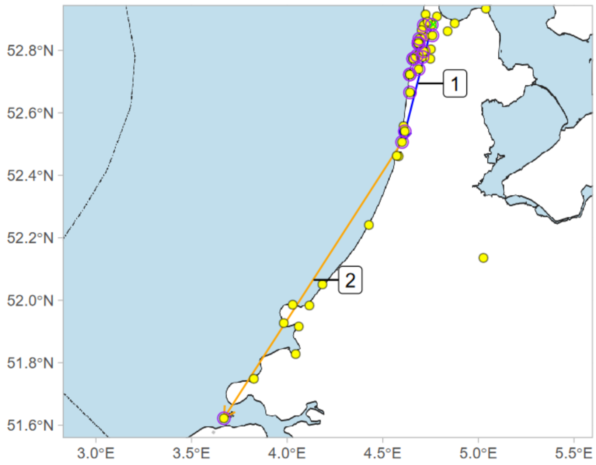


release: 2021-09-18 21:10
1) 2021-09-18 21:34 - 2021-09-19 01:31
2) 2021-09-19 19:24 - 2021-09-25 20:02
final detection: 2021-09-27 22:44

Deployment ID 35271 (adult F)

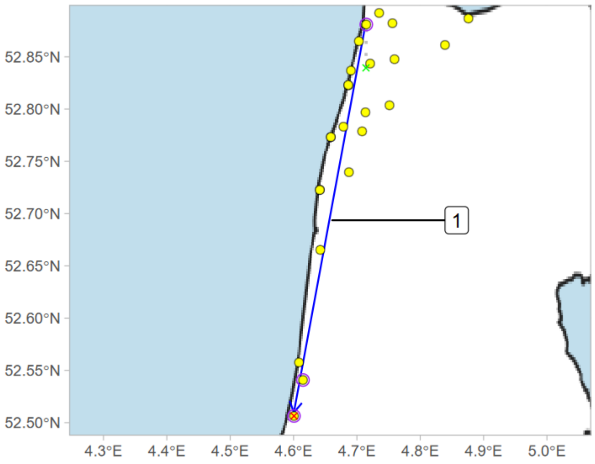


release: 2021-09-09 12:00
1) 2021-09-09 20:18 - 2021-09-09 23:26
final detection: 2021-09-09 23:41

Deployment ID 35272 (adult F)

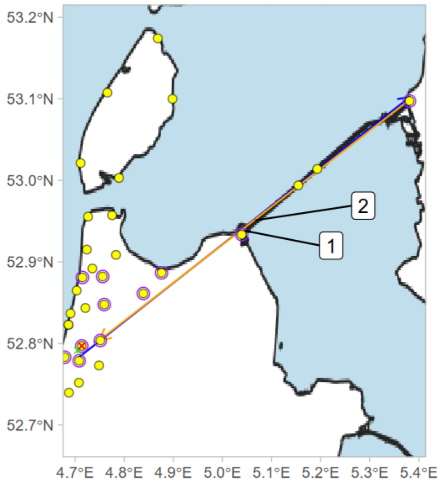


release: 2021-09-18 12:00
1) 2021-09-21 18:32 - 2021-09-21 21:11
2) 2021-09-21 23:46 - 2021-09-26 00:34
final detection: 2021-11-16 11:59

### Deployment ID 35275 (adult F)

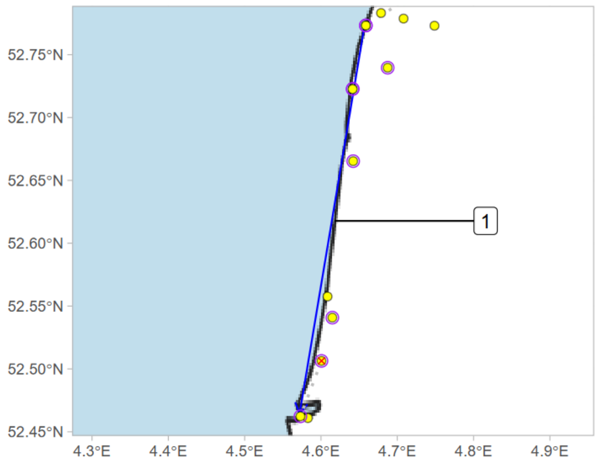


release: 2021-09-08 21:25
1) 2021-09-13 20:00 - 2021-09-13 20:55
final detection: 2021-09-13 21:02

### Deployment ID 35276 (first year M)

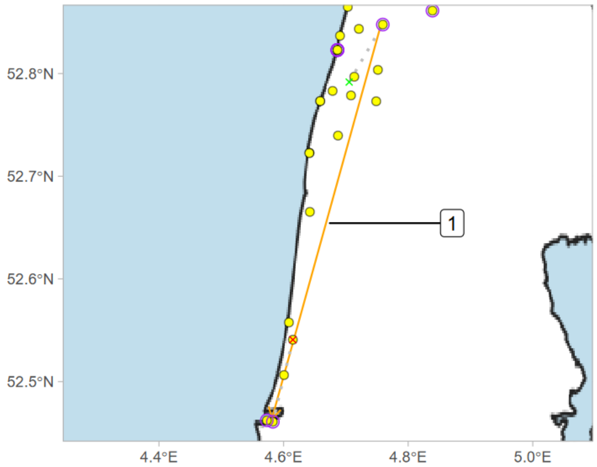


release: 2021-09-08 20:55
1) 2021-09-12 19:30 - 2021-09-17 20:31
final detection: 2021-09-17 20:58

### Deployment ID 35277 (first year M)

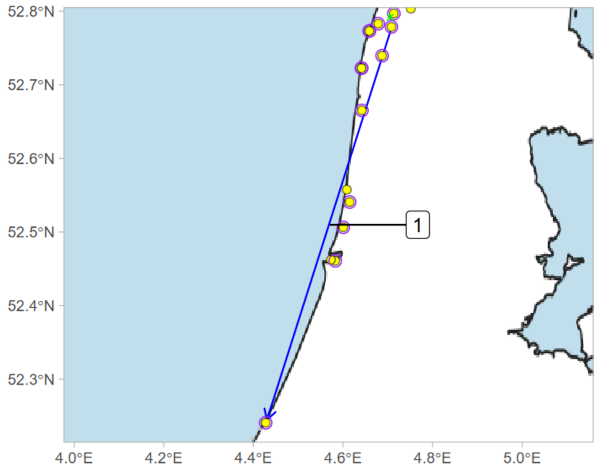


release: 2021-09-08 20:50
1) 2021-09-08 21:06 - 2021-09-09 00:28
final detection: 2021-09-09 22:03

### Deployment ID 35280 (first year M)

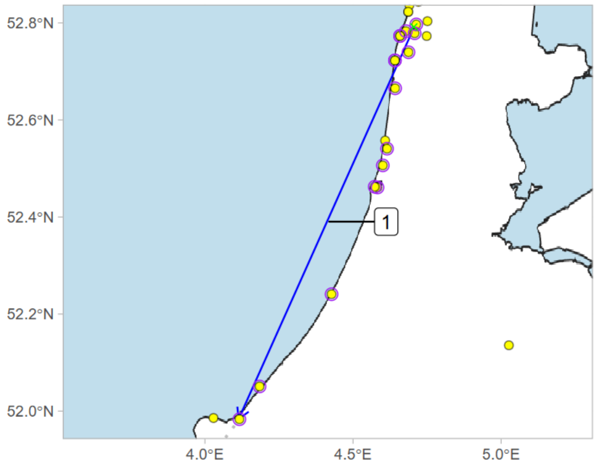


release: 2021-09-08 20:10
1) 2021-09-12 18:52 - 2021-09-12 22:57
final detection: 2021-09-13 21:24

### Deployment ID 35349 (adult F)

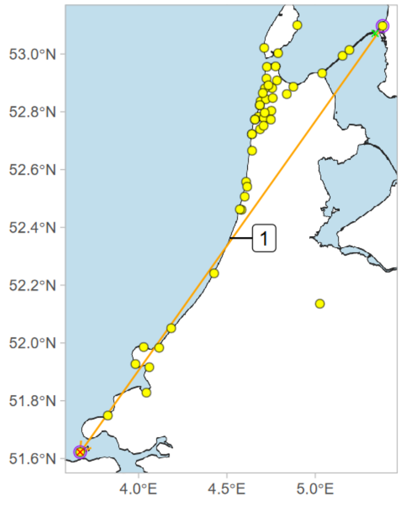


release: 2021-09-07 19:50
1) 2021-09-07 21:55 - 2021-10-08 00:20
final detection: 2021-10-08 00:21

### Deployment ID 35362 (first year F)

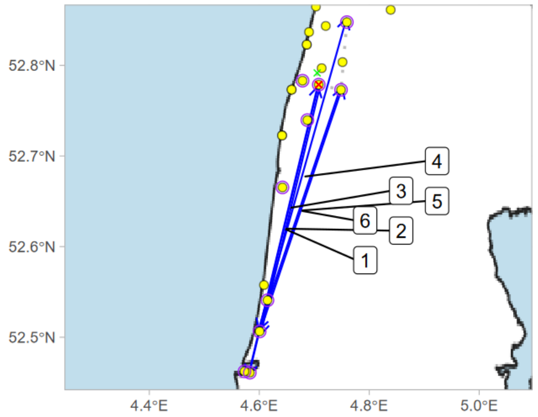


release: 2021-09-08 23:50
1) 2021-09-09 19:30 - 2021-09-09 21:16
2) 2021-09-09 21:16 - 2021-09-10 02:42
3) 2021-09-12 18:57 - 2021-09-12 20:24
4) 2021-09-12 20:24 - 2021-09-13 03:03
5) 2021-09-13 18:57 - 2021-09-13 20:53
6) 2021-09-13 21:57 - 2021-09-14 00:24
final detection: 2021-09-28 18:21

### Deployment ID 35366 (first year F)

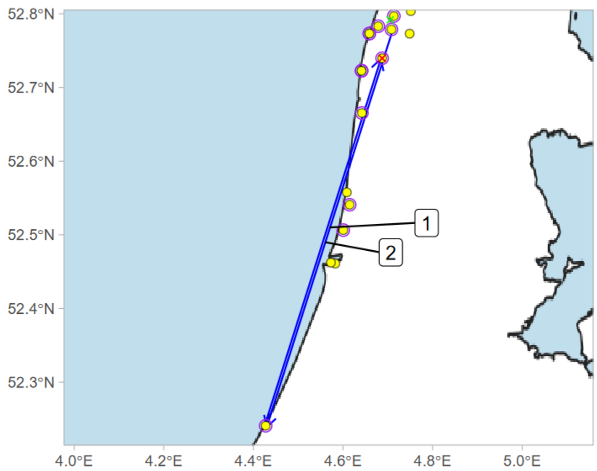


release: 2021-09-09 12:30
1) 2021-09-09 19:01 - 2021-09-09 22:53
2) 2021-09-10 00:21 - 2021-09-10 03:08
final detection: 2021-09-12 19:01

### Deployment ID 35372 (first year M)

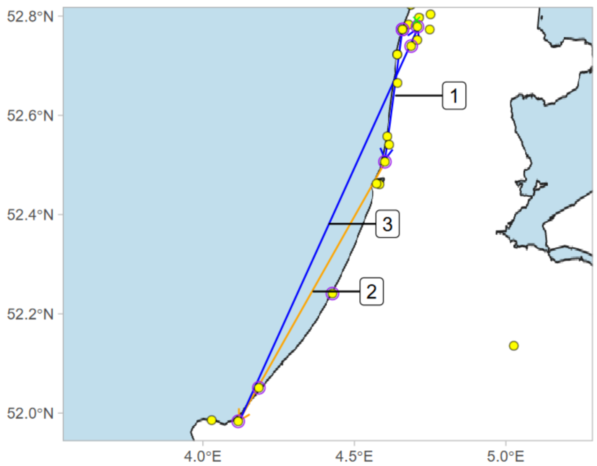


release: 2021-09-08 23:10
1) 2021-09-13 19:29 - 2021-09-13 20:46
2) 2021-09-13 20:46 - 2021-09-19 22:15
3) 2021-09-19 22:15 - 2021-09-20 02:53
final detection: 2021-10-09 21:29

### Deployment ID 35561 (first year M)

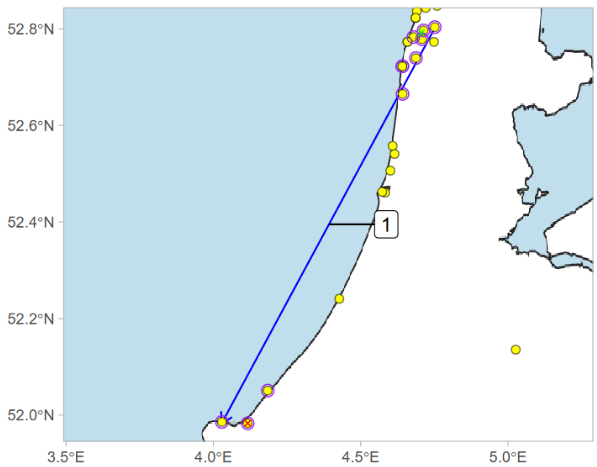


release: 2021-09-17 16:00
1) 2021-09-17 18:52 - 2021-09-18 00:02
final detection: 2021-09-18 00:21

### Deployment ID 35562 (adult F)

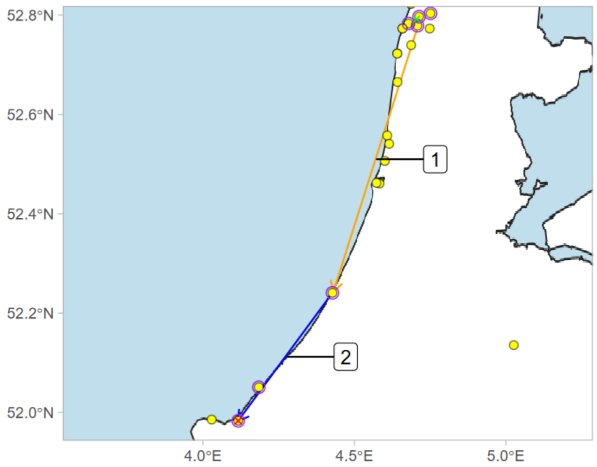


release: 2021-09-09 19:25
1) 2021-09-09 19:43 - 2021-09-17 22:46
2) 2021-09-17 22:46 - 2021-09-18 00:31
final detection: 2021-09-18 00:31

### Deployment ID 35570 (adult F)

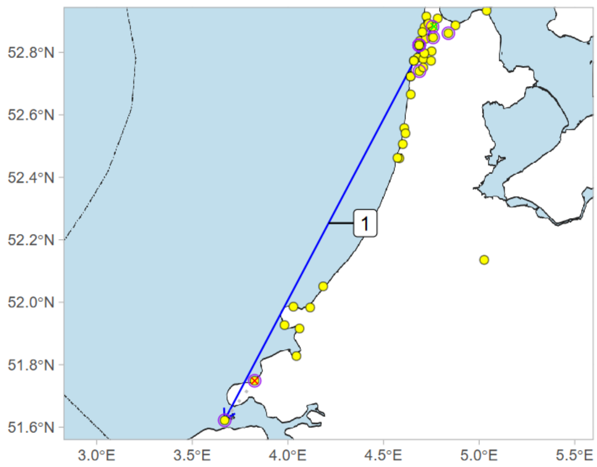


release: 2021-10-12 08:30
1) 2021-10-12 17:30 - 2021-10-13 01:54
final detection: 2021-10-13 17:58

### Deployment ID 35571 (first year M)

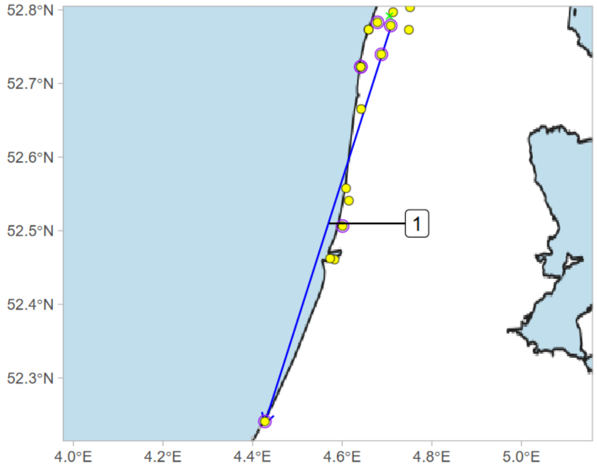


release: 2021-09-09 22:20
1) 2021-09-12 18:50 - 2021-09-12 22:02
final detection: 2021-09-16 20:17

Deployment ID 35572 (first year M)

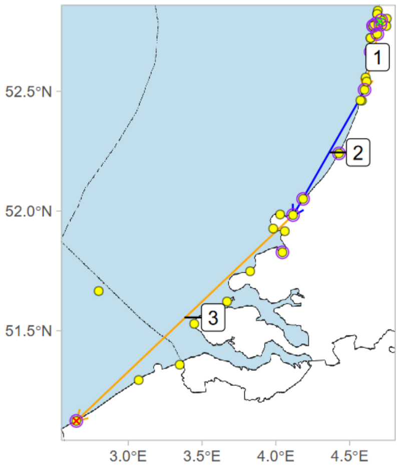


release: 2021-09-09 22:40
1) 2021-09-09 23:18 - 2021-09-12 19:39
2) 2021-09-12 19:39 - 2021-09-12 23:14
3) 2021-09-12 23:14 - 2021-09-20 00:11
final detection: 2021-09-20 00:12

Deployment ID 35578 (adult F)

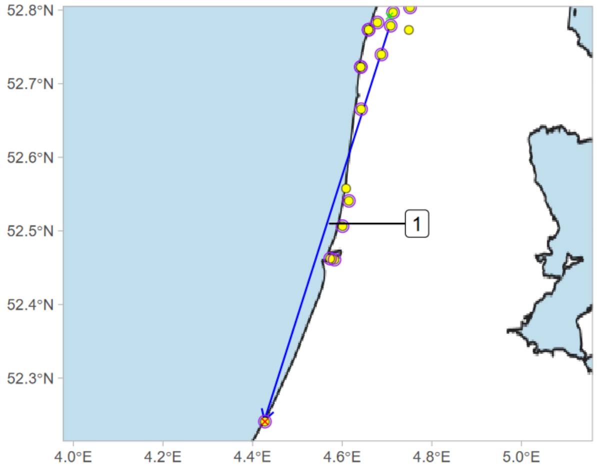


release: 2021-09-09 12:30
1) 2021-09-20 19:08 - 2021-09-20 22:36
final detection: 2021-09-20 22:37

Deployment ID 35580 (adult M)

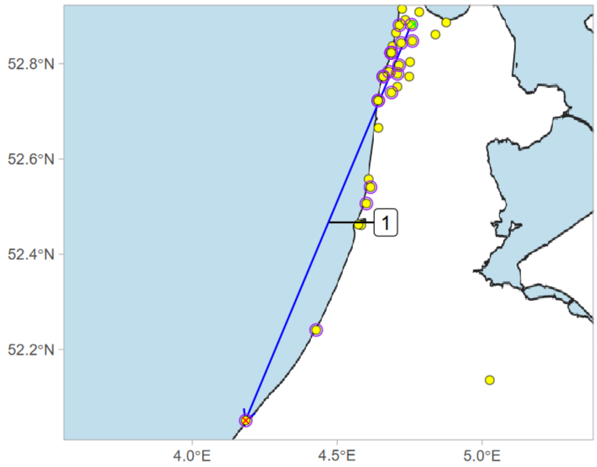


release: 2021-10-12 07:35
1) 2021-10-12 18:10 - 2021-10-12 20:29
final detection: 2021-10-12 20:30

Deployment ID 35585 (first year F)

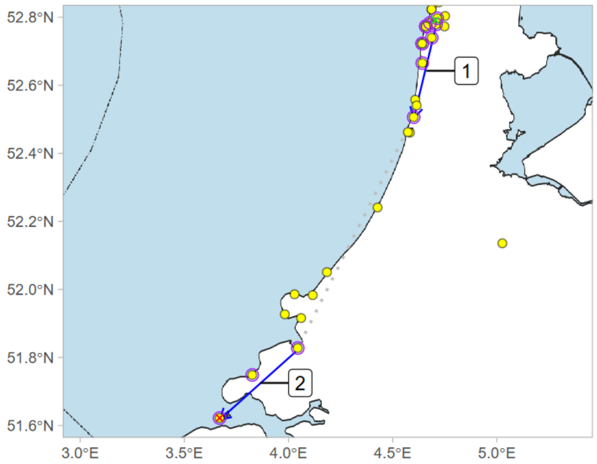


release: 2021-09-18 12:52
1) 2021-09-18 18:41 - 2021-09-18 20:26
2) 2021-09-19 19:29 - 2021-09-19 20:49
final detection: 2021-09-19 23:38

Deployment ID 35586 (first year M)

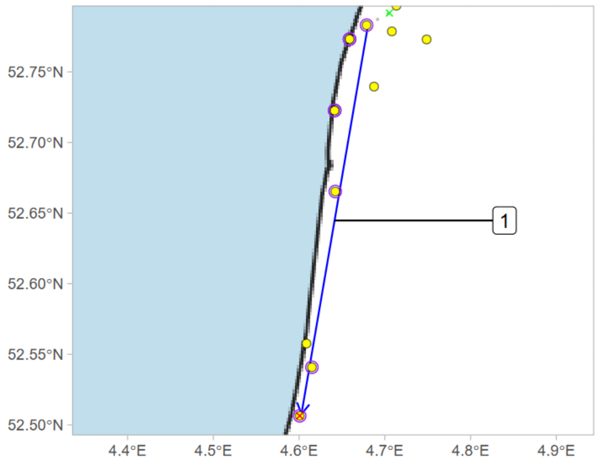


release: 2021-09-18 21:00
1) 2021-09-18 21:24 - 2021-09-18 23:12
final detection: 2021-09-18 23:14

Deployment ID 35587 (first year F)

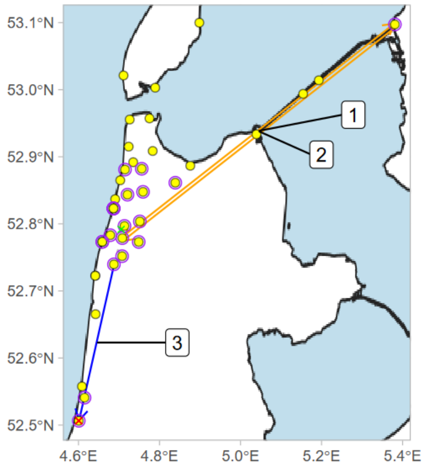


release: 2021-09-18 13:10
1) 2021-09-22 18:34 - 2021-09-25 21:20
2) 2021-09-25 21:20 - 2021-09-26 22:07
3) 2021-10-10 18:17 - 2021-10-10 19:18
final detection: 2021-10-10 19:21

Deployment ID 35588 (first year F)

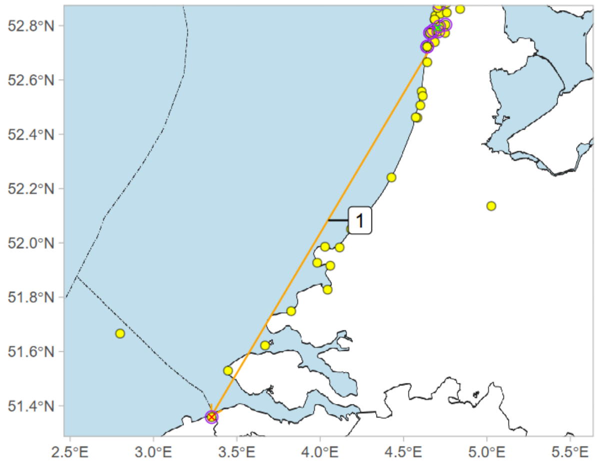


release: 2021-09-09 12:30
1) 2021-09-09 19:02 - 2021-09-18 21:17
final detection: 2021-09-18 21:17

Deployment ID 36029 (first year M)

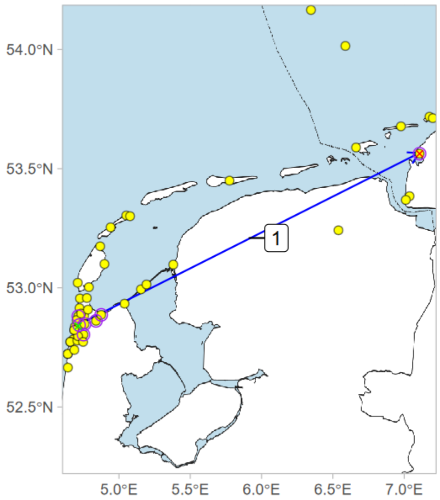


release: 2021-09-18 21:20
1) 2021-09-21 18:47 - 2021-09-22 00:22
final detection: 2021-09-22 00:34

Deployment ID 36045 (first year M)

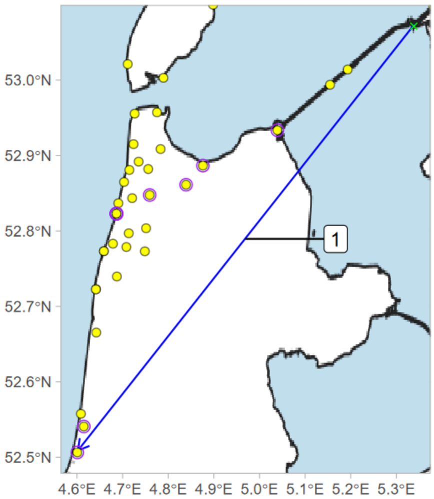


release: 2021-09-26 19:00
1) 2021-09-26 19:00 - 2021-09-27 02:58
final detection: 2021-09-27 23:49

Deployment ID 36234 (adult F)

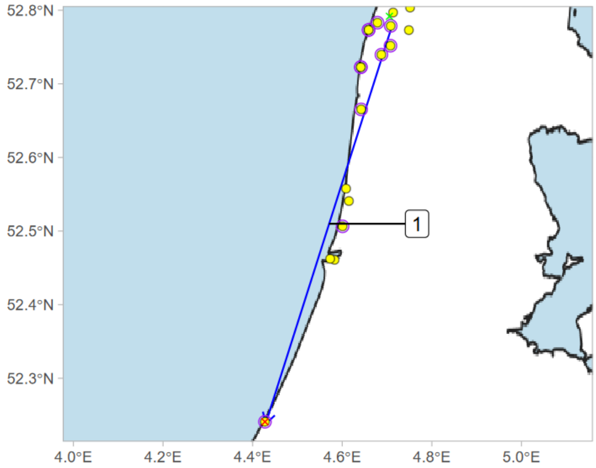


release: 2021-09-24 11:55
1) 2021-10-15 17:38 - 2021-10-15 19:15
final detection: 2021-10-15 19:16

Deployment ID 36236 (first year F)

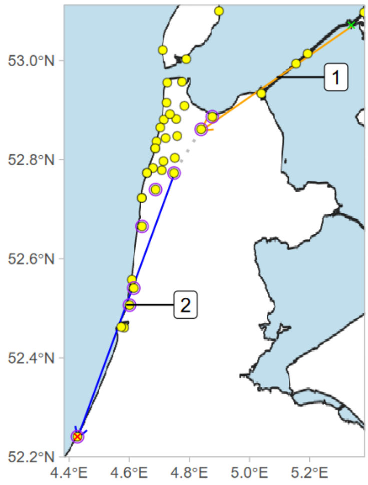


release: 2021-09-21 20:40
1) 2021-09-21 22:40 - 2021-09-26 22:23
2) 2021-09-27 19:47 - 2021-09-28 00:39
final detection: 2021-09-28 00:43

Deployment ID 36237 (first year F)

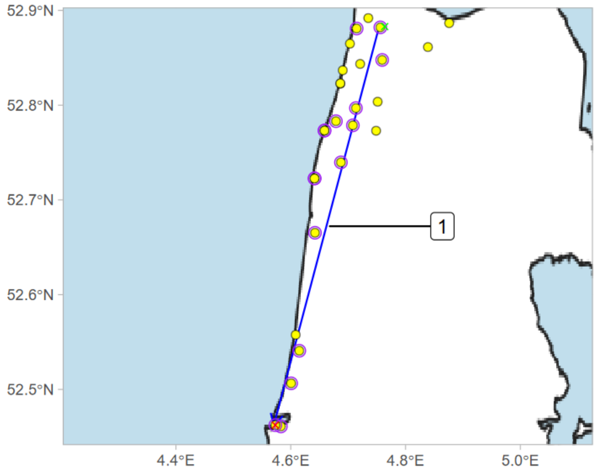


release: 2021-09-23 09:00
1) 2021-10-06 17:39 - 2021-10-06 19:53
final detection: 2021-10-06 19:57

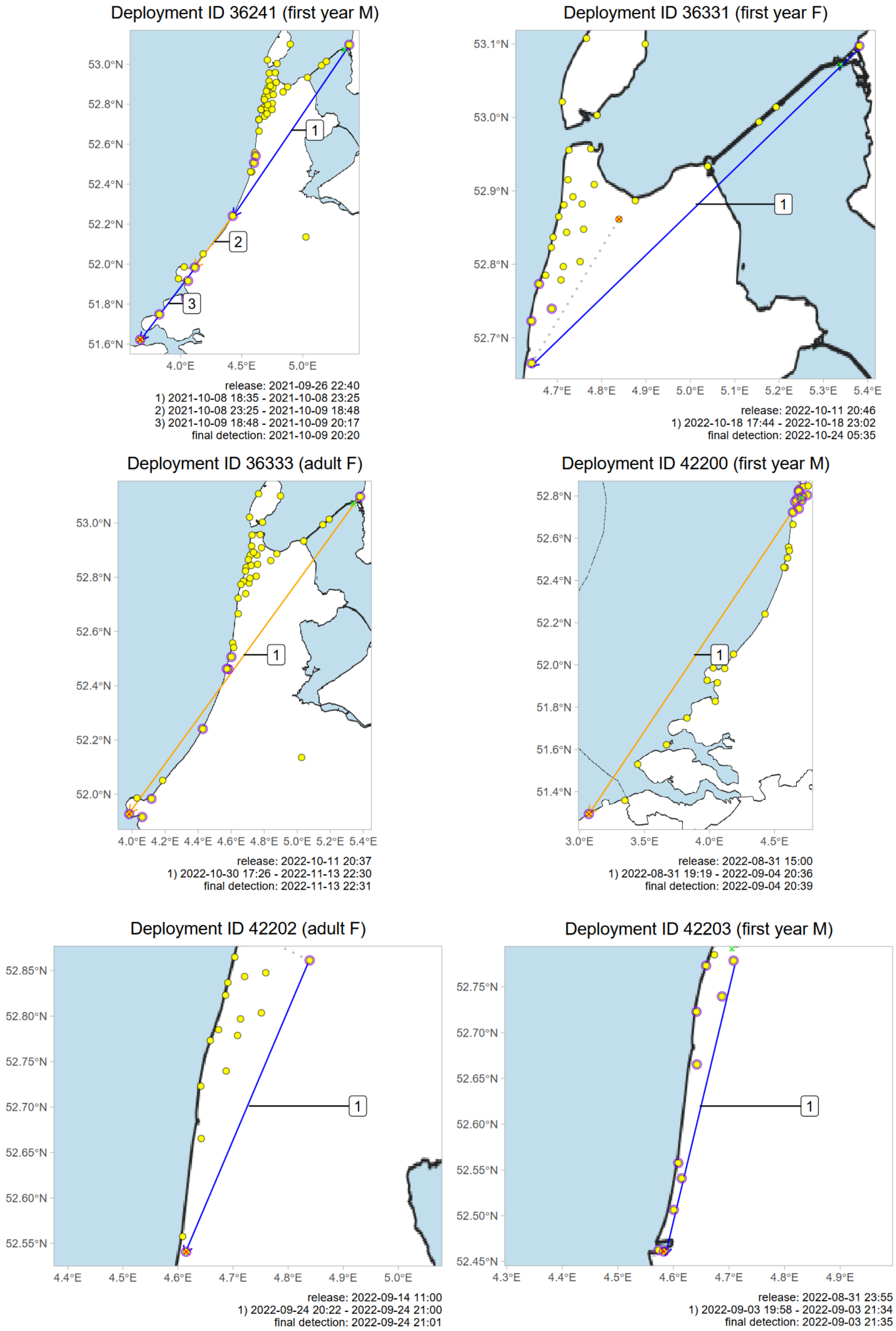

Deployment ID 36241 (first year M)
53.0°N
52.8°N
52.6°N
52.4°N
52.2°N
52.0°N
51.8°N
51.6°N
4.0°E
4.5°E
5.0°E
1
2
3
release: 2021-09-26 22:40
1) 2021-10-08 18:35 - 2021-10-08 23:25
2) 2021-10-08 23:25 - 2021-10-09 18:48
3) 2021-10-09 18:48 - 2021-10-09 20:17
final detection: 2021-10-09 20:20
Deployment ID 36331 (first year F)
53.1°N
53.0°N
52.9°N
52.8°N
52.7°N
4.7°E
4.8°E
4.9°E
5.0°E
5.1°E
5.2°E
5.3°E
5.4°E
1
release: 2022-10-11 20:46
1) 2022-10-18 17:44 - 2022-10-18 23:02
final detection: 2022-10-24 05:35
Deployment ID 36333 (adult F)
53.0°N
52.8°N
52.6°N
52.4°N
52.2°N
52.0°N
4.0°E
4.2°E
4.4°E
4.6°E
4.8°E
5.0°E
5.2°E
5.4°E
1
release: 2022-10-11 20:37
1) 2022-10-30 17:26 - 2022-11-13 22:30
final detection: 2022-11-13 22:31
Deployment ID 42200 (first year M)
52.8°N
52.6°N
52.4°N
52.2°N
52.0°N
51.8°N
51.6°N
51.4°N
3.0°E
3.5°E
4.0°E
4.5°E
1
release: 2022-08-31 15:00
1) 2022-08-31 19:19 - 2022-09-04 20:36
final detection: 2022-09-04 20:39
Deployment ID 42202 (adult F)
52.85°N
52.80°N
52.75°N
52.70°N
52.65°N
52.60°N
52.55°N
4.4°E
4.5°E
4.6°E
4.7°E
4.8°E
4.9°E
5.0°E
1
release: 2022-09-14 11:00
1) 2022-09-24 20:22 - 2022-09-24 21:00
final detection: 2022-09-24 21:01
Deployment ID 42203 (first year M)
52.75°N
52.70°N
52.65°N
52.60°N
52.55°N
52.50°N
52.45°N
4.3°E
4.4°E
4.5°E
4.6°E
4.7°E
4.8°E
4.9°E
1
release: 2022-08-31 23:55
1) 2022-09-03 19:58 - 2022-09-03 21:34
final detection: 2022-09-03 21:35

Deployment ID 42204 (first year M)

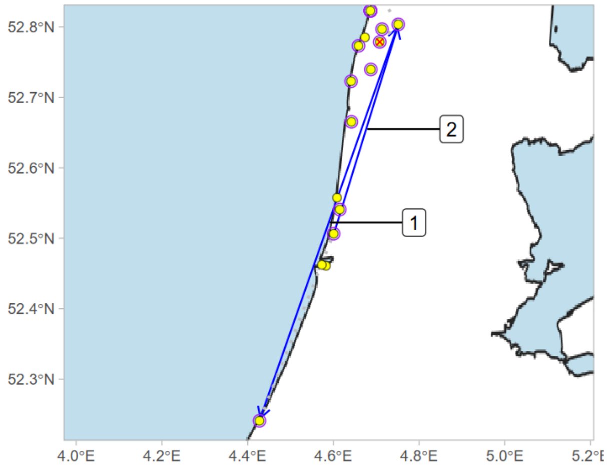


release: 2022-09-12 15:00
1) 2022-09-24 18:55 - 2022-09-24 20:12
2) 2022-09-25 20:38 - 2022-09-25 22:44
final detection: 2022-10-31 17:19

Deployment ID 42206 (first year M)

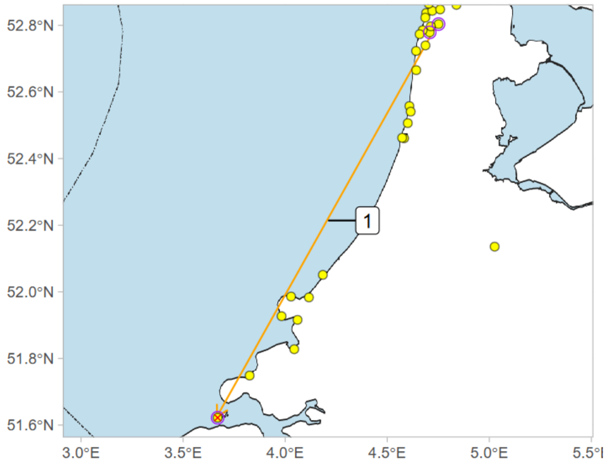


release: 2022-09-06 08:30
1) 2022-09-12 21:30 - 2022-09-14 20:45
final detection: 2022-09-14 20:47

Deployment ID 42209 (adult F)

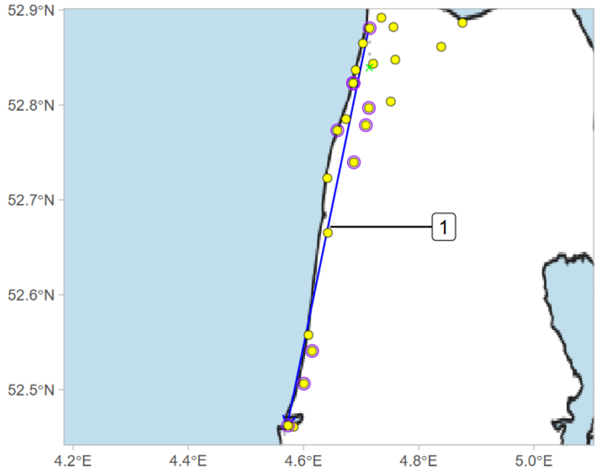


release: 2022-09-06 15:15
1) 2022-09-06 19:05 - 2022-09-06 21:07
final detection: 2022-09-08 20:42

Deployment ID 42214 (first year F)

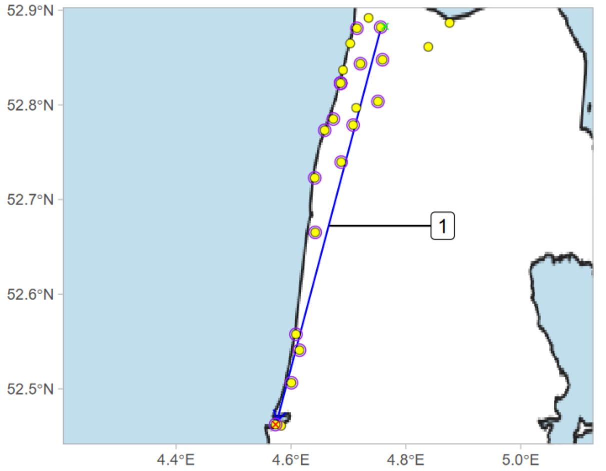


release: 2022-09-01 01:00
1) 2022-09-01 19:25 - 2022-09-01 21:21
final detection: 2022-09-01 21:22

Deployment ID 42215 (adult F)

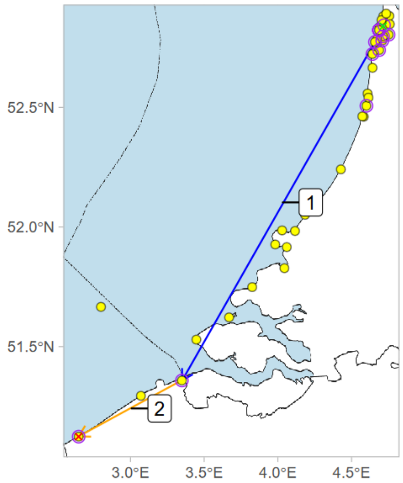


release: 2022-09-06 15:15
1) 2022-09-10 18:54 - 2022-09-11 01:50
2) 2022-09-11 01:50 - 2022-09-11 20:39
final detection: 2022-09-11 20:39

Deployment ID 42216 (first year M)

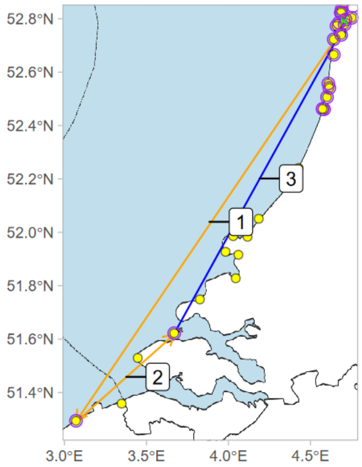


release: 2022-08-31 23:55
1) 2022-09-01 19:26 - 2022-09-02 19:59
2) 2022-09-02 20:09 - 2022-09-03 19:07
3) 2022-09-03 19:07 - 2022-09-04 01:55
final detection: 2022-09-05 20:28

Deployment ID 42220 (first year F)

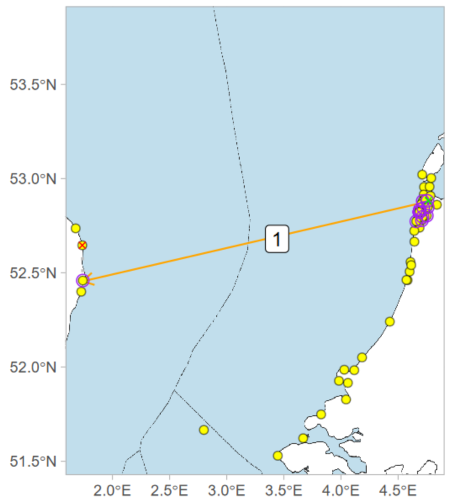


release: 2022-09-01 01:40
1) 2022-09-01 19:21 - 2022-09-09 19:48
final detection: 2022-09-10 22:27

Deployment ID 42221 (first year F)

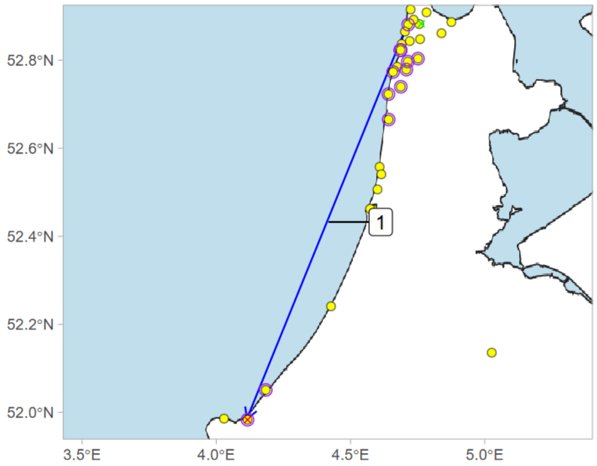


release: 2022-09-06 08:30
1) 2022-09-13 19:47 - 2022-09-13 22:24
final detection: 2022-09-13 22:36

Deployment ID 42222 (first year M)

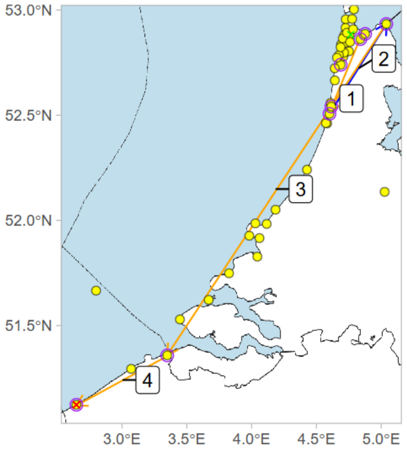


release: 2022-09-01 21:30
1) 2022-09-01 22:38 - 2022-09-04 21:23
2) 2022-09-04 21:23 - 2022-09-04 23:23
3) 2022-09-08 22:46 - 2022-09-11 01:30
4) 2022-09-11 01:30 - 2022-09-11 19:19
final detection: 2022-09-11 19:19

Deployment ID 42226 (first year F)

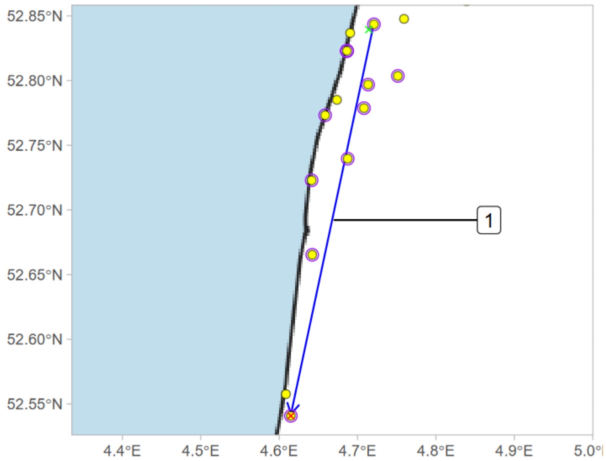


release: 2022-09-06 15:15
1) 2022-09-06 18:52 - 2022-09-06 20:31
final detection: 2022-09-06 20:32

Deployment ID 42227 (first year M)

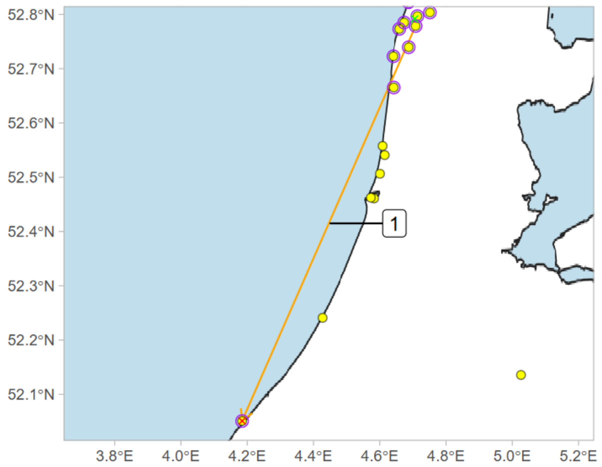


release: 2022-09-05 16:15
1) 2022-09-05 19:00 - 2022-09-08 19:45
final detection: 2022-09-08 20:42

Deployment ID 42228 (first year M)

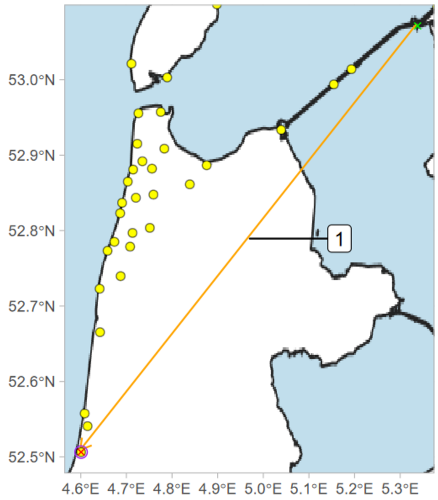


release: 2022-09-04 20:00
1) 2022-09-04 20:00 - 2022-09-06 21:58
final detection: 2022-09-07 20:20

Deployment ID 42232 (adult F)

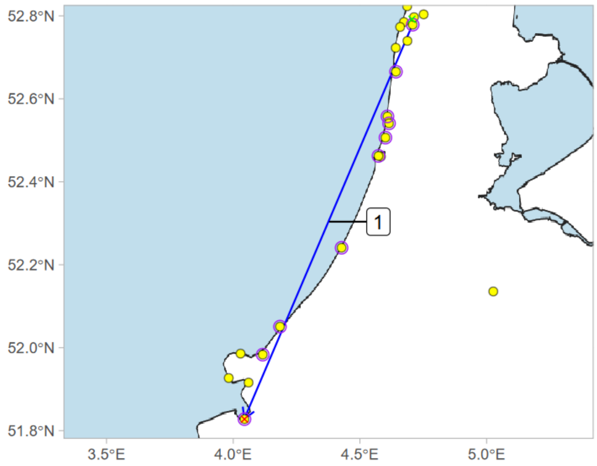


release: 2022-08-31 22:00
1) 2022-09-03 20:01 - 2022-09-04 01:22
final detection: 2022-09-04 02:02

Deployment ID 42235 (first year F)

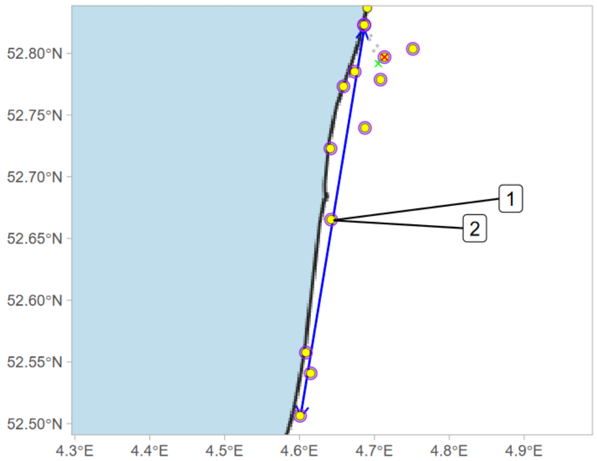


release: 2022-08-31 16:00
1) 2022-09-01 19:38 - 2022-09-01 21:18
2) 2022-09-01 21:50 - 2022-09-01 23:33
final detection: 2022-09-04 19:58

Deployment ID 42236 (first year M)

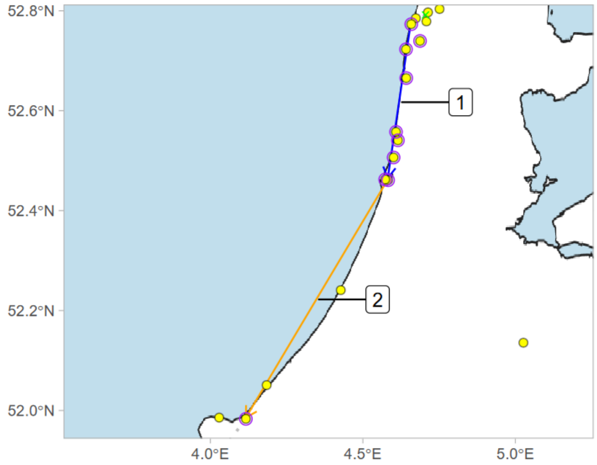


release: 2022-08-31 23:55
1) 2022-09-02 19:22 - 2022-09-02 21:25
2) 2022-09-02 21:25 - 2022-09-03 23:50
final detection: 2022-09-04 02:21

Deployment ID 42246 (adult F)

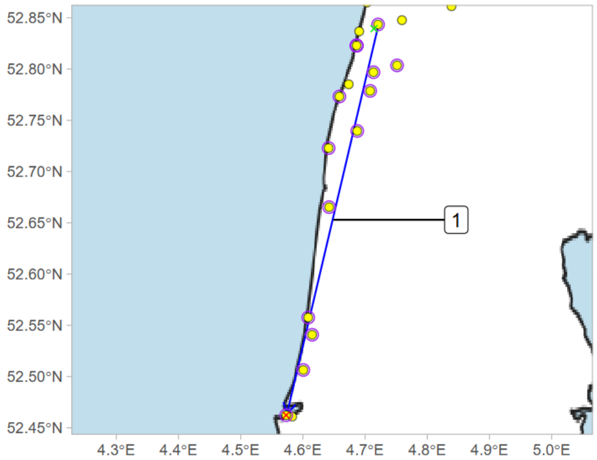


release: 2022-09-01 12:00
1) 2022-09-01 19:22 - 2022-09-01 21:18
final detection: 2022-09-01 21:20

Deployment ID 42247 (first year F)

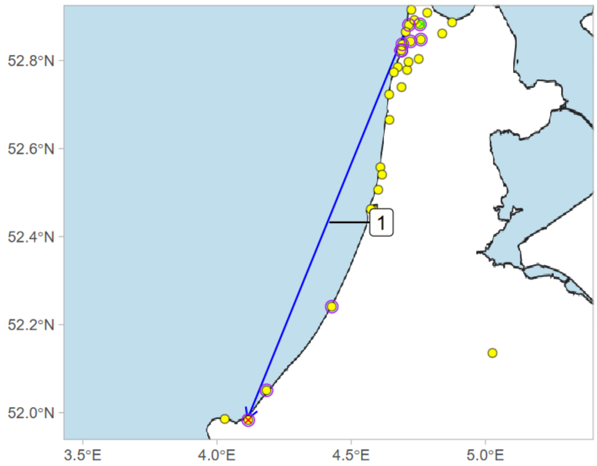


release: 2022-09-01 20:00
1) 2022-09-03 19:50 - 2022-09-04 01:36
final detection: 2022-09-04 01:37

Deployment ID 42248 (first year F)

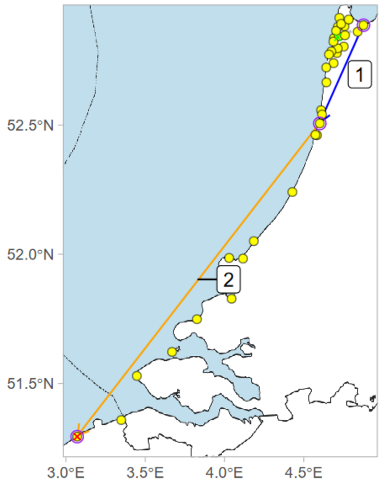


release: 2022-09-01 12:00
1) 2022-09-03 19:20 - 2022-09-03 21:10
2) 2022-09-03 21:10 - 2022-09-13 20:09
final detection: 2022-09-13 20:10

Deployment ID 42249 (adult F)

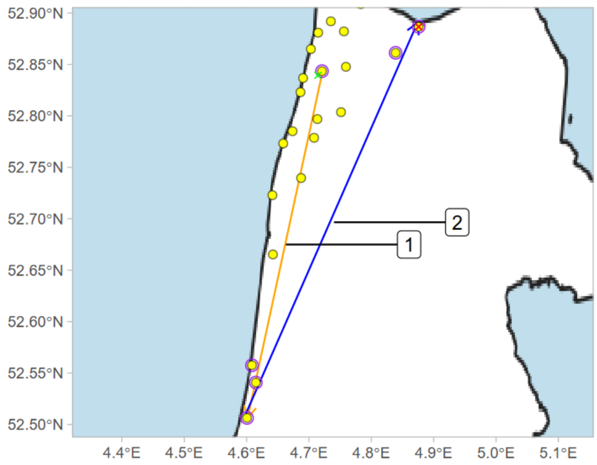


release: 2022-08-31 22:50
1) 2022-08-31 23:15 - 2022-09-01 20:30
2) 2022-09-01 21:01 - 2022-09-02 00:28
final detection: 2022-09-02 23:45

Deployment ID 42251 (first year M)

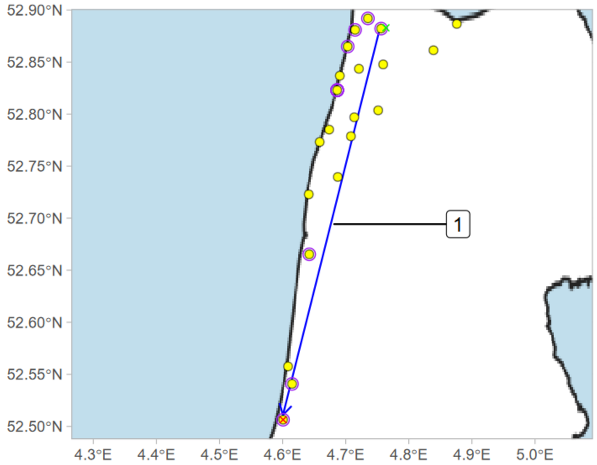


release: 2022-08-31 21:00
1) 2022-08-31 23:25 - 2022-09-01 03:06
final detection: 2022-09-01 03:13

Deployment ID 42252 (first year F)

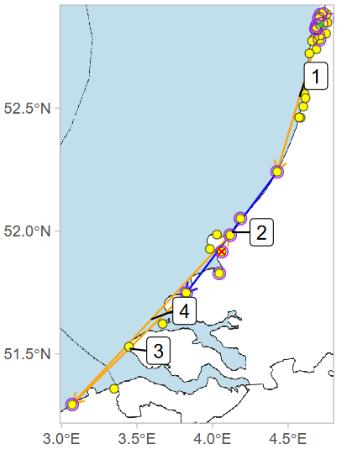


release: 2022-09-01 12:00
1) 2022-09-01 19:21 - 2022-09-03 22:21
2) 2022-09-03 22:21 - 2022-09-04 02:54
3) 2022-09-04 02:54 - 2022-09-05 23:45
4) 2022-09-05 23:45 - 2022-09-13 22:16
final detection: 2022-09-13 22:31

Deployment ID 42256 (adult F)

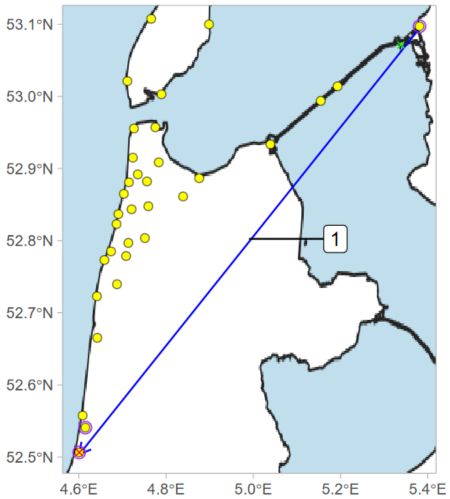


release: 2022-10-11 22:25
1) 2022-11-04 17:27 - 2022-11-04 22:58
final detection: 2022-11-04 23:00

Deployment ID 42259 (first year M)

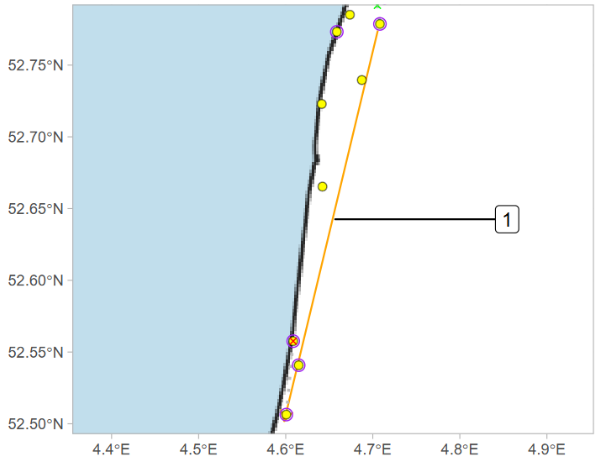


release: 2022-09-01 23:15
1) 2022-09-01 23:41 - 2022-09-02 21:22
final detection: 2022-09-02 21:24

Deployment ID 42260 (first year M)

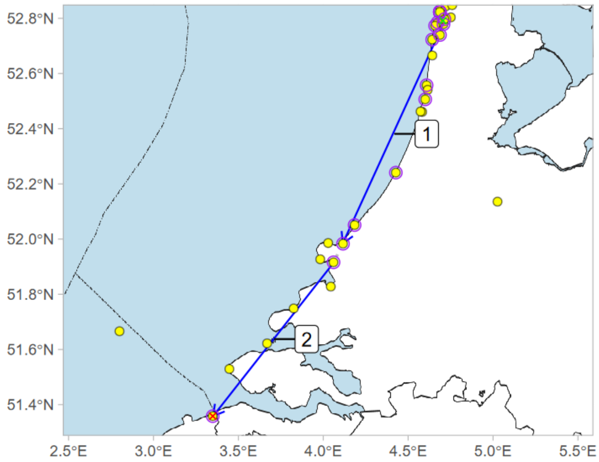


release: 2022-09-01 22:40
1) 2022-09-02 19:56 - 2022-09-03 00:26
2) 2022-09-03 19:05 - 2022-09-03 23:04
final detection: 2022-09-03 23:05

Deployment ID 42266 (first year M)

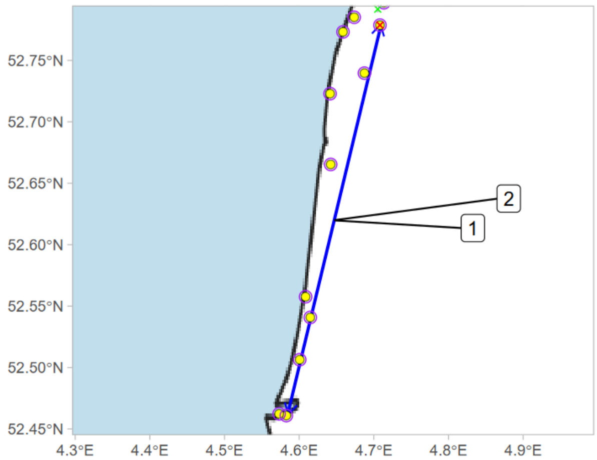


release: 2022-09-01 21:30
1) 2022-09-03 20:07 - 2022-09-03 21:52
2) 2022-09-03 21:52 - 2022-09-04 00:50
final detection: 2022-09-04 20:18

Deployment ID 42269 (adult F)

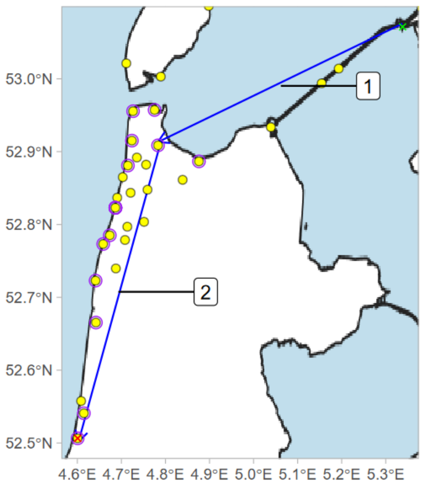


release: 2022-10-11 22:39
1) 2022-10-11 22:39 - 2022-10-12 00:24
2) 2022-10-12 17:45 - 2022-10-12 22:22
final detection: 2022-10-12 22:24

Deployment ID 42271 (first year F)

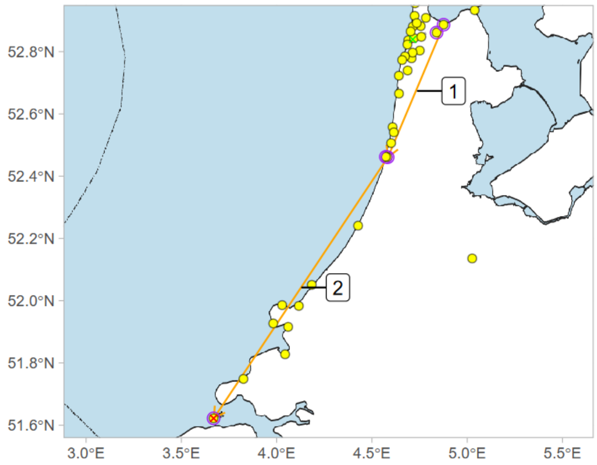


release: 2022-09-22 20:35
1) 2022-10-03 18:47 - 2022-10-16 20:12
2) 2022-10-16 20:12 - 2022-10-18 19:54
final detection: 2022-10-18 19:55

Deployment ID 42274 (first year F)

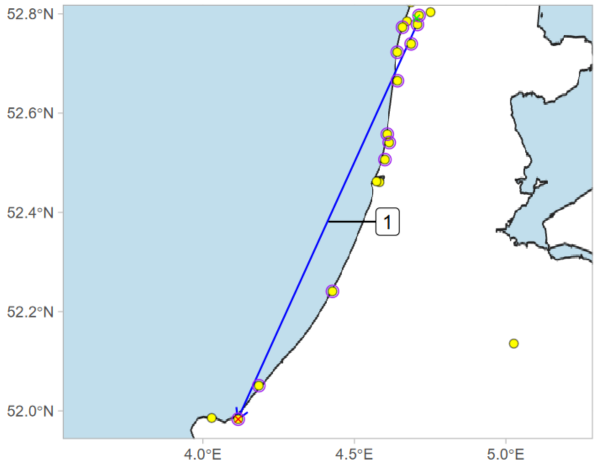


release: 2022-09-01 21:40
1) 2022-09-24 18:52 - 2022-09-24 20:26
final detection: 2022-09-24 20:26

Deployment ID 42276 (first year M)

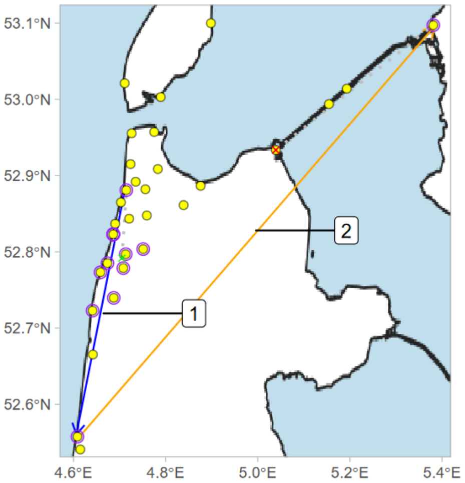


release: 2022-09-01 23:45
1) 2022-09-02 20:28 - 2022-09-02 23:23
2) 2022-09-02 23:23 - 2022-09-04 21:20
final detection: 2022-09-09 02:32

Deployment ID 42278 (first year F)

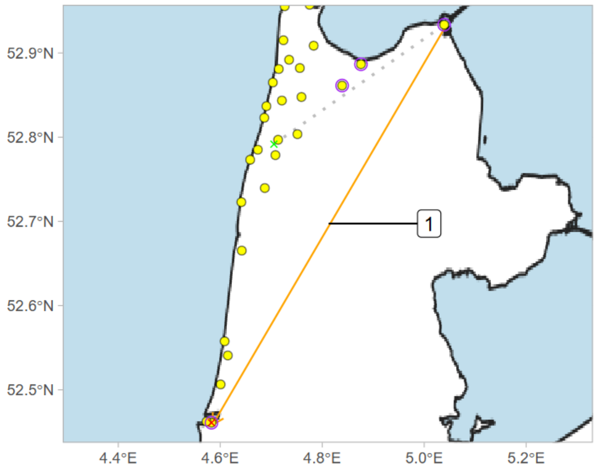


release: 2022-09-01 22:00
1) 2022-09-06 20:43 - 2022-09-08 00:08
final detection: 2022-09-08 00:09

Deployment ID 42290 (first year F)

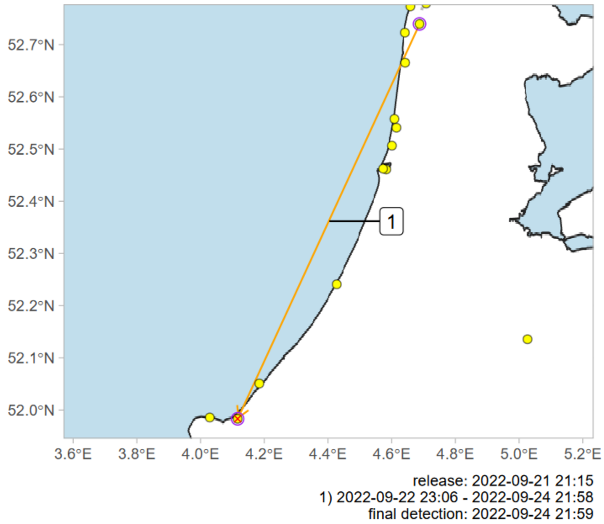


Deployment ID 42292 (first year F)

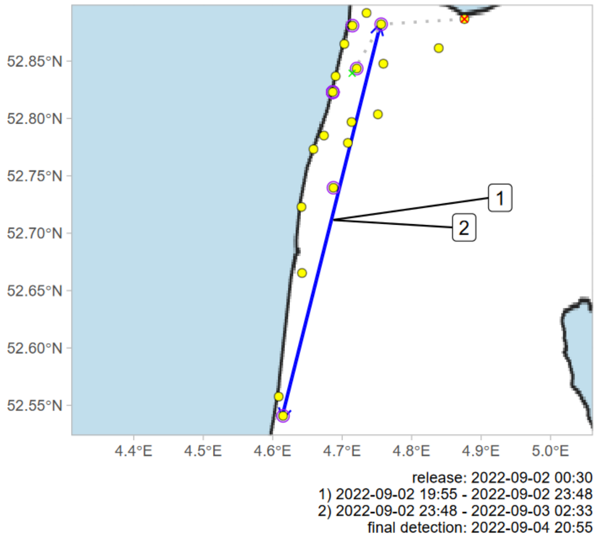


Deployment ID 42296 (first year M)

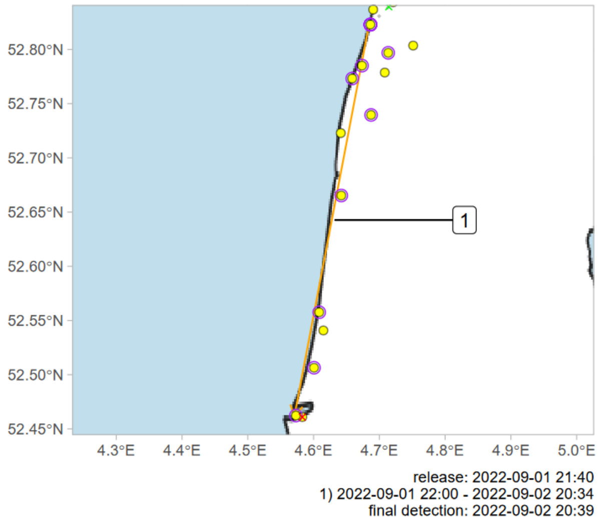


Deployment ID 42300 (first year M)

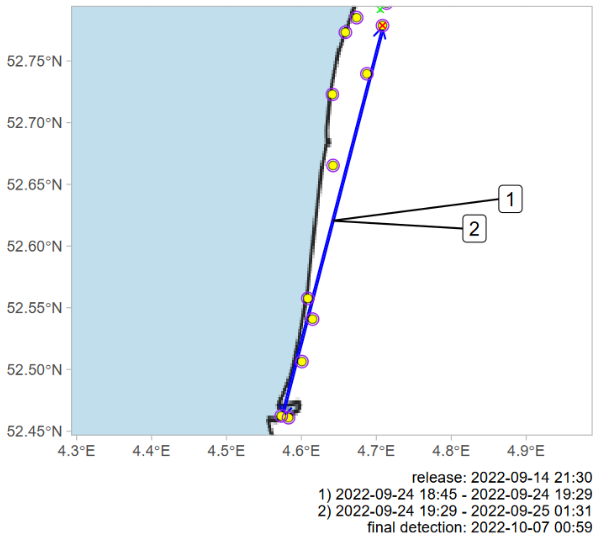


Deployment ID 42301 (first year M)

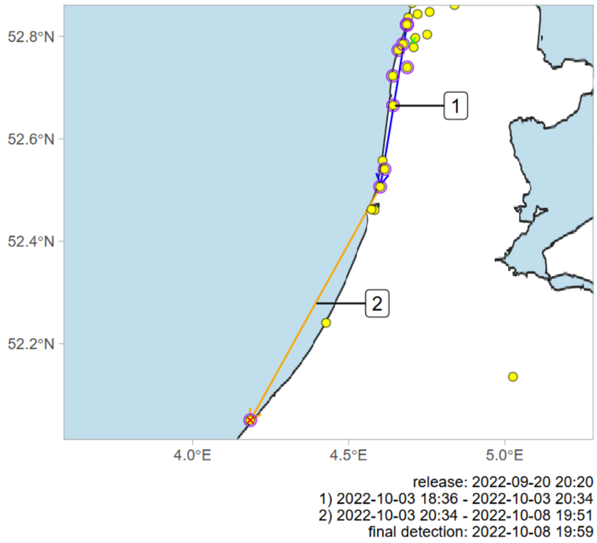


Deployment ID 42306 (adult M)

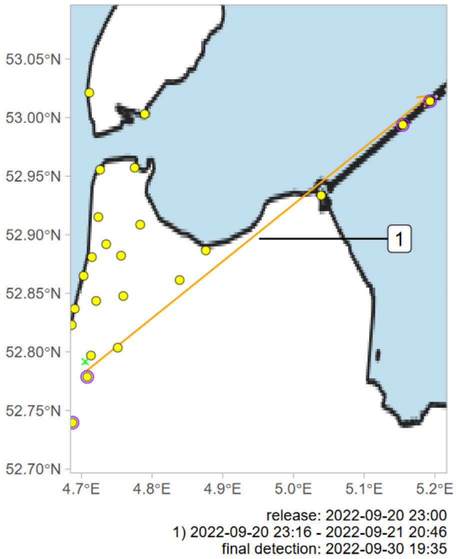

Deployment ID 42307 (first year M)

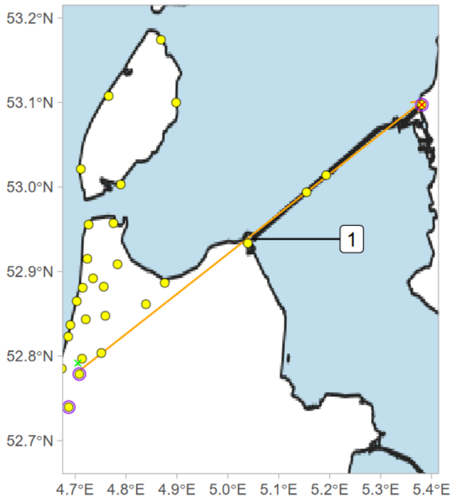


release: 2022-09-20 21:40
1) 2022-09-20 22:03 - 2022-09-21 20:20
final detection: 2022-10-09 17:50

Deployment ID 42315 (first year M)

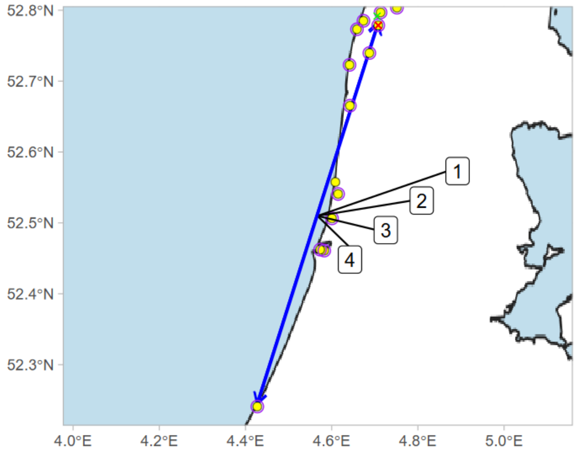


release: 2022-09-07 22:42
1) 2022-09-10 19:15 - 2022-09-10 22:40
2) 2022-09-10 22:40 - 2022-09-11 01:22
3) 2022-09-13 19:24 - 2022-09-13 21:34
4) 2022-09-13 21:34 - 2022-09-14 02:35
final detection: 2022-09-23 17:26

Deployment ID 42316 (first year F)

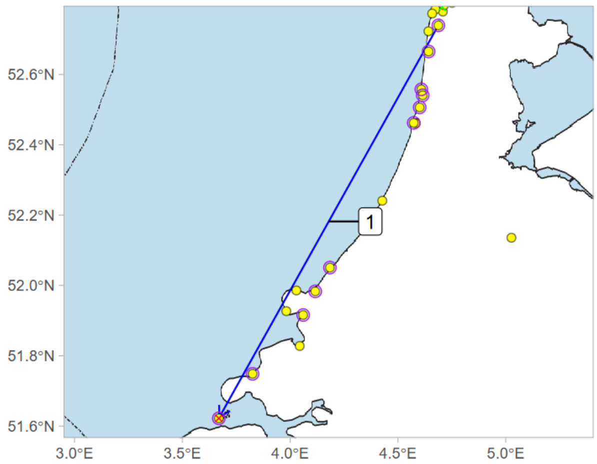


release: 2022-09-07 21:30
1) 2022-09-10 20:15 - 2022-09-11 01:11
final detection: 2022-09-11 01:12

Deployment ID 42327 (adult F)

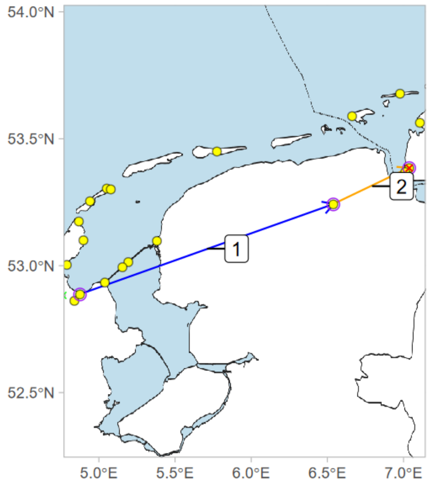


release: 2022-09-22 20:45
1) 2022-09-25 19:38 - 2022-09-25 23:27
2) 2022-09-25 23:27 - 2022-10-02 18:58
final detection: 2022-10-02 18:59

Deployment ID 42408 (adult F)

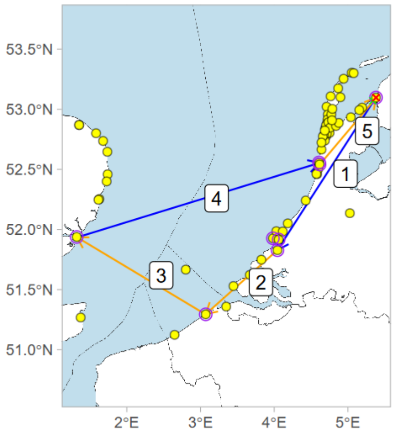


release: 2022-09-11 19:55
1) 2022-09-13 18:51 - 2022-09-14 00:34
2) 2022-09-14 00:34 - 2022-09-15 01:19
3) 2022-09-15 01:19 - 2022-10-03 19:09
4) 2022-10-03 19:09 - 2022-10-04 03:31
5) 2022-10-04 18:08 - 2022-10-06 17:56
final detection: 2022-10-11 17:41

Deployment ID 42412 (adult M)

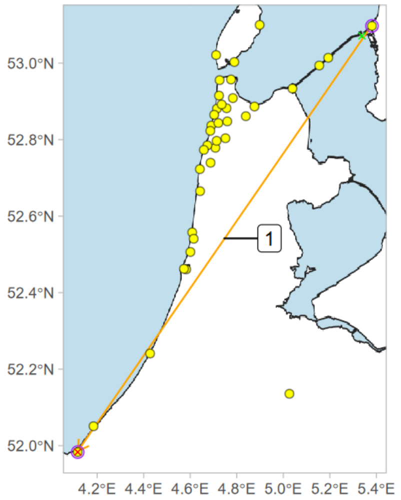


release: 2022-09-22 22:45
1) 2022-09-22 23:10 - 2022-09-24 21:37
final detection: 2022-09-24 21:38
