## Appendix S5 for "Stopover departure decisions and movement patterns of migratory bats during autumn migration"

### Appendix S5: output of the models

#### *Night-to-night departures*

| | mean | sd | 0.0250 | 0.5000 | 0.9750 | Rhat | n.eff | overlap0 | f |
|---|---|---|---|---|---|---|---|---|---|
| beta0 | 0.2290 | 0.4438 | -0.6922 | 0.2480 | 1.0358 | 1.0031 | 2410 | 1 | 0.712961 |
| sigma_ind | 1.0962 | 0.3009 | 0.5376 | 1.0824 | 1.7266 | 1.0041 | 1871 | 0 | 1 |
| beta_jdate_c | 0.0161 | 0.0141 | -0.0136 | 0.0170 | 0.0415 | 1.0009 | 9677 | 1 | 0.87515 |
| beta_jdate_c_sq | -0.0022 | 0.0005 | -0.0032 | -0.0021 | -0.0013 | 1.0040 | 1663 | 0 | 1 |
| beta_crosswind_200 | -0.0412 | 0.0306 | -0.1016 | -0.0409 | 0.0183 | 1.0008 | 7911 | 1 | 0.911472 |
| beta_crosswind_200_sq | 0.0262 | 0.0049 | 0.0169 | 0.0260 | 0.0363 | 1.0009 | 6966 | 0 | 1 |
| beta_tailwind_200 | -0.1739 | 0.0300 | -0.2357 | -0.1727 | -0.1182 | 1.0102 | 648 | 0 | 1 |
| beta_tailwind_200_sq | 0.0093 | 0.0033 | 0.0033 | 0.0092 | 0.0161 | 1.0079 | 841 | 0 | 0.999522 |
| beta_dryness | -1.1506 | 0.4091 | -1.9507 | -1.1499 | -0.3441 | 1.0013 | 4951 | 0 | 0.997394 |
| beta_cloud_by_dryness | 1.2625 | 0.6181 | 0.0707 | 1.2545 | 2.4983 | 1.0009 | 7426 | 0 | 0.981122 |
| beta_temp | -0.1188 | 0.0906 | -0.2998 | -0.1176 | 0.0559 | 1.0026 | 2500 | 1 | 0.907661 |
| beta_Male | 0.7474 | 0.4132 | -0.0071 | 0.7284 | 1.6154 | 1.0006 | 13476 | 1 | 0.9739 |
| beta_Adult | 0.5382 | 0.3589 | -0.1341 | 0.5268 | 1.2740 | 1.0018 | 3803 | 1 | 0.941289 |
| beta_Male_jdate_c | 0.0491 | 0.0226 | 0.0063 | 0.0485 | 0.0955 | 1.0017 | 3820 | 0 | 0.987933 |
| deviance | 455.5168 | 22.6779 | 411.1207 | 455.5765 | 500.1475 | 1.0025 | 2794 | 0 | 1 |

DIC = 711.663

#### *Within-night departures*

| | mean | sd | 0.0250 | 0.5000 | 0.9750 | Rhat | n.eff | overlap0 | f |
|---|---|---|---|---|---|---|---|---|---|
| beta0 | -3.2302 | 3.1440 | -9.7436 | -3.0670 | 2.5576 | 1.0047 | 1561 | 1 | 0.858422 |
| sigma_ind | 13.9426 | 4.0645 | 6.7736 | 13.6892 | 22.6126 | 1.0068 | 1166 | 0 | 1 |
| beta_time_since_sunset | -24.0768 | 6.5363 | -37.6485 | -23.7750 | -12.2344 | 1.0072 | 1104 | 0 | 1 |
| beta_crosswind_200 | 0.4877 | 0.4031 | -0.2289 | 0.4545 | 1.3784 | 1.0020 | 4489 | 1 | 0.906783 |
| beta_tailwind_200 | -0.7979 | 0.4061 | -1.6997 | -0.7584 | -0.1239 | 1.0022 | 3809 | 0 | 0.991033 |
| beta_dryness | 1.4130 | 3.7934 | -6.0642 | 1.3708 | 9.0674 | 1.0027 | 2570 | 1 | 0.648578 |
| beta_cloud_by_dryness | -0.9919 | 4.7340 | -10.4476 | -1.0070 | 8.4862 | 1.0018 | 3843 | 1 | 0.589211 |
| beta_moon_illumination | -3.2637 | 4.3505 | -12.0546 | -3.1433 | 5.1129 | 1.0041 | 1695 | 1 | 0.782272 |
| beta_hrly_temp | 0.4696 | 0.6614 | -0.7606 | 0.4378 | 1.8870 | 1.0026 | 2863 | 1 | 0.7732 |
| deviance | 29.3451 | 12.7512 | 10.4479 | 27.3782 | 59.6913 | 1.0033 | 2502 | 0 | 1 |

DIC = 110.2892

Both models successfully converged based on Rhat values (all < 1.01).

Rhat is the potential scale reduction factor (at convergence, Rhat=1).

For each parameter, n.eff is a crude measure of effective sample size.

overlap0 checks if 0 falls in the parameter's 95% credible interval.

f is the proportion of the posterior with the same sign as the mean;

i.e., our confidence that the parameter is positive or negative.
